# ERK1/2 inhibition disrupts alcohol memory reconsolidation and prevents relapse

**DOI:** 10.1101/2023.12.12.571297

**Authors:** Nofar Rahamim, Mirit Liran, Coral Aronovici, Hila Flumin, Tamar Gordon, Nataly Urshansky, Segev Barak

**Author notes:** Corresponding Author: Segev Barak, School of Psychological Sciences The Sagol School of Neuroscience Tel Aviv University Tel Aviv 69978, Israel.

## Abstract

Relapse to alcohol abuse after periods of abstinence, often caused by cue-induced alcohol craving, is a major challenge in the treatment of alcohol addiction. Therefore, disruption of the cue-alcohol associative memories can diminish the risk of relapse. Upon retrieval, memories become temporarily labile before they reconsolidate in a process that requires protein synthesis. Accumulating evidence suggests that the mammalian target of rapamycin complex 1 (mTORC1), which is responsible for the translation of a subset of dendritic proteins, is crucial for memory reconsolidation. Here, we explored the involvement of two regulatory pathways of mTORC1, namely phosphoinositide 3-kinase (PI3K)-AKT and extracellular regulated kinase1/2 (ERK1/2), in the reconsolidation process in a rat model of non-operant alcohol self-administration. We found that retrieval of alcohol memories using an odor-taste cue increased ERK1/2 activation in the amygdala, but did not affect the PI3K-AKT pathway. Importantly, inhibition of ERK1/2 shortly after alcohol memory retrieval impaired reconsolidation and led to long-lasting suppression of relapse to alcohol drinking. Additionally, we show that attenuation of alcohol memories and relapse was also induced by post-retrieval administration of lacosamide, an inhibitor of collapsin response mediator protein-2 (CRMP2) – a translational product of mTORC1 that is functionally regulated by PI3K-AKT signaling. Together, our findings provide evidence for the crucial role of ERK1/2 and CRMP2 in the reconsolidation of alcohol memories, and mark the FDA-approved drug, lacosamide, as a potential treatment for alcohol use disorder.

## 1. Introduction

Alcohol use disorder (AUD) is a chronic psychiatric disorder, with very limited treatments ^1,2^. As in other substance use disorders, relapse prevention is a main challenge in treating AUD ^2,3^. Relapse is often driven by cue-induced craving, caused by re-exposure to alcohol-associated cues ^4–8^. Thus, disruption of the memories for the cue-alcohol association is expected to suppress cue-induced relapse.

It is believed that upon their retrieval, memories become temporarily unstable during an active state and then undergo a process of re-stabilization, termed memory reconsolidation. Memories have been shown to be labile during their reconsolidation process and consequently can be manipulated ^9–12^. The therapeutic potential of interference with the reconsolidation process in attenuating or altering maladaptive associative memories has been increasingly acknowledged ^6,13–15^. Thus, it has been demonstrated that the inhibition of protein synthesis during memory reconsolidation (i.e., following memory retrieval), suppresses the behavioral expression of various types of memories ^16–22^ including prevention of alcohol relapse ^17,21^, suggesting that memory reconsolidation requires *de-novo* protein synthesis ^9,16,23^.

Mammalian target of rapamycin complex 1 (mTORC1), a kinase that is crucial for the translation of a subset of dendritic proteins, and implicated in synaptic plasticity and memory processes ^24–26^, has been shown to be crucial for alcohol memory reconsolidation ^17^. Specifically, The retrieval of alcohol memories in rats with a history of alcohol intake induced activation of mTORC1 signaling in the central amygdala (CeA), the medial prefrontal cortex (mPFC), and the orbitofrontal cortex (OFC), which led to increased expression of several mTORC1-regulated synaptic proteins ^17^. Administration of the mTORC1 inhibitor rapamycin following alcohol memory retrieval resulted in long-lasting suppression of relapse to alcohol consumption, without affecting memories for a natural reward (sucrose) ^17^. Furthermore, post-retrieval administration of rapamycin was also shown to produce a long-term decrease in the expression of alcohol-, morphine-, and cocaine-conditioned place preference ^21^.

The canonical mTORC1 activator is the phosphoinositide 3-kinase (PI3K)-AKT signaling pathway ^27–31^. When active, AKT phosphorylates, and thereby inactivates, the glycogen synthase kinase-3β (GSK3β) ^32^. Activation of AKT-GSK3β-mTORC1 signaling was observed during the reconsolidation of cocaine ^33–36^ and heroin memories ^37^. Moreover, inhibition of PI3K ^38^ or AKT-dependent activation of mTORC1 ^39^ was shown to impair fear memory reconsolidation. Together, these studies further imply the involvement of PI3K-AKT-mTORC1 signaling in the reconsolidation process.

Another positive regulator of mTORC1 is the extracellular regulated kinase 1/2, which is part of the mitogen-activated protein kinase (MAPK) signaling pathway ^27,28,40^. Upregulation of ERK1/2 was observed following cue-induced relapse to alcohol-seeking ^41,42^, whereas, inhibition of ERK1/2 impaired the reconsolidation of cocaine and morphine memories ^43–46^. This suggests that ERK1/2 activation is also required for the reconsolidation of drug-associated memories.

The PI3K-AKT and the ERK1/2 signaling pathways were both shown to be involved in alcohol consumption and relapse. Thus, PI3K-AKT signaling is thought to underlie the mTORC1 activity that contributes to the escalation and maintenance of alcohol-drinking phenotypes ^27–31^. In addition, the GSK3β substrate and mTORC1 downstream product, collapsin response mediator protein 2 (CRMP2) ^47^, which promotes microtubule assembly and neurite outgrowth ^48–50^, was previously implicated in relapse to alcohol seeking ^51^. ERK1/2 signaling has primarily been characterized as a mechanism that gates the level of alcohol intake ^29,52–54^. Nonetheless, besides mTORC1, the role of these signaling pathways in the reconsolidation of alcohol memories has not been directedly investigated.

Here, we characterized the involvement of the PI3K-AKT and ERK1/2 signaling pathways in alcohol memory reconsolidation. To this end, we used the odor-taste cue retrieval method, previously shown to elicit mTORC1 signaling ^17^. Thus, we examined the effect of alcohol memory retrieval on the activation of several effectors of these pathways, focusing on the mPFC the amygdala, and the NAc, brain regions previously implicated in the reconsolidation of drug memories ^17,33,35,45,46,55,56^. To establish causality, we tested whether inhibition of certain elements in these signaling pathways, namely CRMP2, GSK3β, and ERK1/2, can impair the reconsolidation process and prevent relapse to alcohol drinking.

## 2. Materials and methods

### 2.1. Animals

Male and female Wistar rats (160-240 gr at the beginning of experiments 1-4, and 220-390 gr at the beginning of experiment 5) were bred at the Tel-Aviv University animal facility. Animals were individually housed under a 12-h light-dark cycle (lights on at 7 a.m.) with food and water available ad libitum. Animals were weighed once a week to control for weight loss. All experimental protocols were approved by and conformed to the guidelines of the Institutional Animal Care and Use Committee of Tel Aviv University, and the NIH guidelines (animal welfare assurance number A5010-01). All efforts were made to minimize the number of animals used.

### 2.2. Drugs and Reagents

The alcohol-drinking solution was prepared by diluting ethanol absolute (0005250502000) (BioLab, Jerusalem, Israel) to a 20% (v/v) solution with tap water. Isoflurane was obtained from Piramal Critical Care (Bethlehem, PA, USA). Rabbit antibodies for pAKT-Thr308 (9275), pAKT-Ser473 (4046), total AKT (9272), pGSK3β-Ser9 (9323), total pGSK3β (9315), pCRMP2-Thr514 (9397), total CRMP2 (9393), pERK1/2-Thr202/Tyr204 (4370), total ERK1/2 (9102), pS6-Ser235/236 (2211), total S6 (2217), as well as anti-rabbit (7074) and anti-mouse (7076) horseradish peroxidase (HRP)-linked antibodies, were obtained from Cell Signaling Technology (Denvers, MA, USA). A mouse antibody for GAPDH (sc-32233) was obtained from Santa Cruz Biotechnology (Santa Cruz, CA, USA). A protease/phosphatase inhibitor cocktail (#5872) was purchased from Cell Signaling Technology (Danvers, MA, USA). Pierce BCA protein assay kit (23227) and SuperSignal chemiluminescent substrate (34578) were purchased from Thermo Fisher Scientific (Waltham, MA, USA). Immobilon-P PVDF membrane (IPVH00005) was purchased from Merck (Rehovot, Israel). BLUeye prestained protein ladder (PM007-0500) was purchased from BIO-HELIX (New Taipei City, Taiwan). 12% Criterion TGX precast midi protein gels (5671044) and 4x Laemmli sample buffer (1610747) were purchased from Bio-Rad Laboratories (Rishon Le Zion, Israel). The CRMP2 inhibitor, lacosamide (465325000) (Thermo Scientific Chemicals, Waltham, MA, USA), was solubilized in saline. The GSK3β inhibitor, SB 216763 (HY-12012) (MedChemExpress, Monmouth Junction, NJ, USA), was solubilized in DMSO. The MEK1/2 inhibitor, SL-327 (S1066) (Selleck Chemicals, Houston, TX, USA), was solubilized in DMSO.

### 2.3. Behavioral Procedure

#### Intermittent access to 20% alcohol in 2-bottle choice (IA2BC)

After one week of habituation to individual cages, mice and rats were trained to consume 20% alcohol solution in the Intermittent Access 2-bottle-choice procedure (IA2BC) as previously described ^57–59^. Animals were given three 24-hour sessions of free access to 2-bottle choice per week (tap water and 20% alcohol v/v) on Sundays, Tuesdays, and Thursdays. Alcohol drinking sessions were followed by a 24- or 48-hour deprivation period, in which the animals received only water, producing repeated cycles of intoxication and withdrawal. The position (left or right) of the alcohol and water bottles was alternated between the sessions to control for side preference. Alcohol exposure in this protocol was shown to produce high levels of alcohol intake and blood alcohol concentration, especially in the first hours of drinking ^57–61^. Water and alcohol bottles were weighed before and after each alcohol-drinking session, and consumption levels were normalized to body weight. In the IA2BC experiments that included injections (experiments 2-4), the rats were handled weekly and received several sessions of injection handling with saline the week before the injection of the pharmacological substance.

##### Memory retrieval after IA2BC

After 9-10 weeks of training, rats were subjected to 10 days of abstinence. On the 11^th^ day of abstinence, alcohol memory retrieval was done in the rats’ home cages with 10 min exposure to an empty bottle of alcohol with its tip covered with a drop of alcohol serving as an odor-taste cue, as was previously described^17^. Control rats were left untouched in their home cages.

### 2.4. Western blot

Western blot procedure was conducted as previously described ^62^. Briefly, immediately after dissection, brain tissues were homogenized in a radioimmunoprecipitation assay (RIPA) buffer containing:150 mM NaCl, 5 mM EDTA, 50 mM Tris HCl, 1% Triton X-100, 0.5% sodium deoxycholate, 0.1% sodium dodecyl sulfate (SDS) and protease and phosphatase inhibitors. The homogenates were then centrifuged at 14,600 rpm and the supernatants were collected and stored at −80°C until use. Prior to sample preparations, protein concentrations were determined using BCA assay. An equal amount of each sample (30 μg) was mixed with 4x Laemmli sample buffer that was added with dithiothreitol (DTT) (50 mM), and the mixture was boiled for five min at 100°C. Samples were separated on a 12% polyacrylamide gel and were then transferred onto a PVDF membrane at 100 V for one hour. Membranes were blocked for one hour at room temperature with 5% bovine serum albumin (BSA) in TBST Tris-buffered saline and 0.1% Tween 20 (TBST), and were then incubated overnight at 4°C with the appropriate primary antibody (pAKT-The308 1:500, pAKT-Ser473 1:2000, pGSK3β 1:1000, pCRMP2 1:800, pERK1/2 1:500, pS6 1:1000, GAPDH 1:10,000). Membranes were washed with TBST and probed with the appropriate HRP-conjugated secondary antibodies (anti-rabbit 1:3000, anti-mouse 1:5000) for one hour at room temperature. After washing with TBST, the bound antibodies were visualized using enhanced chemiluminescent (ECL) HRP substrate and captured with the VILBER Fusion FX imaging system. Membranes were then incubated for one hour in a stripping buffer containing 62.5 mM Tris, 100 mM β-mercaptoethanol, and 2% SDS, which was heated to 50°C. The membranes were then washed extensively with double distilled water (DDW) and with TBST, and were re-incubated overnight at 4°C with the appropriate primary antibody (total AKT 1:1000, total GSK3β 1:1000, total CRMP2 1:1000, total ERK1/2 1:1000, total S6 1:1000). Membranes were again washed, incubated with the anti-rabbit secondary antibody, and visualized. Band intensities were quantified using the ImageJ software (NIH), version 1.53k (see Supplementary File for images of the full-length bands). The optical density values of the phosphorylated proteins’ immunoreactivity were normalized to those of the total proteins, and immunoreactivity values of total CRMP2 were normalized to those of GAPDH. The phosphorylated/total protein expression of the experimental groups was calculated as a percentage of the control group.

### 2.5. Experimental design and statistical analysis

The allocation of rats to the experimental groups was done based on their average alcohol drinking, to create groups with approximately equal alcohol drinking levels. Sex was distributed approximately equally across the experiments and was initially analyzed as a factor. The analyses in all the experiments did not yield an interaction between sex and other factors. Therefore, data were collapsed across this factor.

#### 2.5.1. Experiment 1 – effects of alcohol memory retrieval on the activation of the PI3K-AKT and ERK1/2 signaling pathways

Rats were trained to voluntarily consume alcohol in the IA2BC protocol, as described above. After nine weeks of drinking, the rats were subjected to 10 days of abstinence. On the 11th day of the abstinence period, the rats’ alcohol memory was retrieved, using an odor-taste cue, as described above. Control rats did not undergo the memory retrieval procedure and were left untouched. mPFC, NAc, and amygdala tissues were collected 30 and 60 min after the alcohol memory retrieval procedure and were processed for a western-blot analysis. The samples were examined for changes in the phosphorylation levels of AKT, GSK3β, CRMP2, and ERK1/2, as well as changes in the total protein levels of CRMP2. The expression level for each target in each brain region was normalized to the no-retrieval group. The expression data of each target were analyzed by one-way ANOVA with a between factor of Group (no-retrieval, retrieval-30 min, retrieval-60 min).

#### 2.5.2. Experiment 2 – Effects of CRMP2 inhibition on alcohol memory reconsolidation

Rats were trained in the IA2BC protocol for nine weeks, followed by an abstinence period and a procedure of alcohol memory retrieval, as described in Experiment 1. Immediately after the memory retrieval, the rats were injected with the CRMP2 inhibitor, lacosamide (20 mg/kg, i.p., injection volume 1 ml/kg), or with a vehicle solution. The rats’ drinking behavior was tested 24 hours later. After the drinking test, the rats were subjected to two additional weeks of drinking training, followed by 10 days of abstinence. On the 11th day of abstinence, the rats were treated again with lacosamide or with vehicle, but without a prior procedure of alcohol memory retrieval. The treatment was done in a counterbalanced manner, i.e., rats that received lacosamide in the post-retrieval tests, received vehicle in the no-retrieval test and *vice versa*. Three rats that displayed drinking levels lower than 2 gr/kg (according to the average of the last three sessions prior to the abstinence) did not proceed to the no-retrieval test. In each test the following parameters were collected: alcohol consumption (gr/kg), alcohol preference (volume of alcohol consumed / volume of alcohol+water consumed), and water consumption (ml/kg). The data of each parameter in each test were analyzed by an unpaired t-test (vehicle vs. lacosamide).

#### 2.5.3. Experiment 3 – Effects of GSK3β inhibition on alcohol memory reconsolidation

Rats were trained in the IA2BC protocol for 10 weeks, followed by an abstinence period and a procedure of alcohol memory retrieval, as described in Experiment 1. Immediately after the memory retrieval, the rats were injected with the GSK3β inhibitor, SB 216763 (20 mg/kg, i.p., injection volume 0.5 ml/kg), or with a vehicle solution. The rats’ drinking behavior was tested 24 hours later. The parameters that were collected for the analysis were the same as in Experiment 2. The data of each parameter were analyzed by an unpaired t-test (vehicle vs. SB 216763).

#### 2.5.4. Experiment 4 – Effects of ERK1/2 inhibition on alcohol memory reconsolidation

Rats were trained in the IA2BC protocol for 10 weeks, followed by an abstinence period and a procedure of alcohol memory retrieval, as described in Experiment 1. Immediately after the memory retrieval, the rats were injected with the inhibitor of ERK1/2 upstream regulator MEK1/2, SL-327 (50 mg/kg, i.p., injection volume 0.6 ml/kg), or with a vehicle solution. The rats’ drinking behavior was tested 24 hours later and 14 days later. Between the two tests, the rats were abstinent from alcohol. After the second drinking test, the rats were subjected to three additional weeks of drinking training, followed by 10 days of abstinence, and an additional no-retrieval drinking test, as was done in Experiment 2. The parameters that were collected for the analyses were the same as in Experiment 2. The data of each parameter in each test was analyzed by an unpaired t-test (vehicle vs. SL-327). Rats were then subjected again to three additional weeks of drinking, followed by an abstinence period. On the 11th day of the abstinence period, the rats’ alcohol memory was retrieved, and amygdala tissues were collected two hours later and were processed for western blot analysis. Control rats did not undergo the memory retrieval procedure and were left untouched. The samples were examined for changes in the phosphorylation levels of ERK1/2. The phosphorylation levels of the retrieval group were normalized to the no-retrieval group. Data were analyzed by an unpaired t-test (no-retrieval vs. retrieval).

#### 2.5.4. Experiment 5 – Effects of systemic administration of the MEK1/2 inhibitor on the phosphorylation of ERK1/2 and S6 in the amygdala

Naïve rats were separated into individual cages and after one week, during which they were handled for injections, the rats were injected with SL-327 (50 mg/kg. i.p., injection volume 0.6 ml/kg). Control rats were injected with a vehicle solution. Amygdala tissues were collected 90 min afterward ^63,64^ and were processed for western blot analysis. The samples were examined for changes in the phosphorylation levels of ERK1/2 and S6. The phosphorylation levels of the SL-327 group were normalized to the vehicle group. Data were analyzed by an unpaired t-test (vehicle vs. SL-327).

## 3. Results

### Alcohol memory retrieval does not lead to activation of the PI3K-AKT signaling pathway

First, we examined whether the retrieval of alcohol memories activates the AKT-PI3K signaling pathway. To this end, rats were trained to consume alcohol in the IA2BC procedure for nine weeks. The average alcohol intake during the last five drinking sessions was 4.22 gr/kg (SD=1.65). Rats were then subjected to 10 days of abstinence. On the 11^th^ day of abstinence, alcohol memories were retrieved via the presentation of an odor-taste cue (see Methods). We then determined the phosphorylation levels of AKT, GSK3β, and CRMP2 in the mPFC, NAc, and amygdala, 30 and 60 min after memory retrieval. Since CRMP2 is also a translational product of mTORC1 activation ^65^, we also examined changes in the total protein levels of CRMP2 (see Experiment 1 in Methods).

As shown in Figure 1, retrieval of alcohol memory did not induce any changes in the phosphorylation levels of AKT, GSK3β and CRMP2 (one-way ANOVA: mPFC: pAKT (Thr308) [F(2,29)=0.07, p=0.935), pAKT (Ser473) [F(2,28)=0.14, p=0.874], pGSK3β [F(2,28)=0.66, p=0.523], pCRMP2 [F(2,26)=0.03, p=0.975]; NAc: pAKT (Thr308) [F(2,28)=0.68, p=0.514], pAKT (Ser473) [F(2,28)=1.28, p=0.293], pGSK3β [F(2,27)=0.24, p=0.791], pCRMP2: [F(2,28)=0.6, p=0.558]; Amygdala: pAKT (Thr308) [F(2,27)=0.04, p=0.961], pAKT (Ser473) [F(2,29)=0.23, p=0.793], pGSK3β [F(2,28)=0.19, p=0.831], pCRMP2 [F(2,28)=0.39, p=684]) (Figure S1).

**Figure 1.**
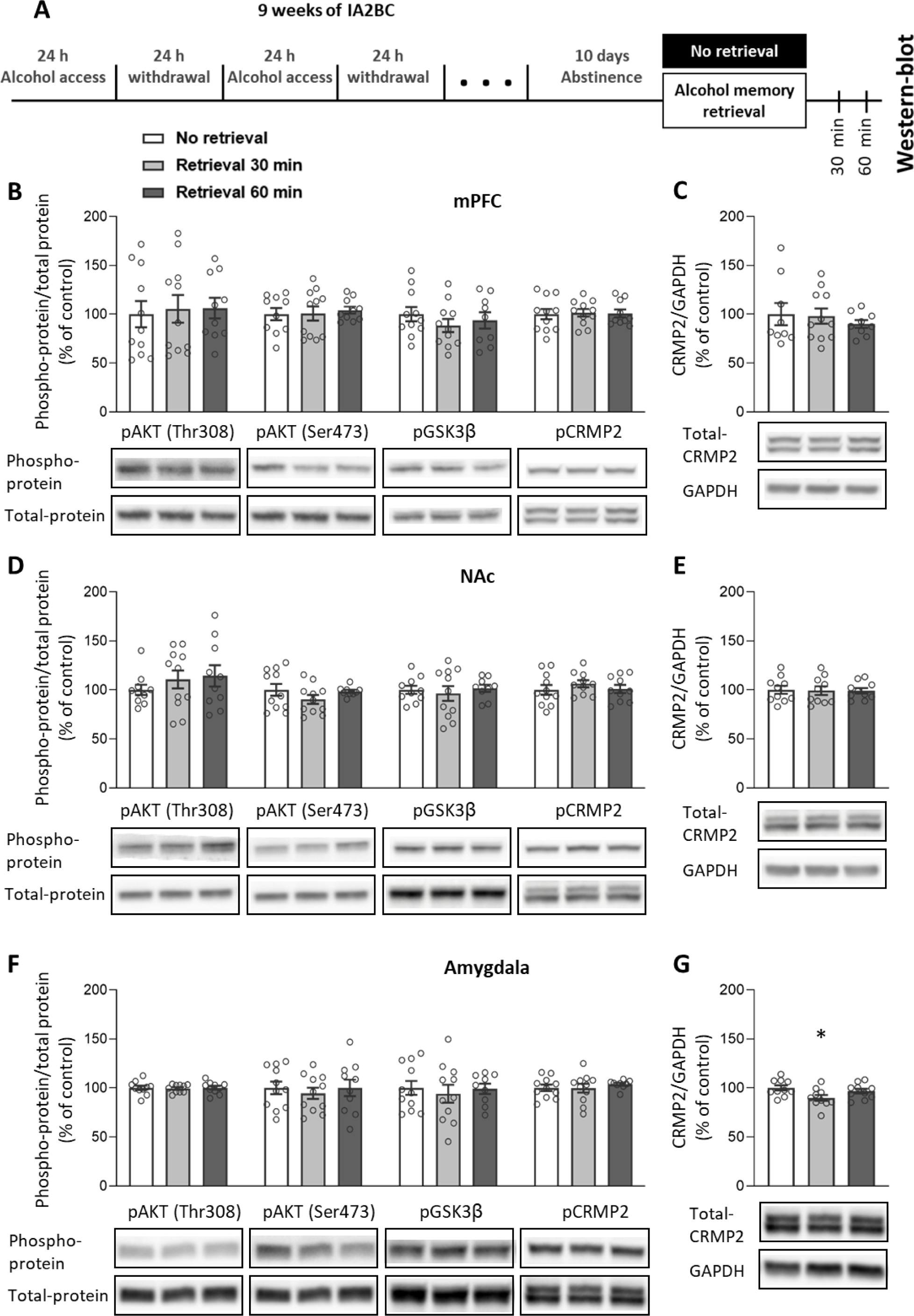
Alcohol memory retrieval has no effect on the activation of the PI3K-AKT signaling pathway. **A.** Experimental design and timeline. Rats consumed alcohol in the intermittent access to 20% alcohol in a 2-bottle choice paradigm for nine weeks, followed by 10 days of abstinence. After the abstinence period, alcohol memory was retrieved using an odor-taste cue. Tissues were collected 30 min and 60 min after the retrieval. Phosphorylation and total protein levels were determined by western blot. **B-G.** Phospho-protein levels of AKT (Thr308, Ser473), GSK3β (Ser9), and CRMP2 (Thr514) were normalized to the total protein immunoreactivity (B, D, F). Total protein levels of CRMP2 were normalized to GAPDH (C, E, G). Bar graphs represent mean ±S.E.M. of the percent of change from the control No retrieval group. n=9-11. * p<0.05 (No retrieval vs. Retrieval 30 min).

In the amygdala, a mild reduction in the expression of total-CRMP2 was observed 30 min after the alcohol memory retrieval (one-way ANOVA: F(2,7)=3.67, p=0.019. Dunnett post hoc: No-retrieval vs. retrieval 30-min, p=0.024). CRMP2 levels in the mPFC and the NAc were not affected by the retrieval procedure (one-way ANOVA: mPFC [F(2,26)=0.39, p=0.679]; NAc [F(2,27)=0.02, p=0.982]). These findings suggest that the retrieval of alcohol memory and the process of reconsolidation of alcohol memories do not involve PI3K-AKT-GSK3β signaling.

### CRMP2 inhibition following alcohol memory retrieval reduces subsequent alcohol consumption

The western blot analysis described above did not provide evidence for alteration of CRMP2 activation following alcohol memory retrieval. Nonetheless, elevated levels of CRMP2 were previously observed following the reinstatement of alcohol-seeking induced by alcohol-priming, while CRMP2 inhibition reduced reinstatement ^51^. Therefore, we next investigated whether inhibition of CRMP2 would affect alcohol memory reconsolidation, resulting in reduced relapse to alcohol drinking. To this end, we used lacosamide, an FDA-approved antiepileptic drug ^66^, which inhibits CRMP2’s ability to promote tubulin polymerization, and thereby impairs neurite outgrowth and branching^67,68^.

Rats were trained to consume alcohol in the IA2BC procedure for 11 weeks and were then subjected to 10 days of abstinence. On the next day, alcohol memory retrieval was conducted. Immediately afterward, the rats were injected with lacosamide (20 mg/kg, i.p., dose based on Liu et al. ^69^) or with vehicle. The rats’ drinking behavior was tested 24 hours later (see Experiment 2 in Methods).

As shown in Figure 2B-C, rats treated with lacosamide following memory retrieval consumed less alcohol and showed a lower preference for alcohol, compared with vehicle-treated controls (t-test: alcohol intake [t(19)=5.50, p<0.0001]; alcohol preference [t(19)=2.31, p=0.032]).

**Figure 2.**
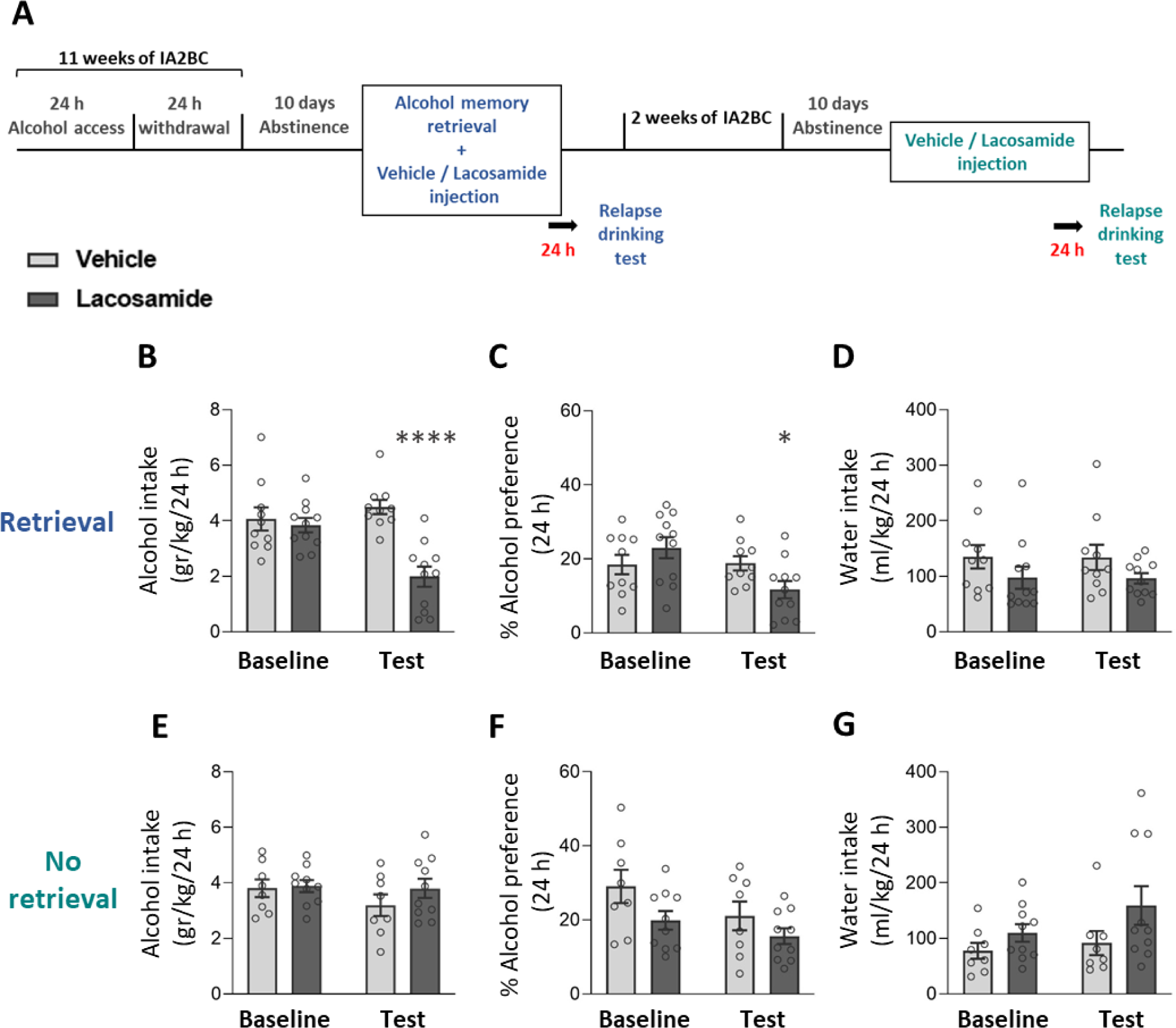
Lacosamide treatment after alcohol memory retrieval leads to reduced relapse to alcohol consumption and preference. **A.** Experimental design and timeline. Rats consumed alcohol in the intermittent access to 20% alcohol 2-bottle choice paradigm for 11 weeks, followed by 10 days of abstinence. After the abstinence period, alcohol memory was retrieved using an odor-taste cue. Immediately afterward, rats were injected with lacosamide (20 mg/kg) or vehicle. Relapse to alcohol drinking was tested 24 h later. After a 2-week re-training, the rats were subjected again to 10 days of abstinence and were then given an additional treatment of lacosamide or vehicle, without a preceding alcohol memory retrieval, and relapse was tested 24 h later. **B-G.** Alcohol intake (B, E), alcohol preference (C, F), and water intake (D, G) in a baseline and relapse test conducted a day after lacosamide treatment given following alcohol memory retrieval (B-D) or without memory retrieval (E-G). Baseline levels of alcohol/water intake/preference represent the average of the last five (B-D) or three sessions (E-G) prior to abstinence. Bar graphs represent mean ±S.E.M. of alcohol intake/preference. n=8-11. * p<0.05, **** p<0.0001 (Vehicle vs. Lacosamide).

To validate that the effect of lacosamide on relapse to alcohol drinking was due to an impaired reconsolidation process, we tested the effects of lacosamide without prior memory retrieval. To this end, we re-trained the rats in the IA2BC for two additional weeks. We then repeated the experiment as above, except that lacosamide was injected without prior memory retrieval, and a relapse to alcohol drinking test was held on the next day.

As shown in Figure 2E-F, lacosamide treatment without prior alcohol memory retrieval had no effect on the rats’ relapse to alcohol drinking or preference (t-test: alcohol intake [t(16)=1.16, p=0.263]; alcohol preference [t(16)=1.3, p=0.213]). Importantly, lacosamide treatment had no effect on water intake in both tests, regardless of alcohol memory retrieval (t-test: post-retrieval test [t(19)=1.58, p=0.131]; no-retrieval test t(16)=1.55, p=0.141]) (Figure 2D,G).

These results suggest that inhibition of CRMP2 after alcohol memory retrieval impairs the reconsolidation of alcohol memories, and thereby attenuates post-abstinence relapse to alcohol consumption.

### GSK3β inhibition following alcohol memory retrieval does not affect relapse to alcohol consumption

The ability of CRMP2 to bind to tubulin is negatively regulated by GSK3β ^47,70^. GSK3β, which was previously implicated in the reconsolidation of heroin-^37^ and cocaine-^33–36^ memories, also acts as a negative regulator of mTORC1 ^28^. Additionally, its activity was suggested to contribute to memory consolidation and reconsolidation by affecting the balance between N-methyl-D-aspartate (NMDA) receptor-dependent long-term potentiation (LTP) and depression (LTD) ^71,72^. Since we found that administration of the CRMP2 inhibitor, lacosamide, impairs the reconsolidation of alcohol memory, we next examined whether the inhibition of GSK3β following alcohol memory retrieval will also affect alcohol memory reconsolidation and relapse.

Rats were trained as described above and were injected with the GSK3β inhibitor SB 216763 (2.5 mg/kg, i.p., dose based on Shi et al. ^33^) or vehicle immediately after memory retrieval, followed by a relapse test 24 hours later (see Experiment 3 in Methods).

We found that SB 216763 had no effects on alcohol or water consumption or alcohol preference (t-test: alcohol intake [t(22)=0.29, p=0.775]; alcohol preference [t(21)=0.12, p=0.902]; water intake [t(21)=0.66, p=0.514]) (Figure 3). These results indicate that inhibition of GSK3β does not affect alcohol memory reconsolidation.

**Figure 3.**
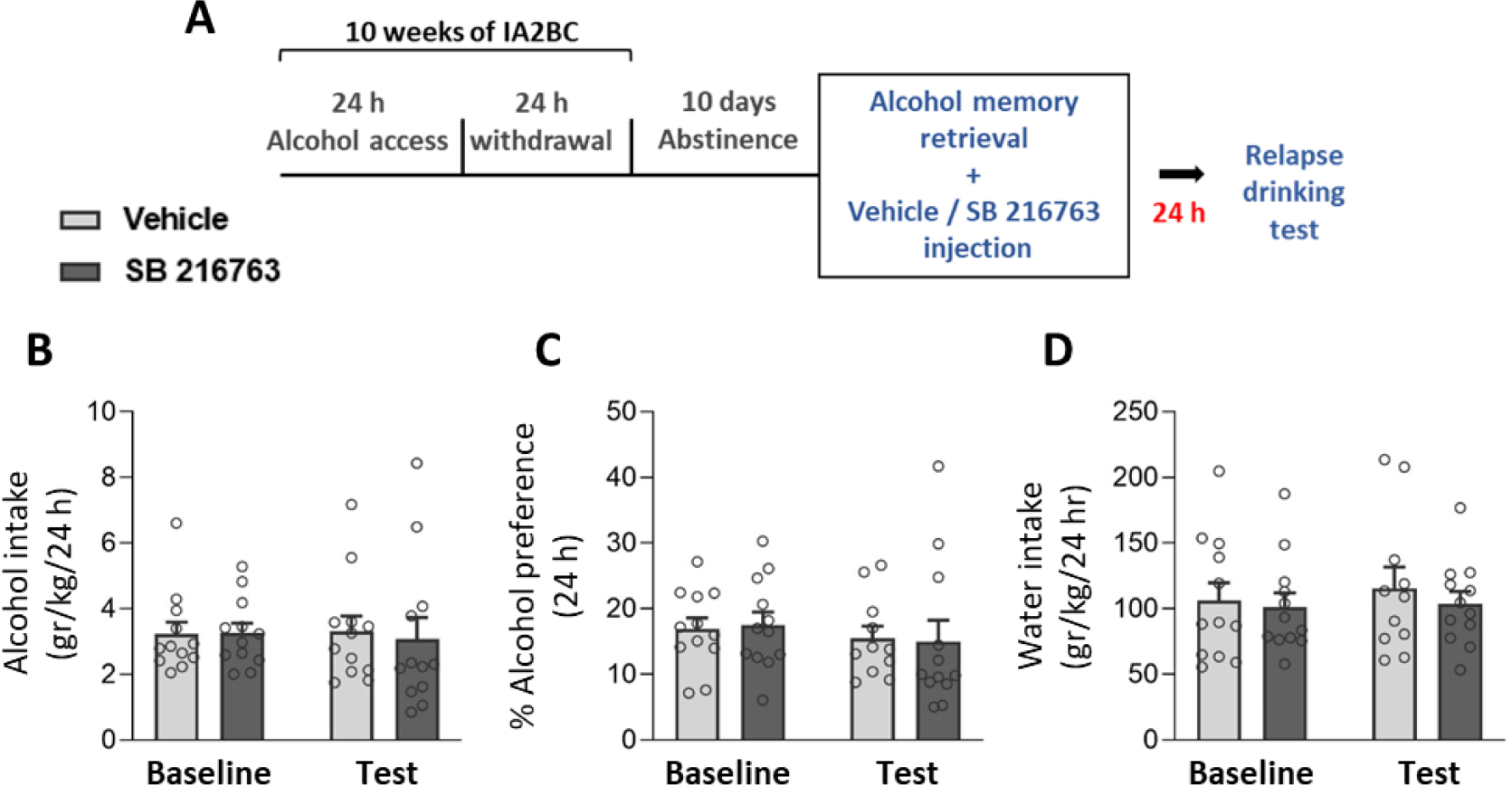
SB 216763 treatment after alcohol memory retrieval has no effect on relapse to alcohol consumption and preference. **A.** Experimental design and timeline. Rats consumed alcohol in the intermittent access to 20% alcohol in a 2-bottle choice paradigm for 10 weeks, followed by 10 days of abstinence. After the abstinence period, alcohol memory was retrieved using an odor-taste cue. Immediately afterward, rats were injected with SB 216763 (2.5 mg/kg) or vehicle. The relapse to alcohol drinking was tested 24 h later. **B-C.** Alcohol intake (B), alcohol preference (C), and water intake (D) in a baseline and relapse test conducted a day after the SB 216763 treatment. Baseline levels of alcohol intake/preference represent the average of the last five sessions prior to abstinence. Bar graphs represent mean ±S.E.M. of alcohol intake/preference. n=11-12.

### Alcohol memory retrieval increases ERK1/2 activation in the amygdala

Although our findings may provide support for the involvement of CRMP2, a target of the PI3K-AKT signaling pathway, in the reconsolidation of alcohol memories, we found no indication for altered activity of AKT, GSK3β or CRMP2 following retrieval of alcohol memory. Therefore, we next set out to explore whether ERK1/2, another major signaling cascade that also serves as a positive regulator of mTORC1 ^28^, is involved in the alcohol memory reconsolidation process.

First, we tested whether the retrieval of alcohol memories affects ERK1/2 phosphorylation, and therefore, its activation. We tested samples from the mPFC, NAc, and amygdala, which were collected 30 min and 60 min after the alcohol memory retrieval (see Experiment 1 in Methods).

As shown in Figure 4B, alcohol memory retrieval did not induce significant changes in the phosphorylation levels of ERK1/2 30 min or 60 min after the memory retrieval (one-way ANOVAs: mPFC [F(2,25)=0.88, p=0.429]; NAc [F(2,27)=0.79, p=0.464]; amygdala [F(2,29)=1.75, p=0.192]) (Figure S1). However, there was a non-significant trend towards increased ERK1/2 phosphorylation in the amygdala after 60 min (Dunnett post hoc: No-retrieval vs. retrieval 60-min, p=0.246). Therefore we next tested for changes in ERK1/2 phosphorylation in the amygdala two hours after memory retrieval (see Experiment 4 in Methods), and found a significant increase in the phosphorylation level of ERK1/2 in the group that underwent retrieval of alcohol memory, compared to no retrieval controls (t-test: t(14)=2.51, p=0.025) (Figures 4D and S2). These findings indicate that amygdalar ERK1/2 activation occurs following alcohol memory retrieval, suggesting that this pathway may be involved in the reconsolidation of alcohol memories.

**Figure 4.**
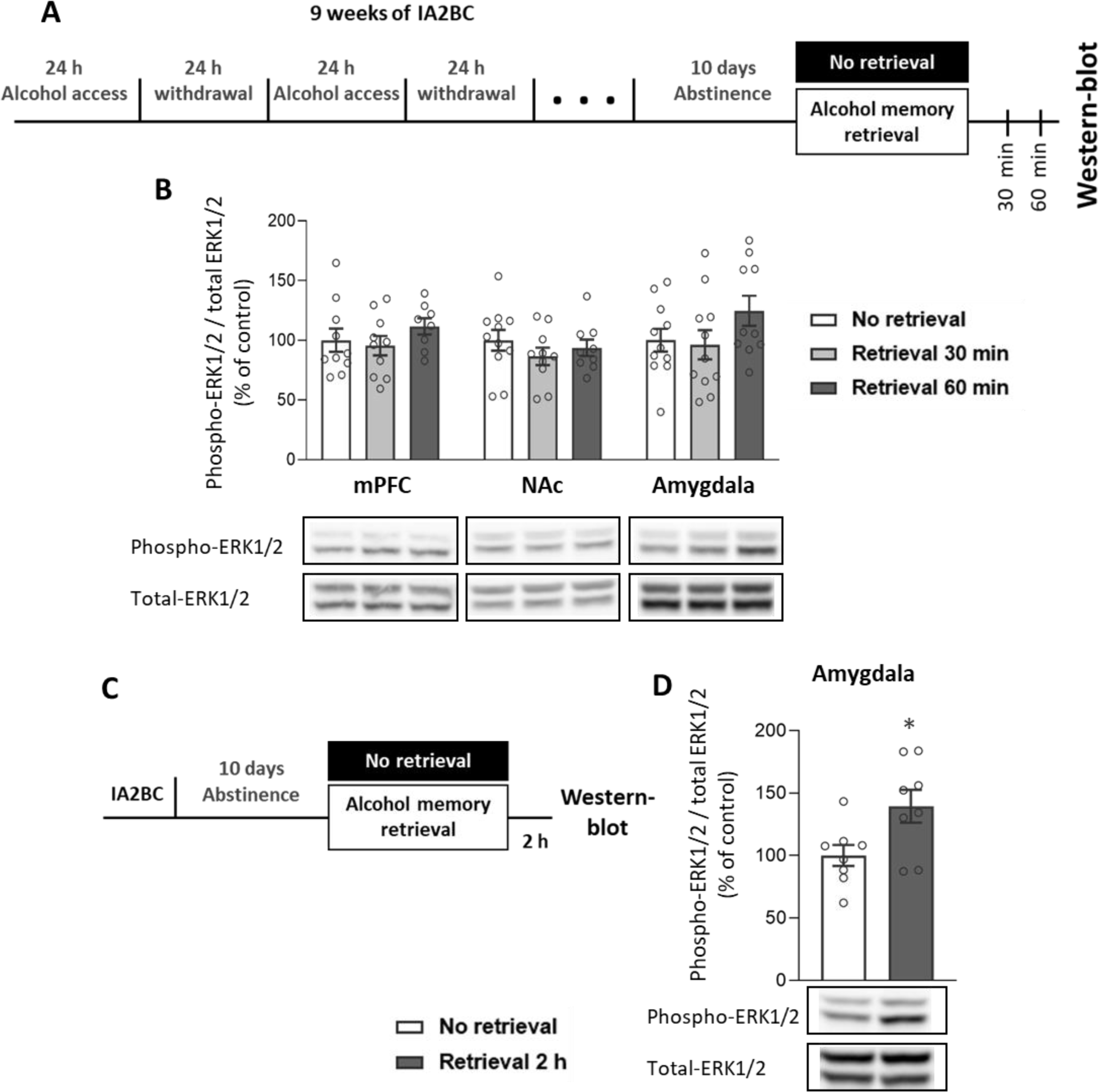
Alcohol memory retrieval increases ERK1/2 phosphorylation in the amygdala. **A.** Experimental design and timeline. Rats consumed alcohol in the intermittent access to 20% alcohol in a 2-bottle choice paradigm for nine weeks, followed by 10 days of abstinence. After the abstinence period, alcohol memory was retrieved using an odor-taste cue. Tissues were collected 30 min and 60 min after the retrieval. Phosphorylation and total protein levels were determined by western blot. Phospho-protein levels of ERK1/2 (Thr202, Tyr204) were normalized to the total protein levels. **B.** Phospho-ERK1/2 (Thr202, Tyr204) levels were normalized to total-ERK1/2 immunoreactivity in the mPFC, the NAc, and the amygdala. **C-D.** A follow-up experiment, in which tissues were collected two hours after the alcohol memory retrieval (C, experimental design and timeline). **D.** Phospho-ERK1/2 levels, normalized to total-ERK1/2 levels. Bar graphs represent mean ±S.E.M. of the percent of change from the control No retrieval group. 30/60 min experiment: n=8-11; 2-h experiment: n=8. * p<0.05.

### ERK1/2 inhibition following alcohol memory retrieval reduces relapse to alcohol consumption

Having shown that alcohol memory retrieval increased the phosphorylation, and therefore the activation, of ERK1/2 in the amygdala, we next set out to establish the causal role of ERK1/2 signaling in the reconsolidation of alcohol memories. To this end, we next tested the effect of post-retrieval ERK1/2 inhibition on relapse to alcohol drinking. We used SL-327 ^73^ – an inhibitor of mitogen-activated protein kinase kinase 1/2 (MEK1/2), the upstream regulator of ERK1/2.

After 10 weeks of alcohol consumption and 10 days of abstinence, alcohol memories were retrieved as described above, and immediately afterward the rats were injected with SL-327 (50 mg/kg, i.p., dose based on Sarantis et al. ^73^) or vehicle. The relapse to alcohol drinking was tested 24 hours later, as well as 14 days later. The rats were abstinent from alcohol between the two tests (see Experiment 4 in Methods).

We found that SL-327-treated rats consumed less alcohol one day and 14 days after the retrieval+treatment, compared with vehicle-treated controls (Figure 5B-C), and demonstrated a lower preference for alcohol one day, but not 14 days, after the treatment (t-test: post-retrieval test – 1 day: alcohol intake [t(23)=3.76, p=0.001], alcohol preference [t(23)=2.87, p=0.009]; post-retrieval test – 14 days: alcohol intake [t(23)=2.28, p=0.033], alcohol preference [t(23)=1.43, p=0.165]). Water intake was not affected by the treatment (t-test: post-retrieval test – 1 day [t(23)=0.46, p=0.650]; post-retrieval test – 14 days [t(23)=0.14, p=0.893]) (Figure 5D).

**Figure 5.**
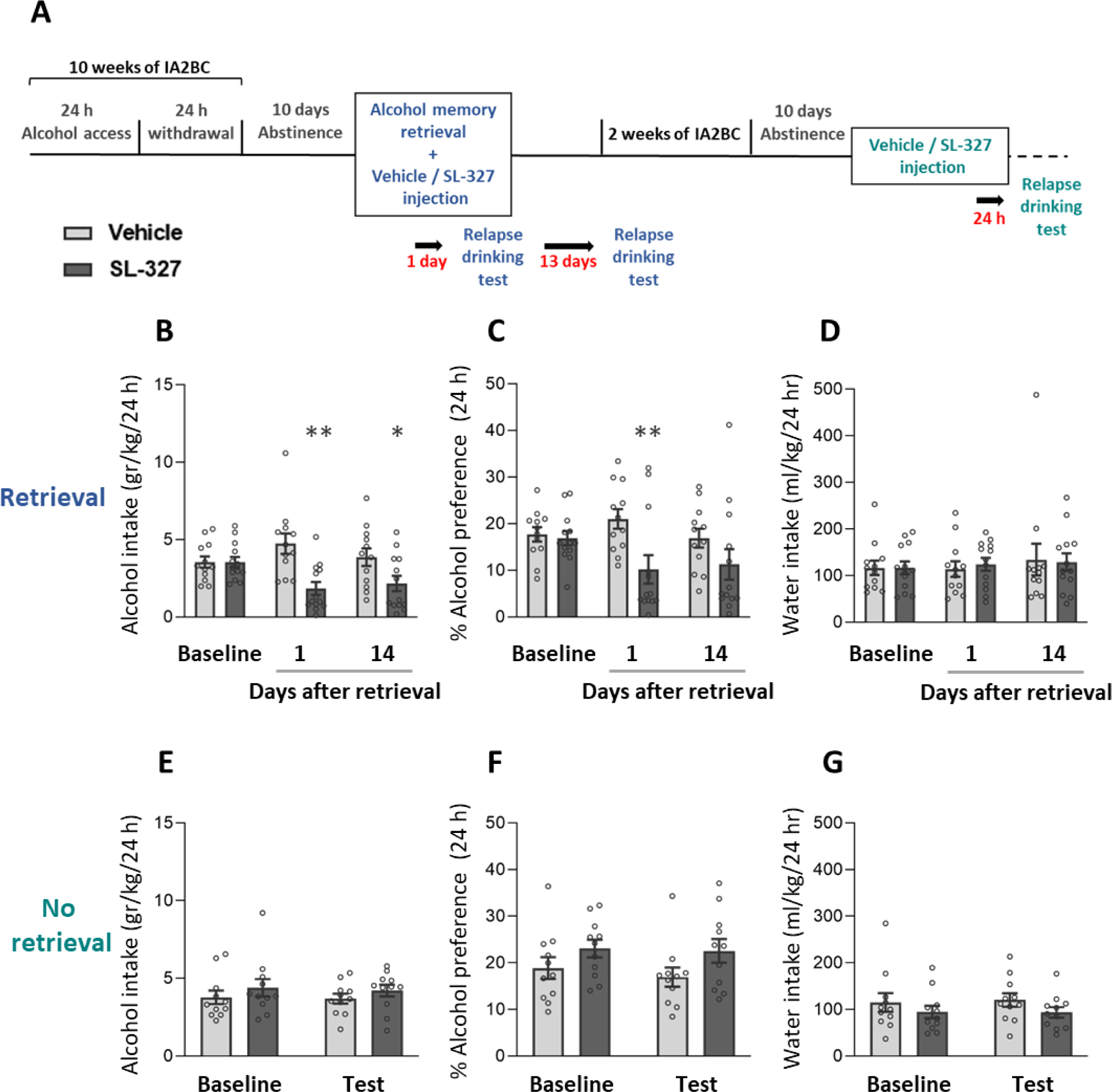
SL-327 treatment after alcohol memory retrieval leads to reduced relapse to alcohol consumption and preference. **A.** Experimental design and timeline. Rats consumed alcohol in the intermittent access to 20% alcohol in a 2-bottle choice paradigm for 10 weeks, followed by 10 days of abstinence. After the abstinence period, alcohol memory was retrieved using an odor-taste cue. Immediately afterward, rats were injected with SL-327 (50 mg/kg) or vehicle. Relapse to alcohol drinking was tested 24 h later and 14 days later. After a 3-week re-training, the rats were subjected again to 10 days of abstinence and were then given an additional treatment of SL-327 or vehicle, without a preceding alcohol memory retrieval, and relapse was tested 24 h later. **B-G.** Alcohol intake (B, E), alcohol preference (C, F), and water intake (D, G) in a baseline and relapse test conducted after SL-327 treatment given following alcohol memory retrieval (B-D) or without memory retrieval (E-G). Baseline levels of alcohol/water intake/preference represent the average of the last five (B-D) or three (E-G) sessions prior to abstinence. Bar graphs represent mean ±S.E.M. of alcohol intake/preference. n=11-13. * p<0.05, ** p<0.01 (Vehicle vs. SL-327).

To further assess whether SL-327’s effects on relapse to alcohol drinking were the result of an impaired reconsolidation process, we re-tested the effects of ERK1/2 inhibition without prior memory retrieval. Thus, rats received drinking re-training for three additional weeks, followed by 10 days of abstinence, and were then, with no prior memory retrieval, treated with SL-327 or with vehicle, in a counterbalanced manner. Twenty-four hours after the treatment, an additional relapse to drinking test was held.

We found that in the absence of prior alcohol memory retrieval, SL-327 treatment had no effects on the rats’ alcohol or water drinking or on alcohol preference (Figure 5E-G, t-tests: alcohol intake [t(20)=1.05, p=0.305]; alcohol preference [t(20)=1.72, p=0.101]; water intake [t(20)=1.46, p=0.161]).

Together, these results indicate that inhibition of ERK1/2 reduces relapse to alcohol consumption in a long-lasting manner, but only when applied following alcohol memory retrieval, suggesting that inhibition of ERK1/2 impairs alcohol memory reconsolidation.

Finally, as we found that alcohol memory retrieval increased ERK1/2 phosphorylation in the amygdala, we validated that systemic SL-327 injection indeed inhibits ERK1/2 phosphorylation in the amygdala. In addition to ERK1/2, we also examined changes in the phosphorylation level of S6 ribosomal protein, a downstream effector of mTORC1, which was shown to be elevated in the CeA following alcohol memory retrieval ^17^ (see Experiment 5 in Methods). For this validation, we injected rats with SL-327. Amygdala tissues were dissected 90 min after the injection ^63,64^ and were processed for western blot analysis. As expected, SL-327 treatment led to reduced phosphorylation of ERK1/2 (t-test: t(11)=5.24, p=0.0003). Surprisingly, a trend toward increased phosphorylation of S6 was observed following the SL-327 treatment (t-test: t(12)=2.06, p=0.06) (Figures 6 and S3).

**Figure 6.**
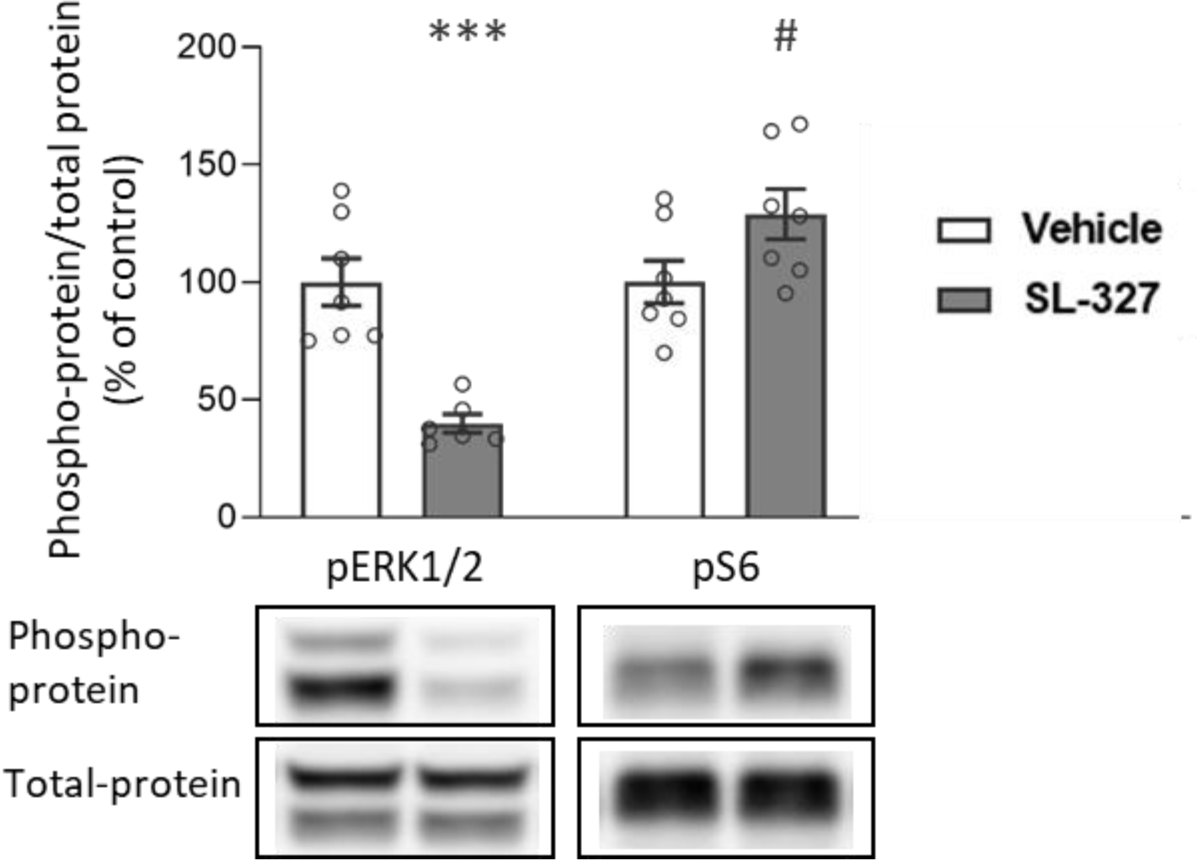
SL-327 injection leads to reduced ERK1/2 phosphorylation in the amygdala. Rats were injected with SL-327 (50 mg/kg) or with vehicle. Amygdala tissues were collected 90 min after the injection. Phosphorylation and total protein levels were determined by western blot. Phospho-protein levels of ERK1/2 (Thr202, Tyr204) and S6 (Ser235/236) were normalized to the total-protein immunoreactivity. Bar graphs represent mean ±S.E.M. of the percent of change from the control group. n=6-7. # p=0.06, ** p<0.01.

These results imply that the observed effects of ERK1/2 inhibition on alcohol memory reconsolidation and relapse are not necessarily mediated by a diminished activity of mTORC1.

## 4. Discussion

We show here that the retrieval of alcohol-associated memories in animals with a history of long-term alcohol drinking evokes the amygdalar activation of ERK1/2 signaling, but not of the PI3K-AKT signaling pathway. Furthermore, we demonstrate that the post-retrieval inhibition of the ERK1/2 pathway results in long-lasting suppression of relapse to alcohol drinking, suggesting that this signaling pathway plays a role in alcohol memory reconsolidation. In addition, we demonstrate suppressed relapse following the post-retrieval inhibition of CRMP2, a translational product of mTORC1 signaling. In contrast, the post-retrieval inhibition of GSK3β, an effector of the PI3K-AKT signaling pathway that regulates CRMP2 activity ^70^, did not affect relapse to alcohol consumption. Together, our findings suggest that the ERK1/2 signaling pathway, as well as CRMP2, plays a critical role in alcohol memory reconsolidation, and that these pathways may provide targets for preventing relapse to alcohol drinking.

### ERK1/2 is crucial for alcohol memory reconsolidation

Our results suggest that ERK1/2 has a crucial role in the reconsolidation of alcohol memories. These findings are in line with previous reports that showed that the reinstatement of operant alcohol-seeking behavior, evoked by exposure to cues that were present during the self-administration training, led to increased ERK1/2 activation in the basolateral amygdala (BLA) ^41,42^ and the NAc shell ^41^. Here, we show that the mere exposure to the drug-associated cues is sufficient to induce ERK1/2 activation, even in the absence of the operant-seeking behavior.

Importantly, the increase in ERK1/2 activation was observed in the amygdala, but not in the mPFC and the NAc, which are also known as key regions for drug memory reconsolidation ^17,33,44,56,62,74–76^. These results are aligned with the findings of Barak et al. that demonstrated increased activation of mTORC1, following retrieval of alcohol memories, using two different retrieval methods ^17^. Thus, when the memory was retrieved by exposure to the context of the alcohol self-administration chamber, mTORC1 activation was increased in the CeA and the prelimbic and orbitofrontal cortices. However, when the retrieval was conducted in the home cages by the presentation of odor-taste cues, as in the present study – mTORC1 activation was increased only in the CeA. These observations resonant the view according to which the amygdala encodes the Pavlovian association between the unconditioned stimulus (*i.e.* the reinforcing effect of alcohol) and the conditioned stimuli (*i.e.* the odor and taste of alcohol), while cortical regions, as well as the NAc, encodes the instrumental and motivational aspects of the memory ^7,17,56^.

Interestingly, inhibition of ERK1/2, systemically or locally in the mPFC, was previously shown to increase alcohol-reinforced operant responding ^52,54^ and alcohol drinking in the home cage ^53^. Here, we show that when ERK1/2 inhibition is applied following alcohol memory retrieval, it interferes with the reconsolidation of the alcohol memory, leading to the opposite outcome, i.e., reduced relapse to alcohol drinking in the home cage.

The essential role of ERK1/2 in the reconsolidation process has been demonstrated repeatedly in studies of fear memory reconsolidation ^77–81^. Furthermore, previous studies have shown that systemic ^44,45^, intra-BLA ^43^, or intra-NAc ^46^ ERK1/2 inhibition in conjunction with drug-memory retrieval disrupts the reconsolidation of drug memories, as reflected by suppressed subsequent cocaine and morphine-seeking behavior. As far as we know, the present study provides the first evidence for the involvement of ERK1/2 in the reconsolidation of alcohol memories.

Our present finding that ERK1/2 is activated in the amygdala following alcohol memory retrieval, and that its inhibition disrupts alcohol memory reconsolidation, taken together with our previous finding that mTORC1 inhibition in the amygdala yields similar results ^17^, led us to test whether both effects shared a common pathway. Specifically, ERK1/2 is a positive regulator of mTORC1 ^27,28,40^, so ERK1/2 inhibition may have impaired mTORC1 activation, which is essential for the reconsolidation process ^17^. Surprisingly, we found that the systemic administration of the MEK1/2 inhibitor did not result in reduced phosphorylation of S6, which is often used as a marker for mTORC1 activity ^17,30,51^. Therefore, it is not likely that ERK1/2 and mTORC1 share the same mechanism of action in alcohol memory reconsolidation disruption.

Importantly, ERK1/2 is a key component of various intracellular signaling pathways that underlie synaptic and neuronal plasticity ^82–85^. ERK1/2 signaling is crucial for the activation of several transcription factors that were implicated in memory reconsolidation, including ETS-like gene-1 (ELK-1) ^45^ and cAMP response element binding (CREB) ^45,62,86,87^. Indeed, cocaine and morphine memory retrieval led to increased activation of ERK1/2 and CREB in the NAc, which was suppressed by ERK1/2 inhibition ^45,46^. Moreover, we have recently demonstrated that retrieval of alcohol memories, by exposing mice to the alcohol-associated context, led to increased activation of CREB in the mPFC and dorsal hippocampus, and was followed by elevated mRNA expression of the CREB transcriptional targets, *Arc* and *Zif268* ^62^. Post-retrieval downregulation of *Arc* expression in the dorsal hippocampus disrupted the expression of alcohol place preference ^62^. Therefore, it is plausible that the impaired reconsolidation that we observed as a result of ERK1/2 inhibition, was mediated via the MAPK-CREB pathway, rather than the mTORC1 pathway.

### CRMP2 inhibition impairs alcohol memory reconsolidation

Our findings also suggest a possible contribution of CRMP2 to the reconsolidation of alcohol memories. Specifically, we found that post-retrieval inhibition of CRMP2, by the FDA-approved drug lacosamide, led to impaired reconsolidation of the alcohol memory, which was reflected in diminished relapse to alcohol drinking. Nonetheless, we found no effect of alcohol memory retrieval on the expression of CRMP2 protein or on the phosphorylation of CRMP2 in the site that GSK3β phosphorylates. Conversely, alcohol-priming-induced reinstatement of alcohol place preference was previously shown to increase NAc CRMP2 levels ^51^. Increased expression of CRMP2 was also observed in the mPFC after exposure to cocaine-associated cues during reinstatement test for operant cocaine-seeking ^88^. Our observation, which indicates a lack of change in CRMP2 protein level after alcohol memory retrieval, raises the possibility that exposure to drug-related cues alone, in the absence of exposure to the drug itself or behavioral expression of drug seeking, is not sufficient to induce changes in CRMP2 expression.

Systemic inhibition of CRMP2 using lacosamide was previously shown to decrease alcohol binge drinking ^69^ and to prevent the reinstatement of alcohol place preference ^51^ when the treatment was administered shortly before the behavioral test. Here, we demonstrate that lacosamide also interferes with the reconsolidation of the alcohol memory, consequently reducing relapse to alcohol drinking. CRMP2 serves as a multifunctional protein, mainly known for its ability to promote neurite outgrowth by regulating microtubule dynamics ^48,50,89^. Therefore, the activity of CRMP2 is essential for morphological changes that are involved in synaptic plasticity ^90,91^. The CRMP2 inhibitor we used, (*R*)-Lacosamide (Vimpat), impairs neurite outgrowth by diminishing the ability of CRMP2 to promote tubulin polymerization ^67,92^. Thus, our results suggest that CRMP2-mediated tubulin polymerization is crucial for memory reconsolidation. A note of caution is due here since lacosamide is an anti-epileptic drug that was shown to reduce neuronal excitability by enhancing slow inactivation of voltage-gated sodium channels ^68,93^. Therefore, the observed effect of lacosamide on the reconsolidation of alcohol memories is not necessarily mediated by CRMP2-dependant mechanisms. This possibility may also explain our finding that GSK3β inhibition did not affect alcohol memory reconsolidation, although GSK3β is a negative regulator of CRMP2 ^47,70^. Interestingly, GSK3β inhibition was shown to disrupt the reconsolidation of fear-^71^ and drugs-^33,34,37,94^ memories. It has been speculated ^71,95^ that this effect stems from the ability of GSK3β inhibitors to block LTD induction ^72,96^, which was suggested to be important for the maintenance of previously established memories ^97^. Nonetheless, unlike the reconsolidation of other types of memories, our findings suggest that GSK3β does not mediate the reconsolidation of alcohol memories.

To conclude, our findings point to the crucial role of ERK1/2 and CRMP2 in the reconsolidation of alcohol-associated memories and subsequent relapse. Nevertheless, our results do not provide a definitive elucidation of the signaling pathways governing alcohol memory reconsolidation. Future research endeavors should aim to establish these connections more comprehensively. Importantly, the ability of the CRMP2 inhibitor lacosamide, an FDA-approved drug, to disrupt the reconsolidation of alcohol memories, can pave the way to a novel therapeutic option for the treatment of alcohol use disorder.

## Acknowledgments

The research was supported by funds from the United States – Israel Binational Science Foundation (BSF) grant 2017022 and from the Israel Science Foundation (ISF) grant 508/20. Segev Barak is the Stephen Harper Chair of Translational Neuroscience at the Faculty of Social Sciences, Tel Aviv University.

## Author contributions

NR and SB designed the research; NR, ML, CA, HL, TG and NU performed the research; NR and SB analyzed the data and wrote the paper.

## Supplementary data

Supplementary file includes images of the uncropped full-length western-blot bands.

