## Supplementary data for "ERK1/2 inhibition disrupts alcohol memory reconsolidation and prevents relapse"

***Supplementary Figure 1***

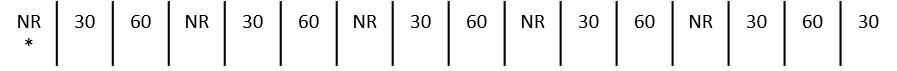

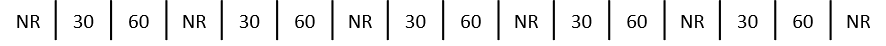

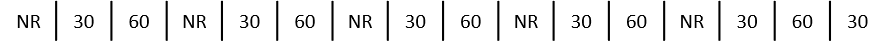

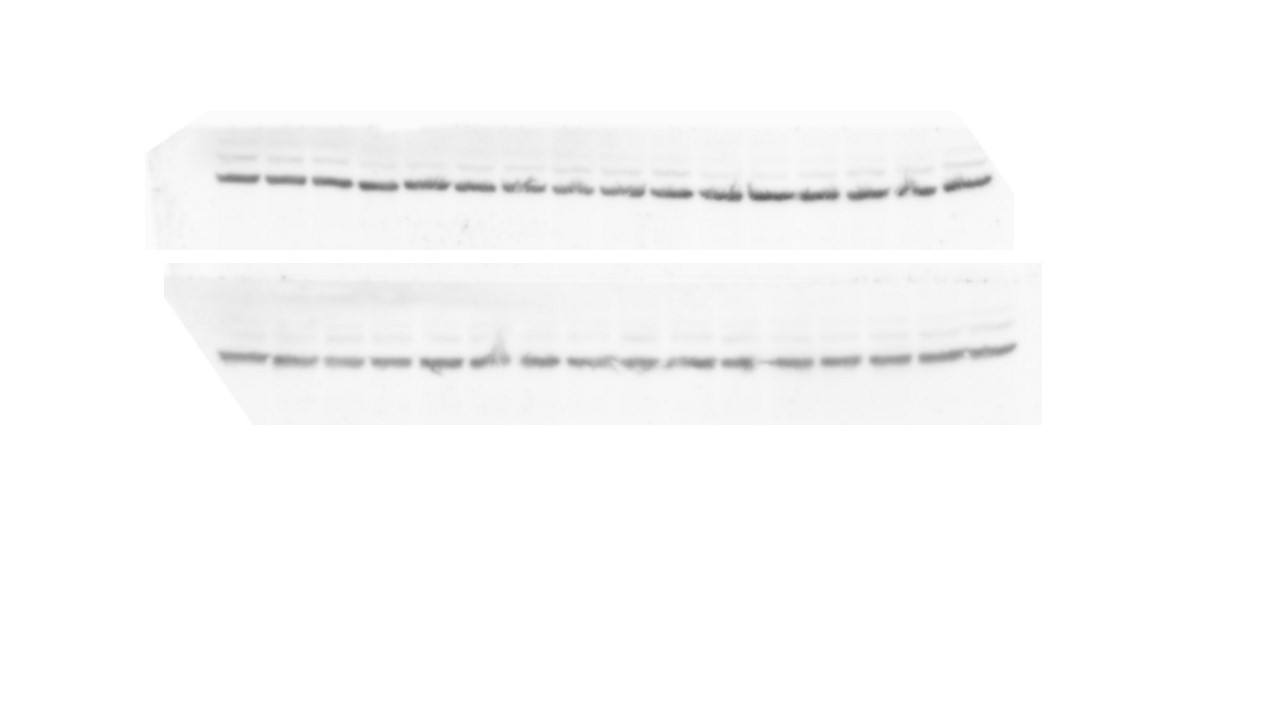

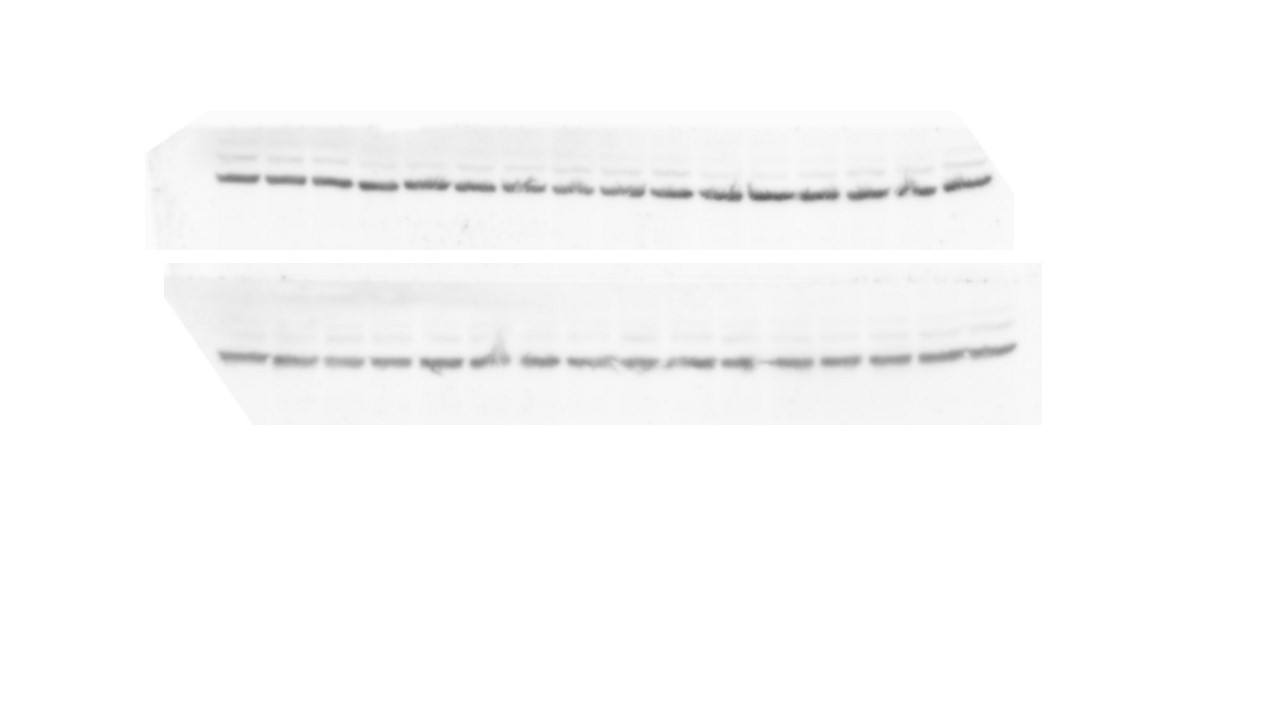

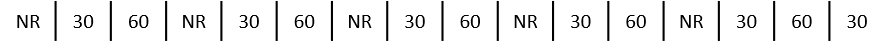

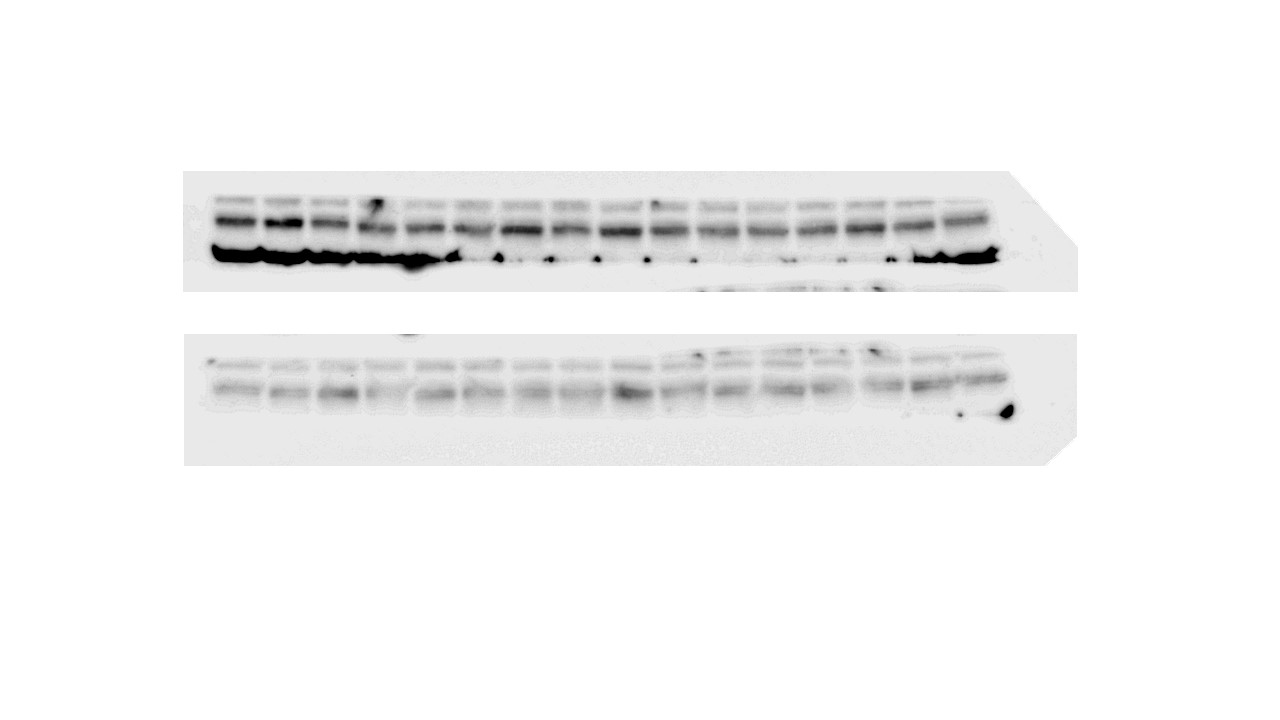

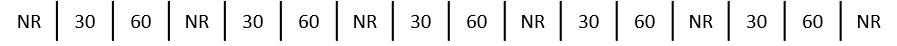

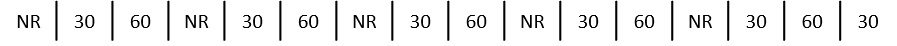

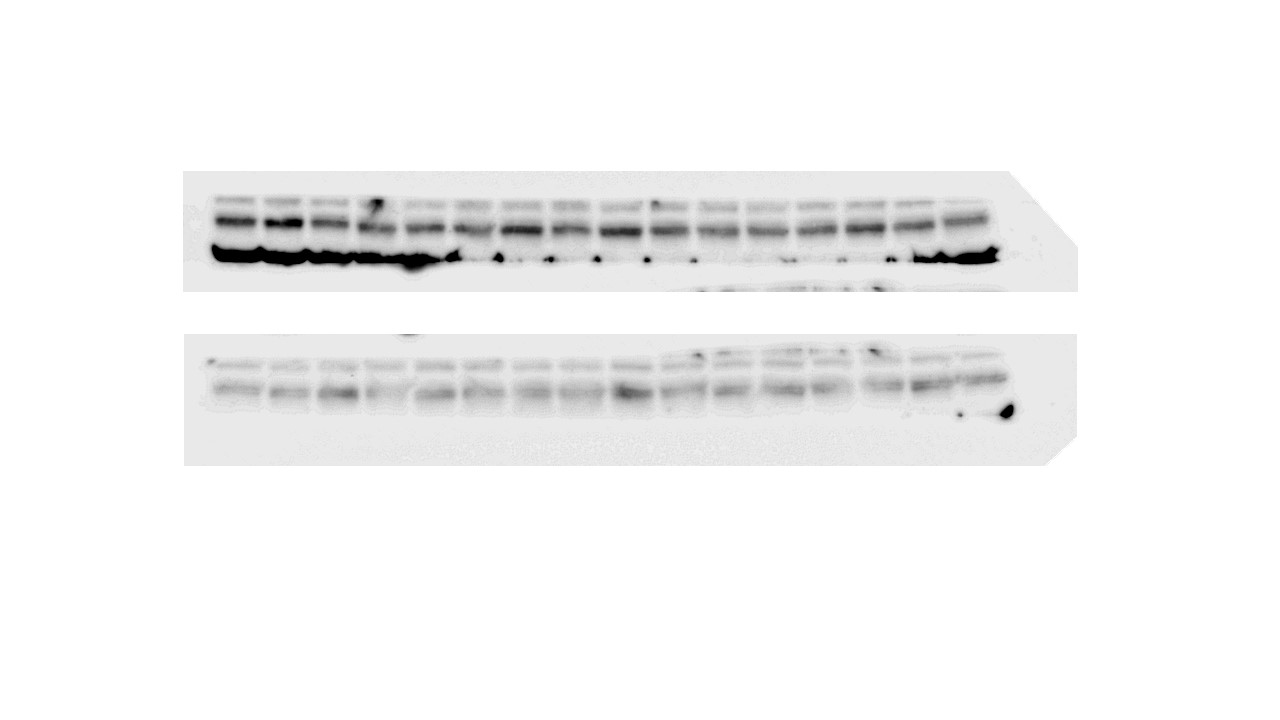

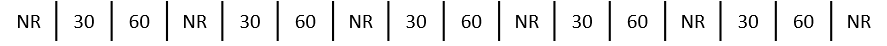

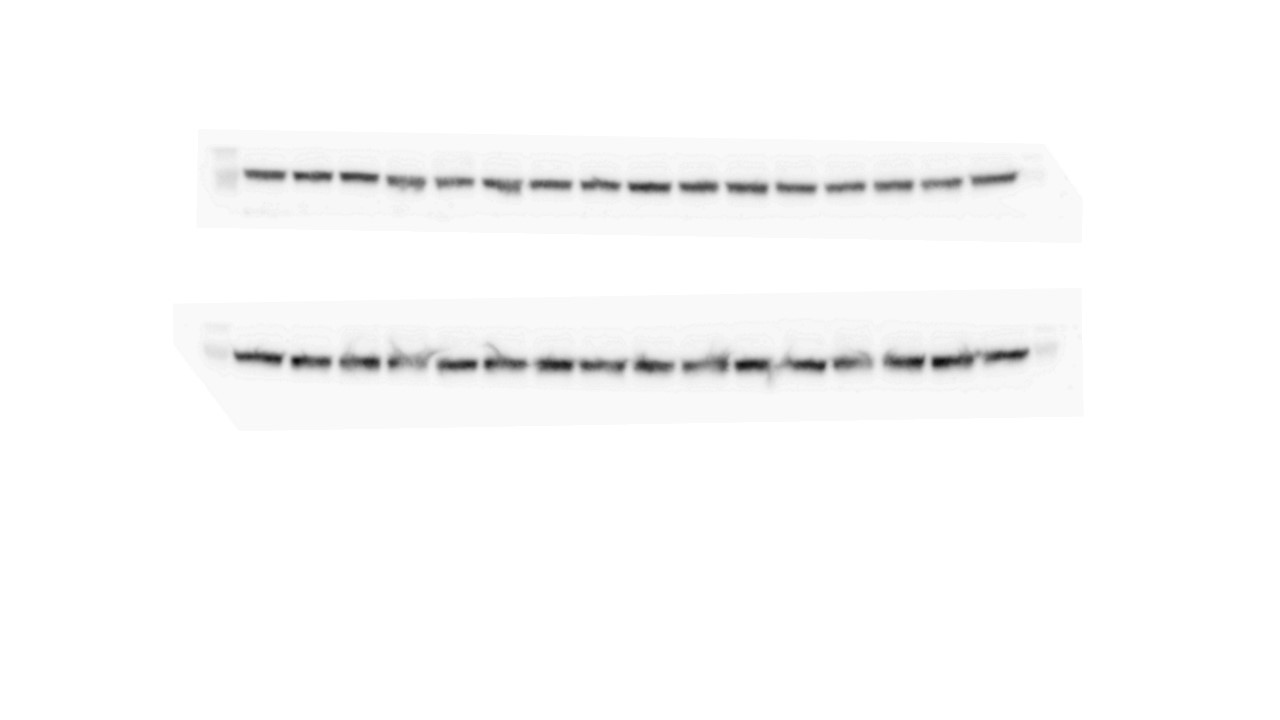

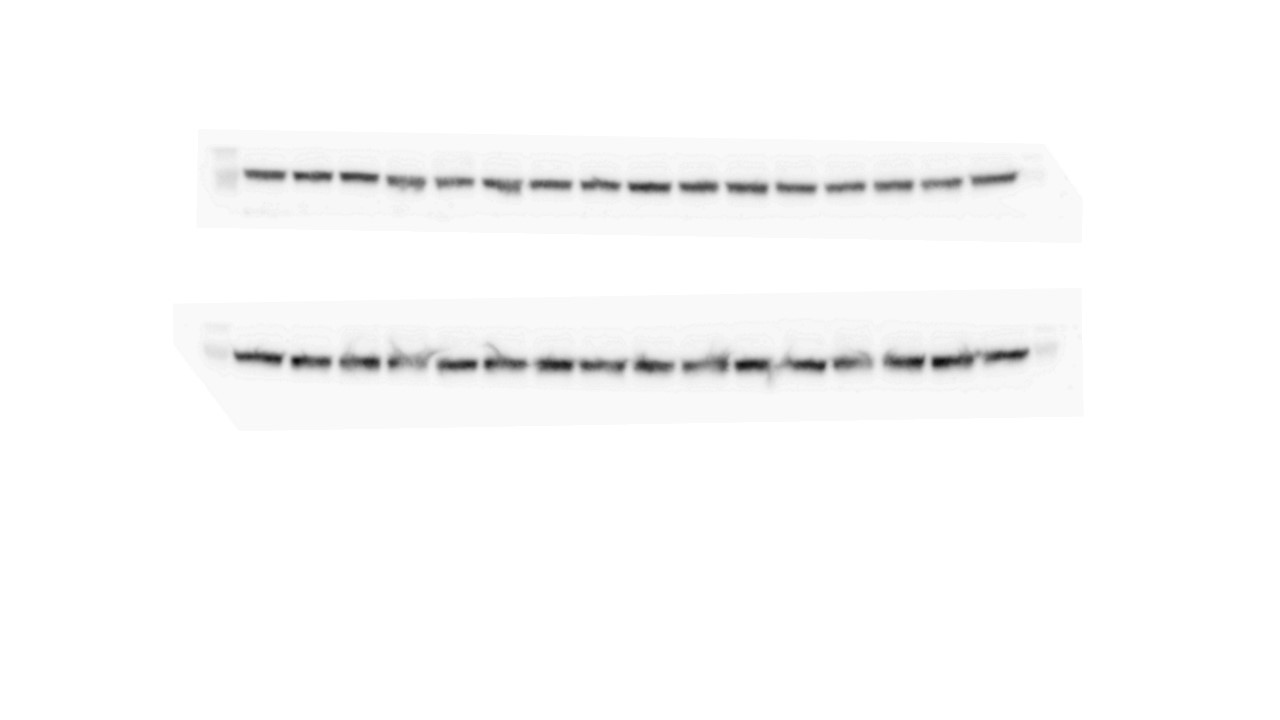

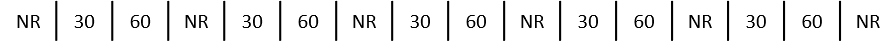

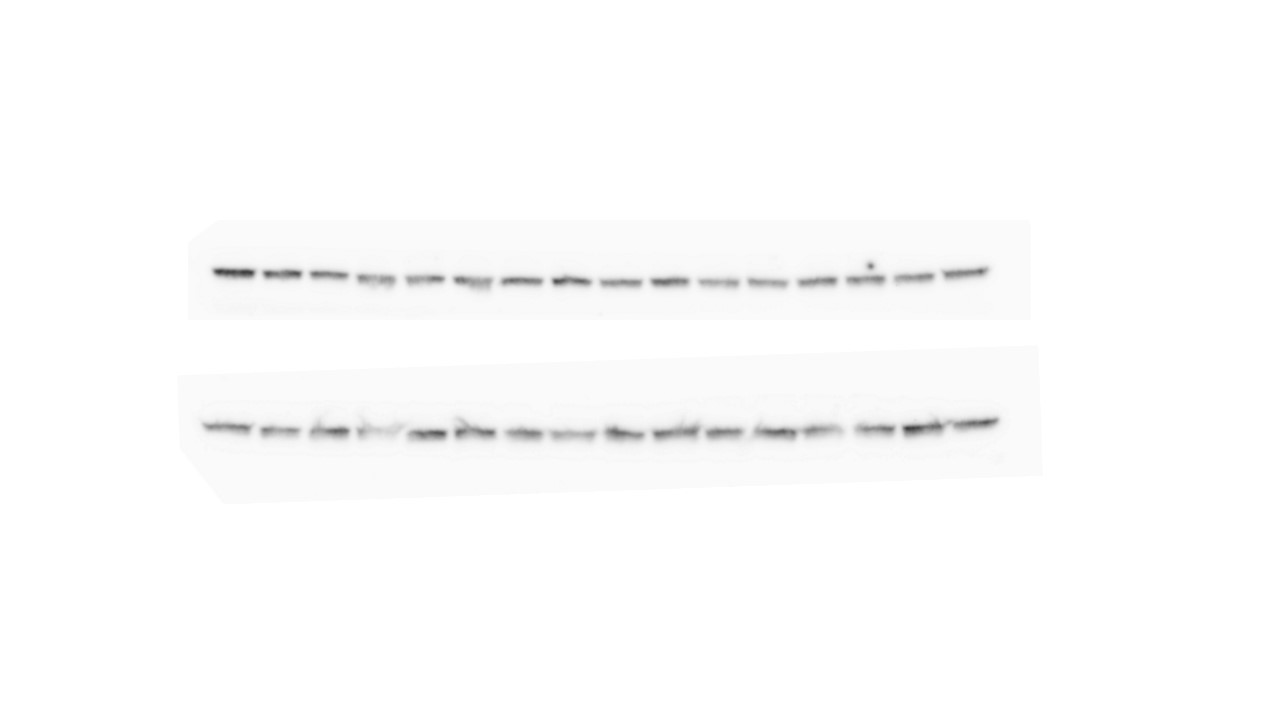

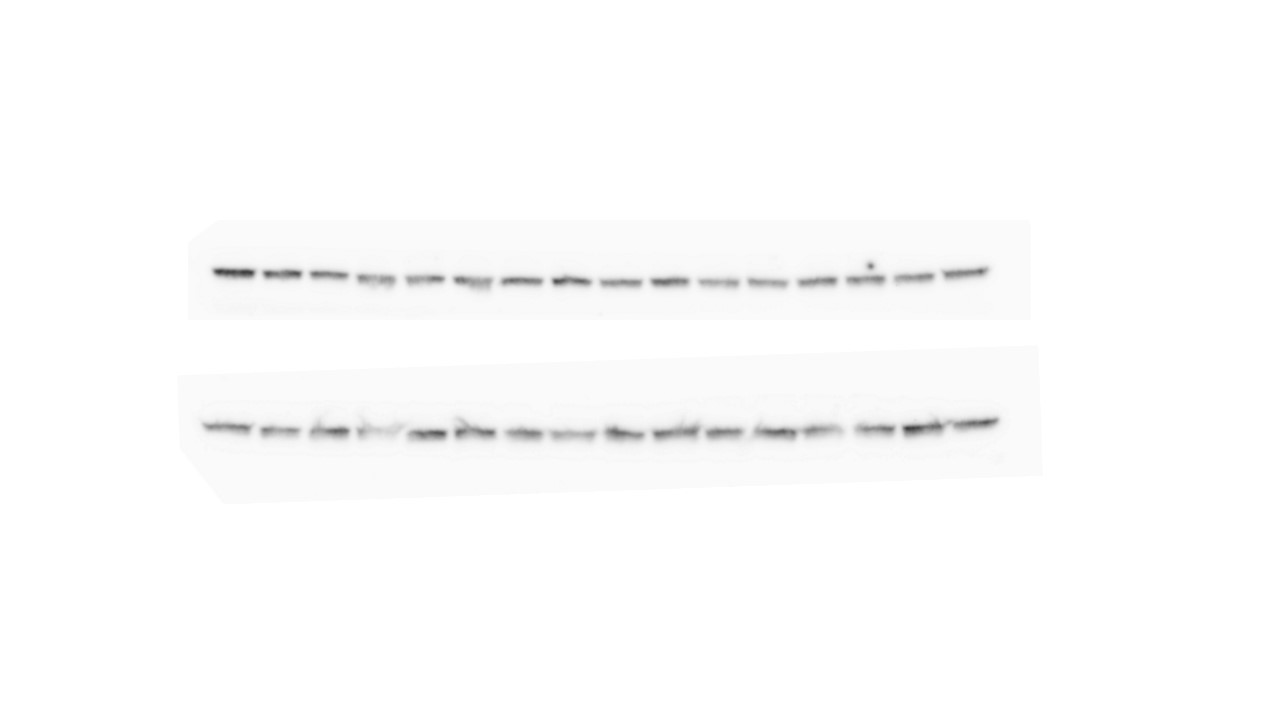

GAPDH – 37 kDa

Total AKT - 60 kDa

pAKT-Ser473 – 60 kDa

**A**

**mPFC**

pAKT-Thr308 – 60 kDa

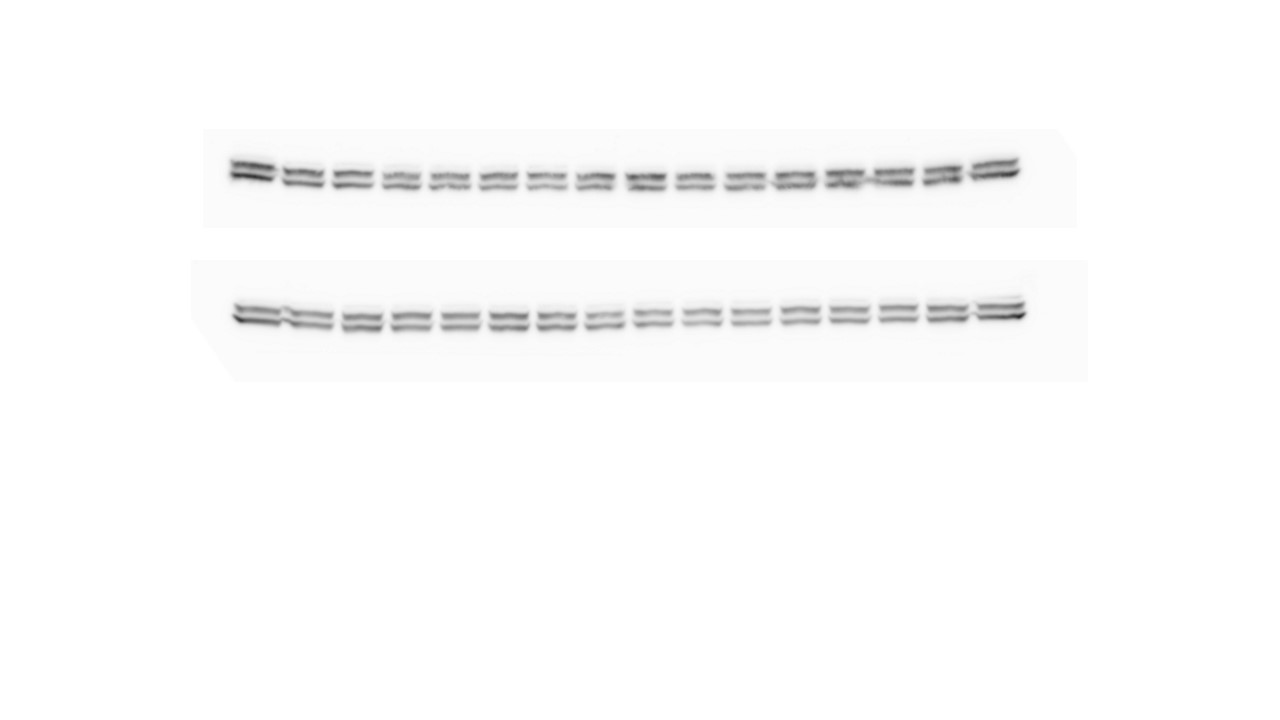

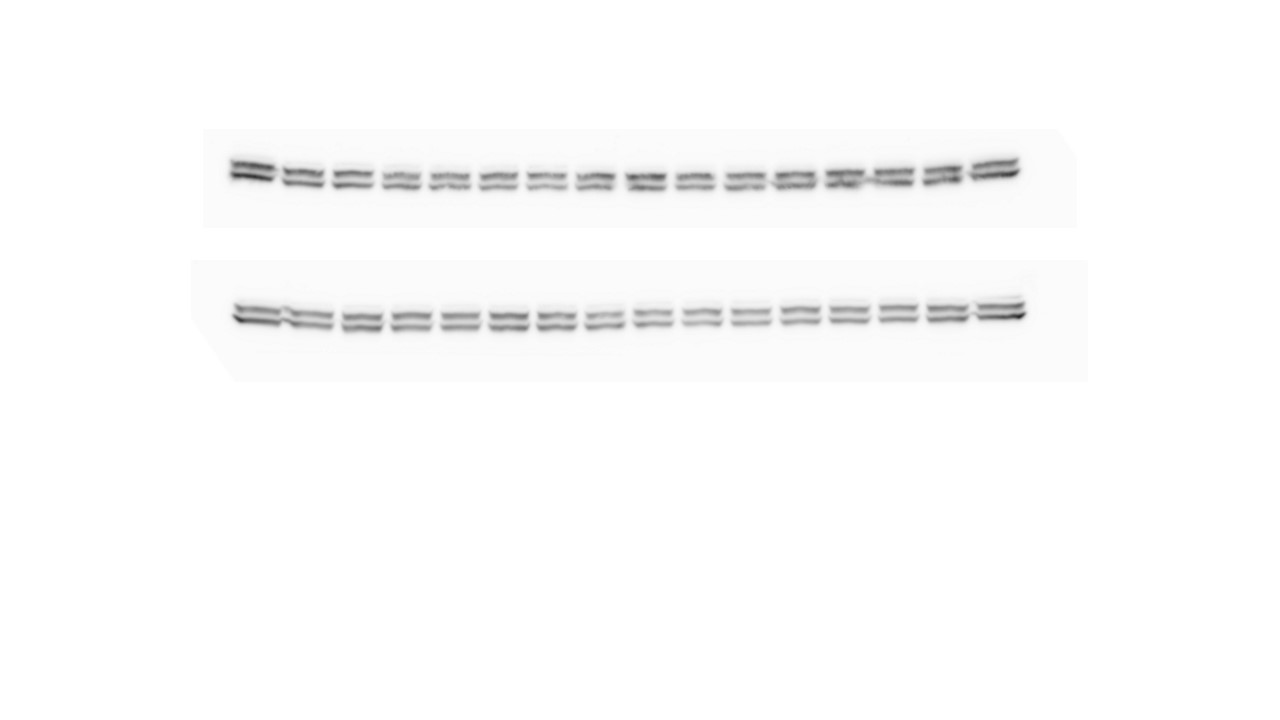

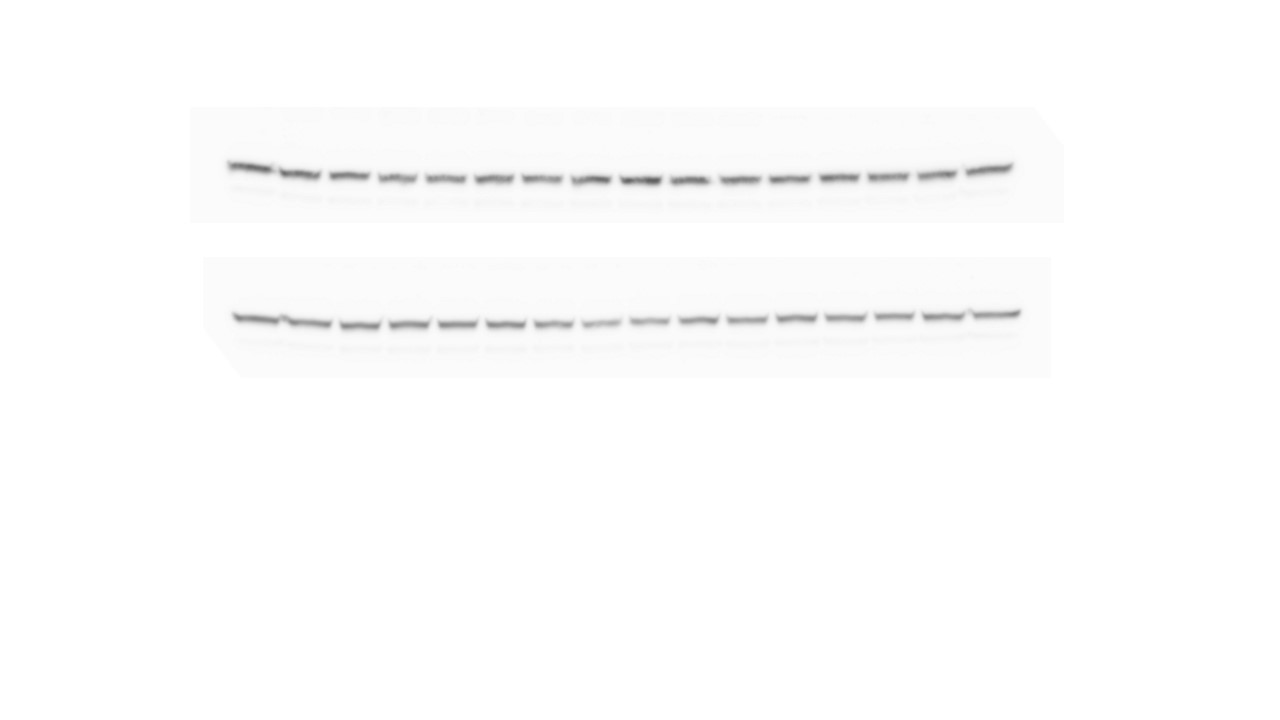

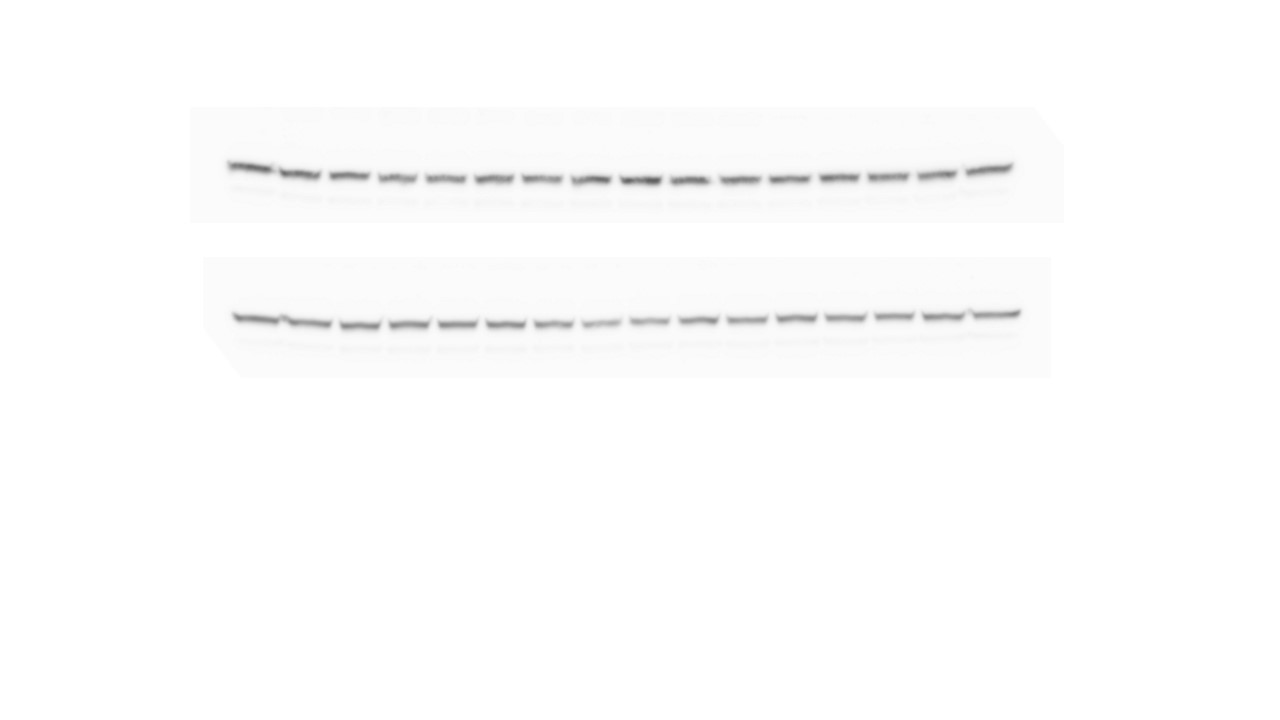

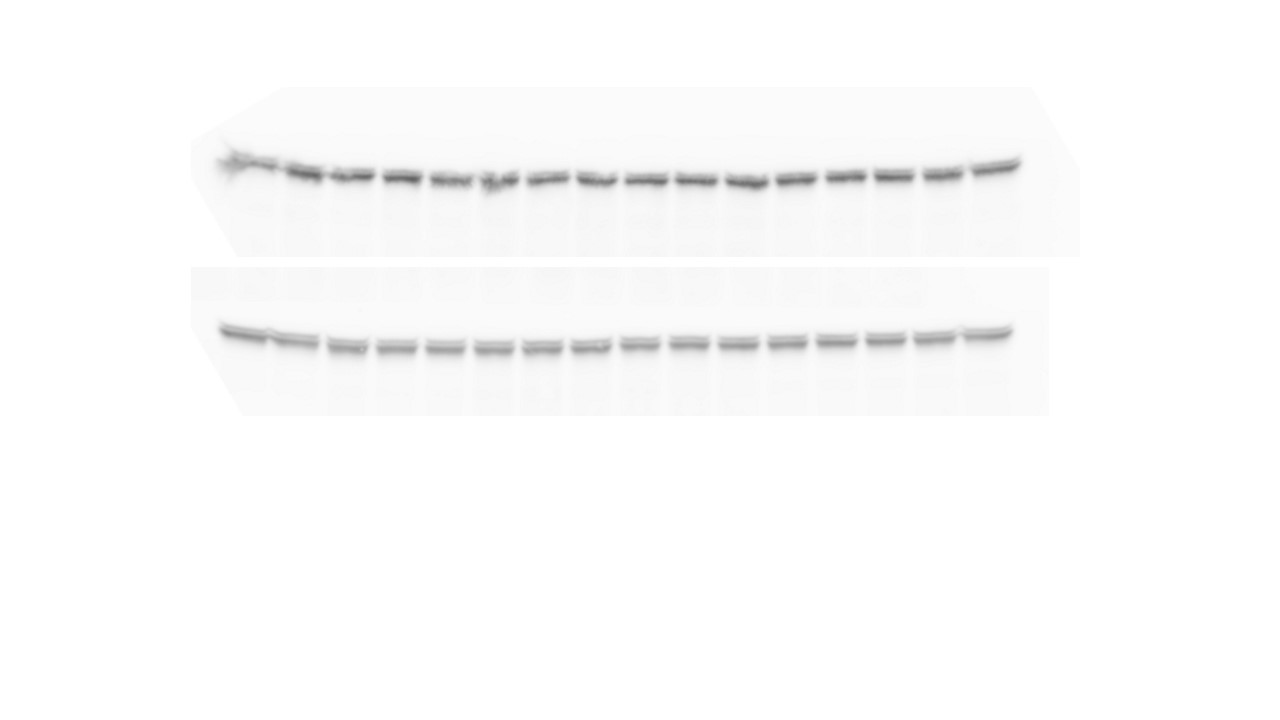

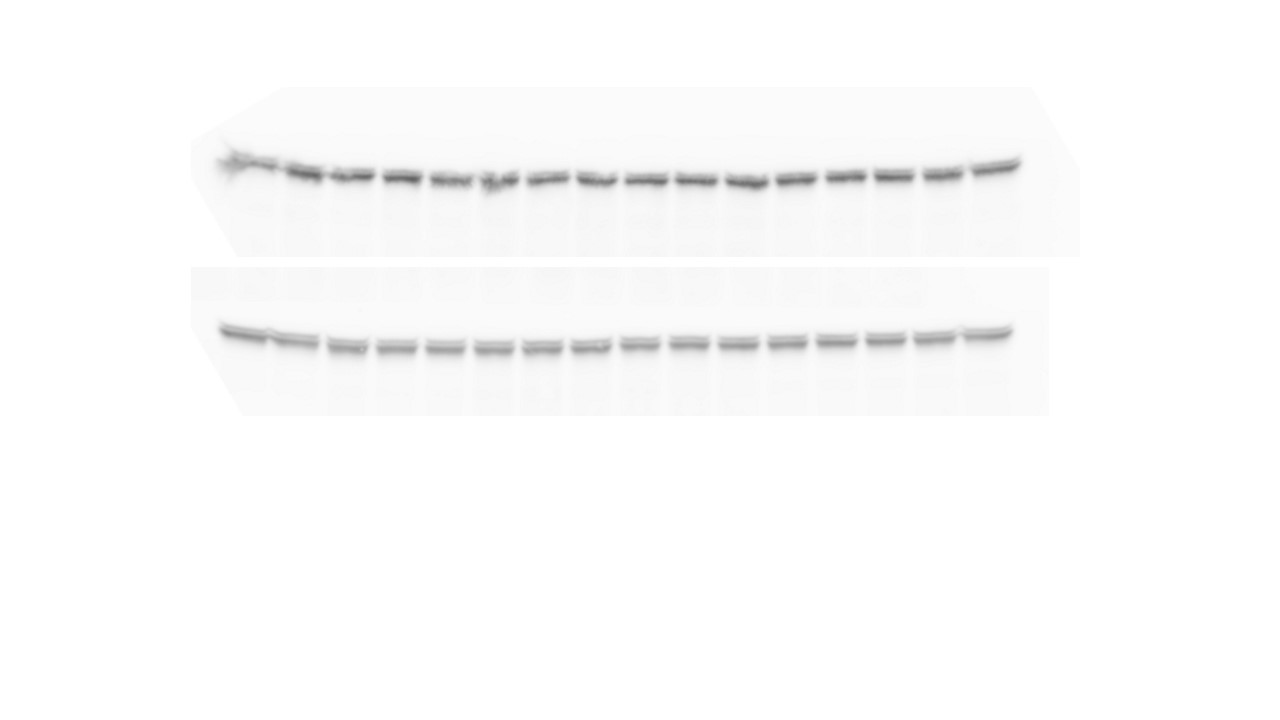

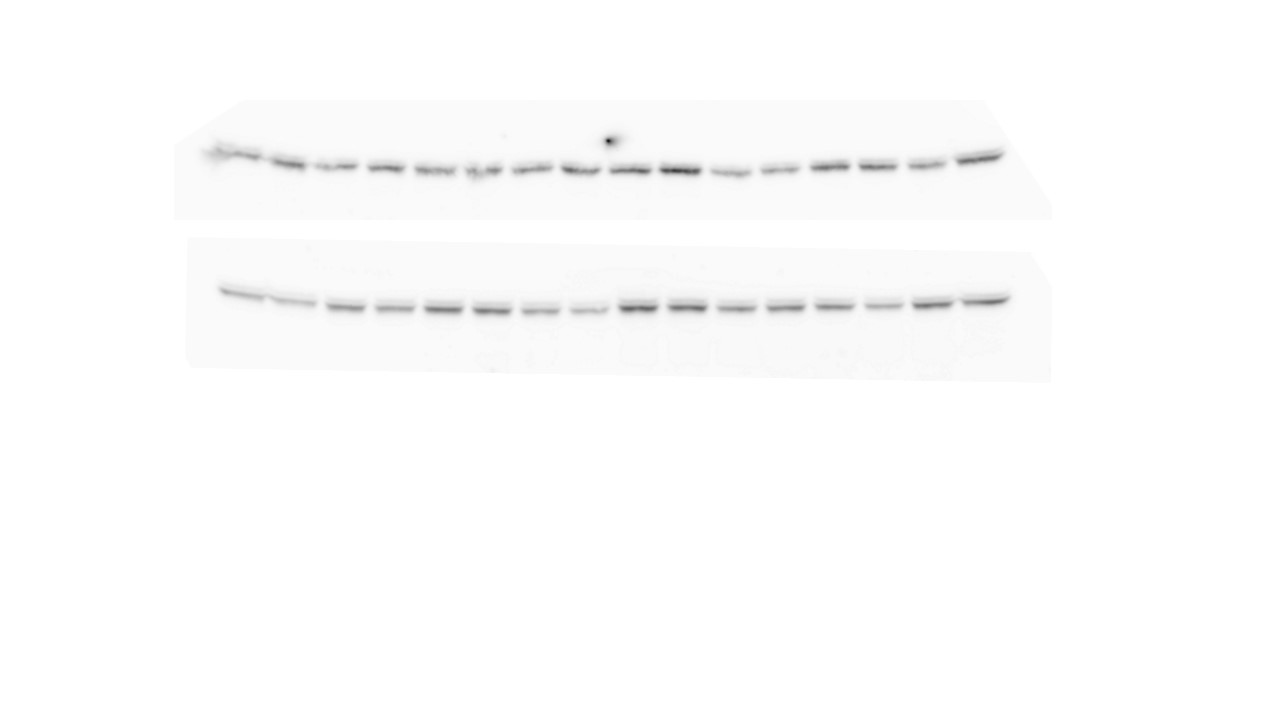

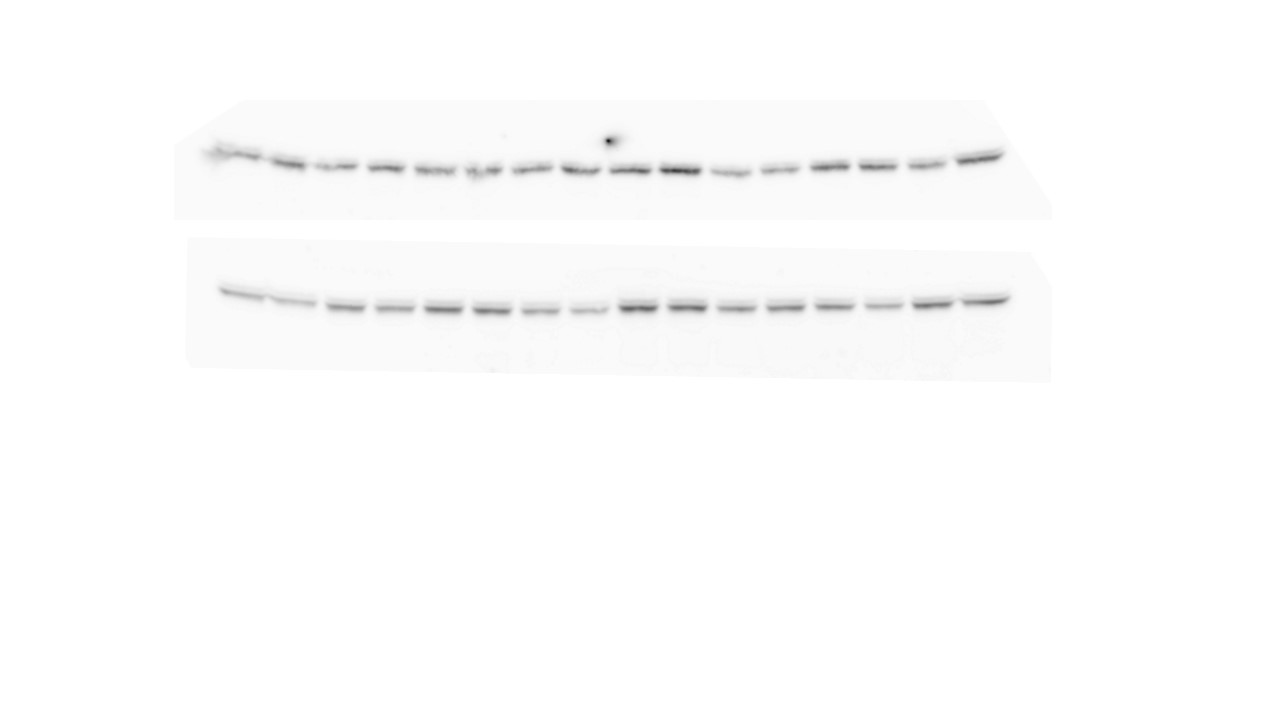

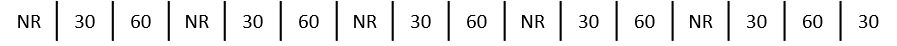

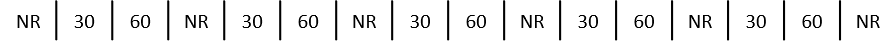

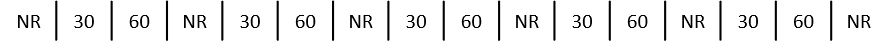

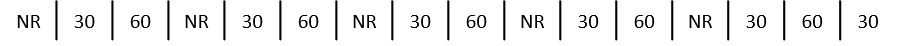

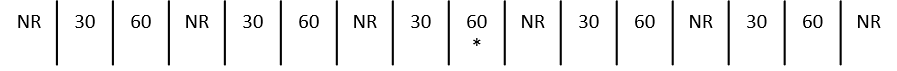

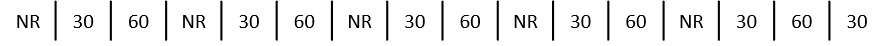

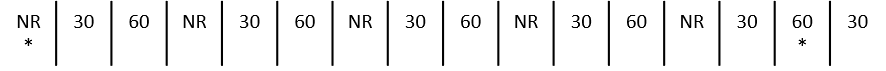

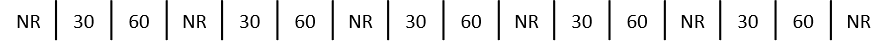

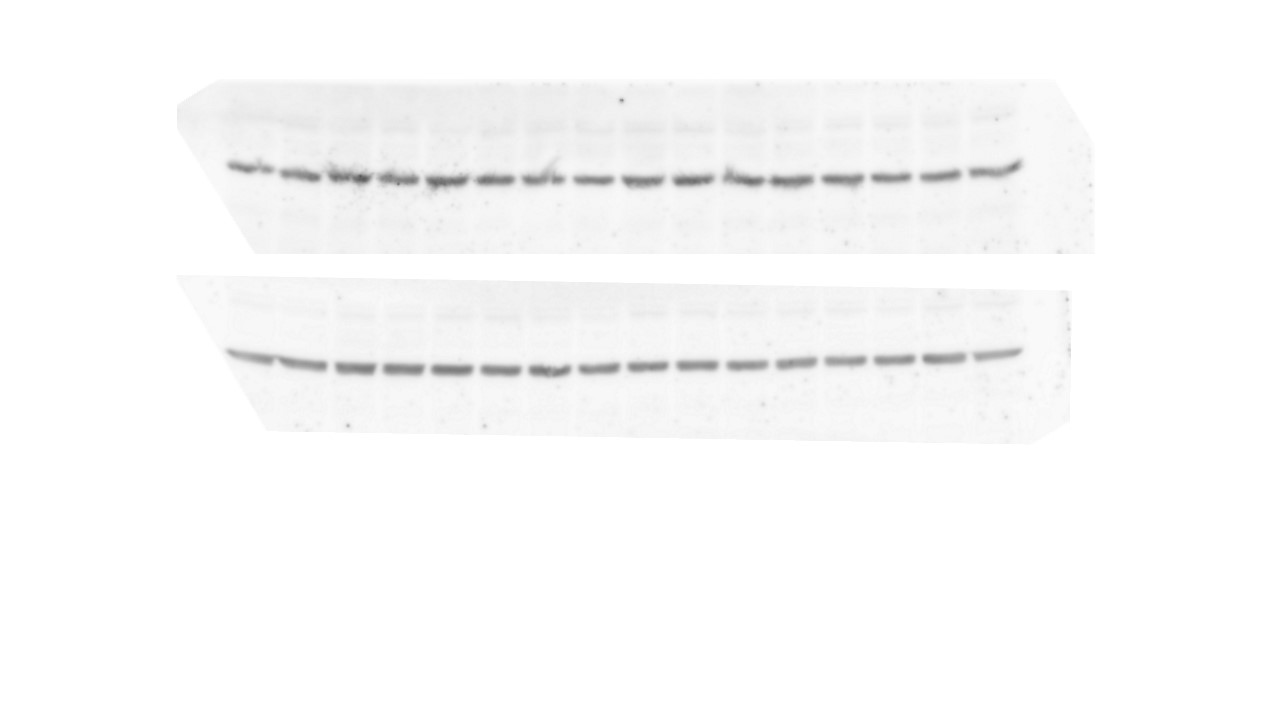

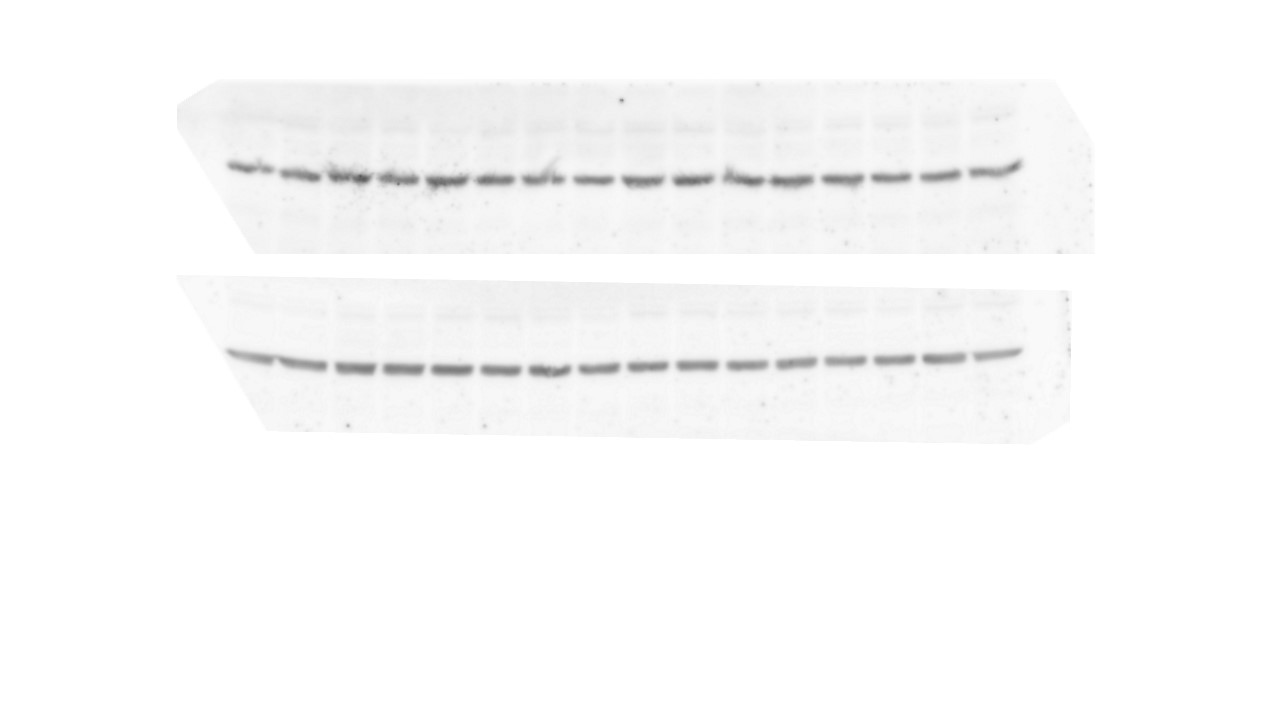

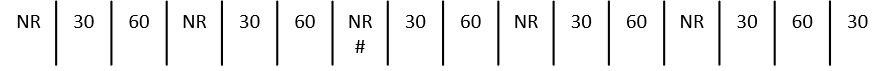

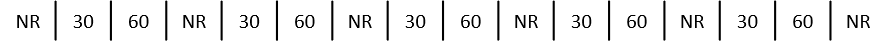

**mPFC**

pGSK3β-Ser9 – 46 kDa

Total GSK3β – 46 kDa

pCRMP2-Thr514 – 60-80 kDa

Total CRMP2 – 60-80 kDa

GAPDH – 37 kDa

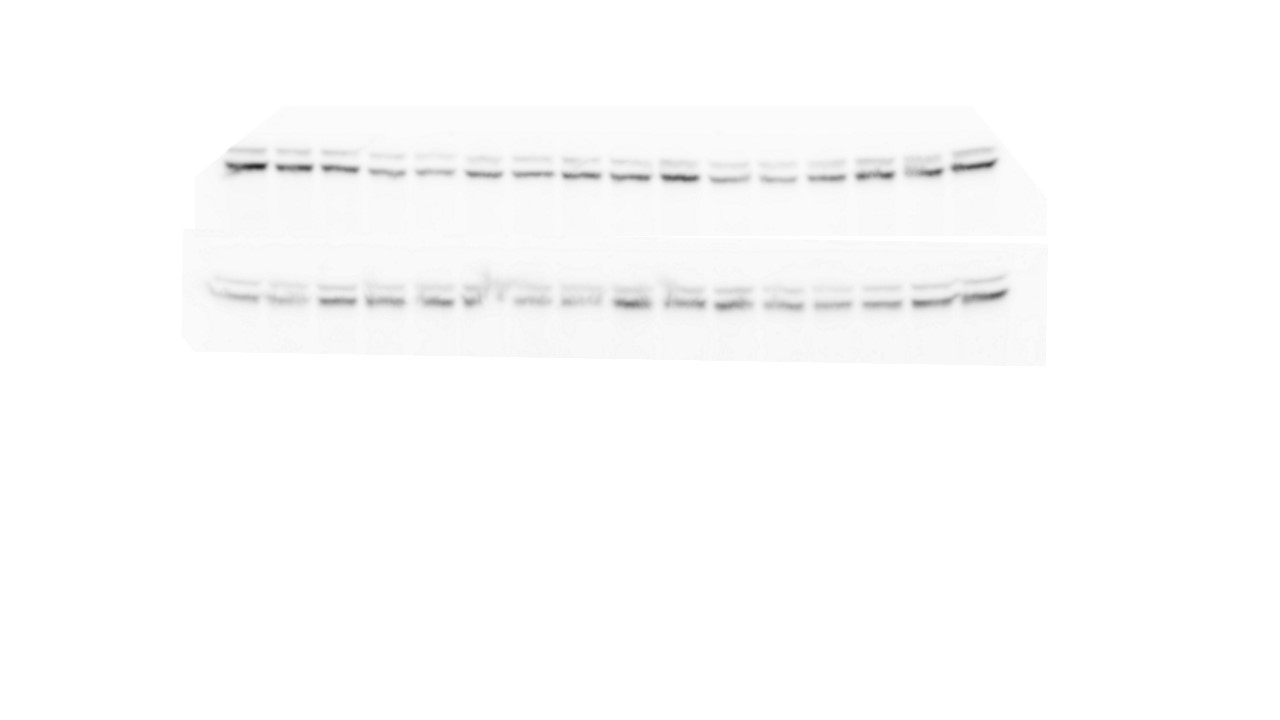

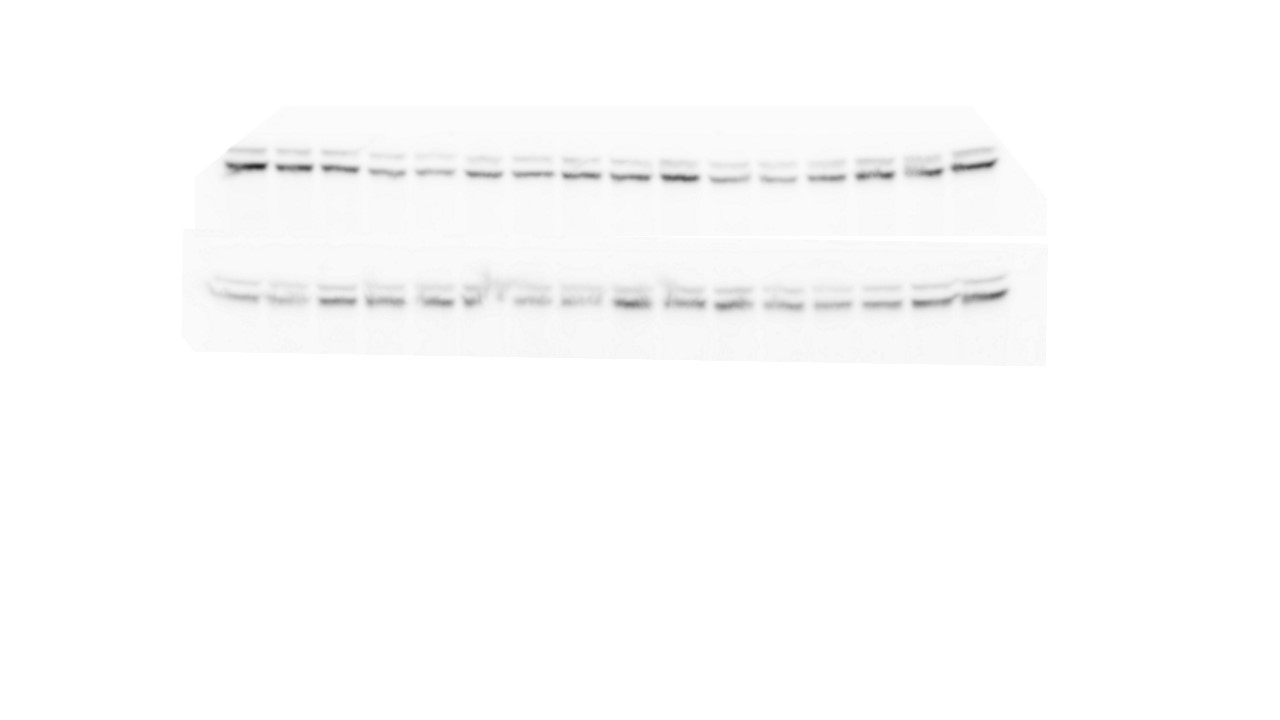

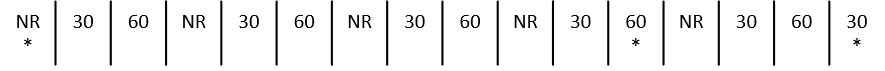

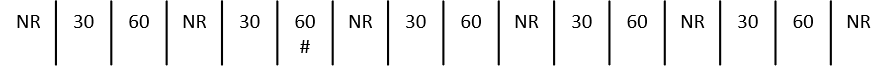

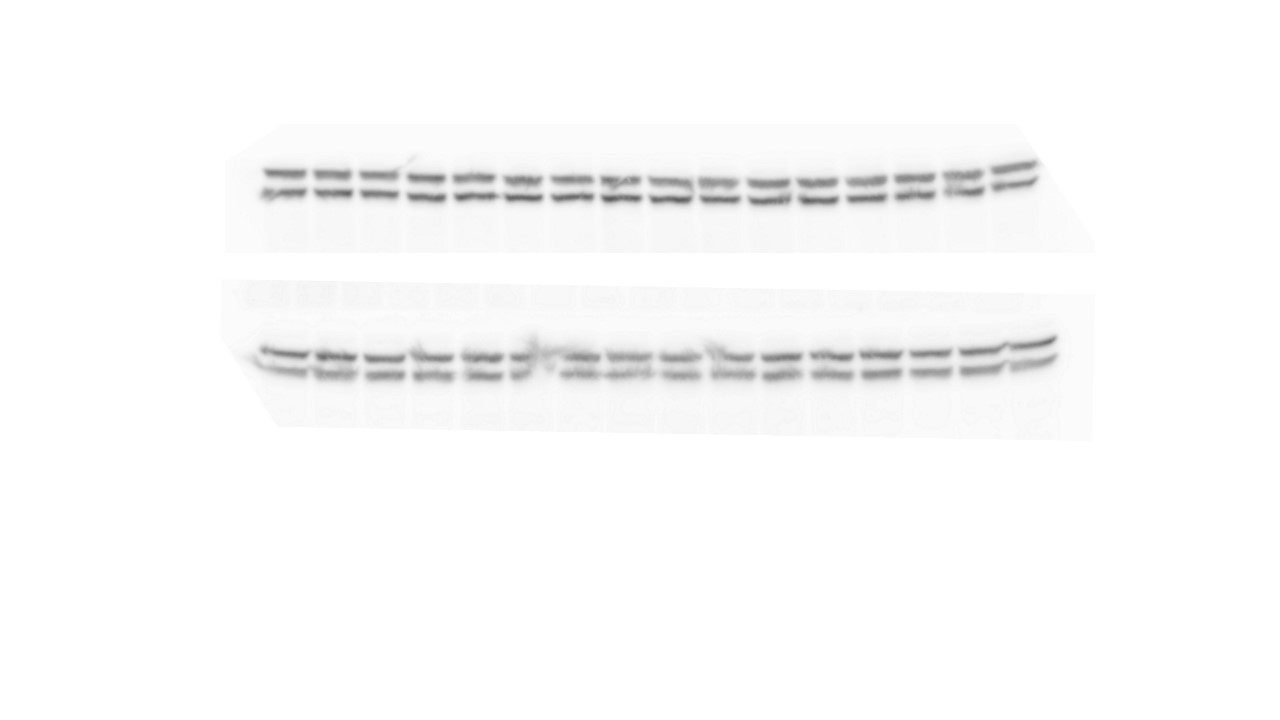

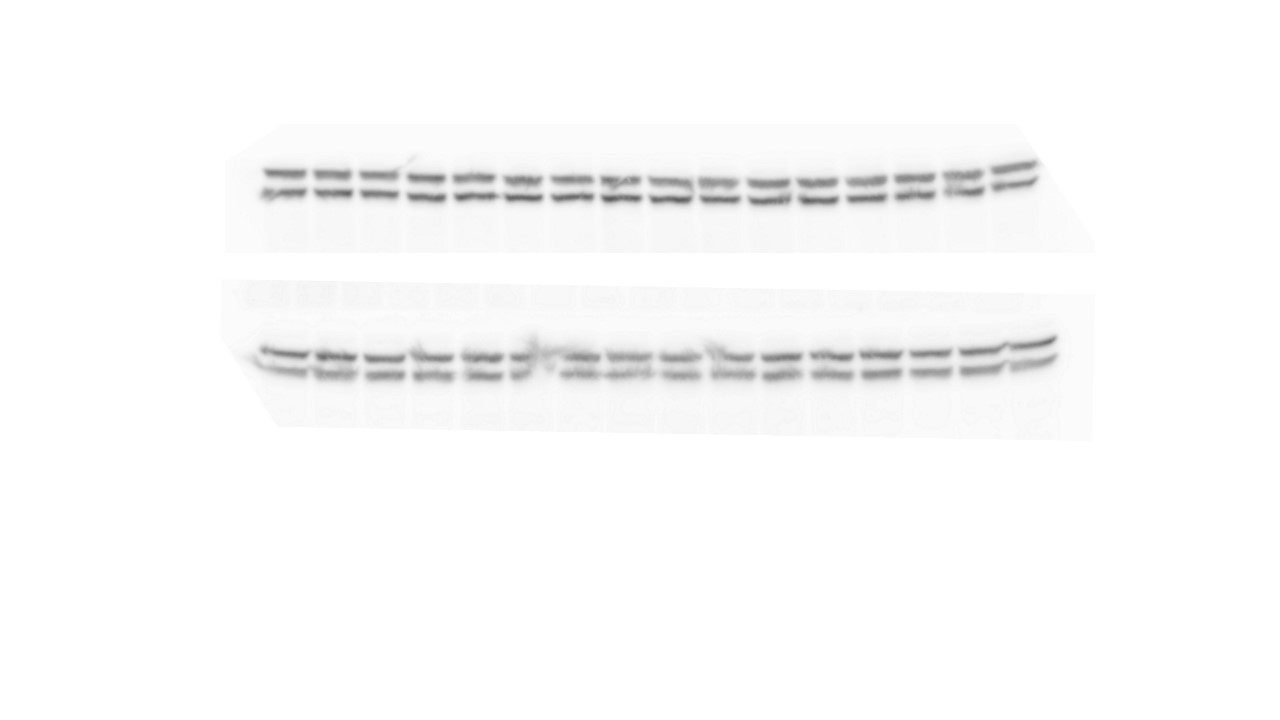

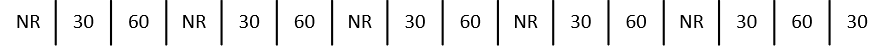

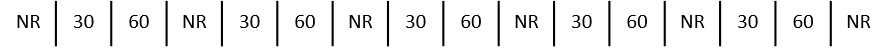

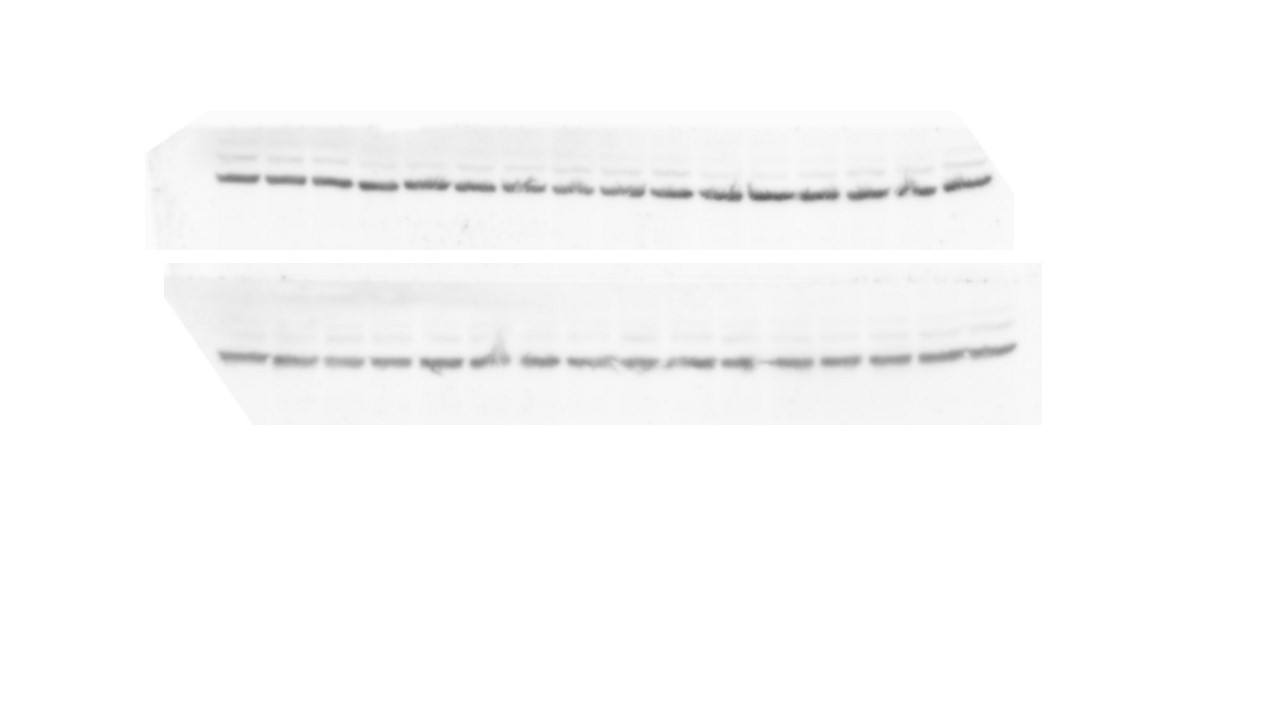

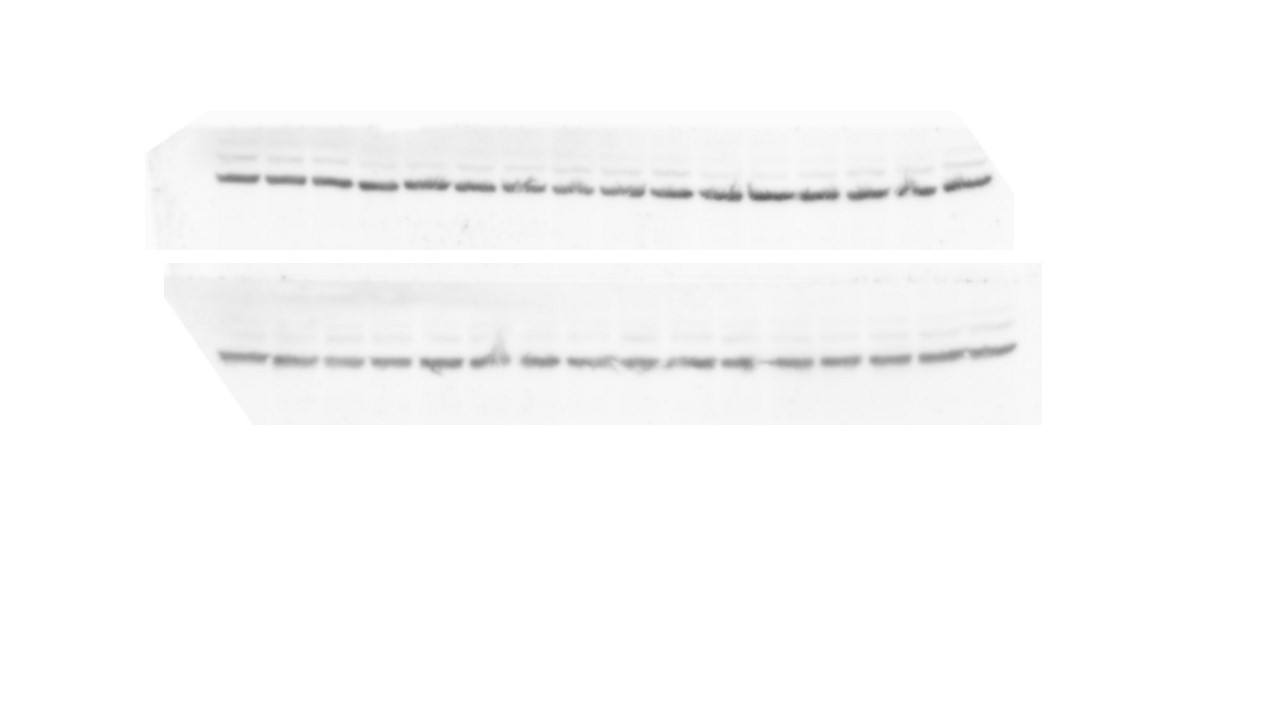

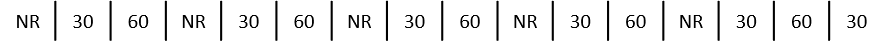

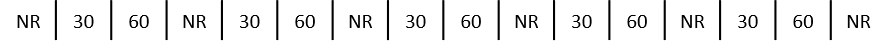

pERK1/2-Thr202/Tyr204 – 44/42 kDa

Total ERK1/2 – 44/21 kDa

**mPFC**

GAPDH – 37 kDa

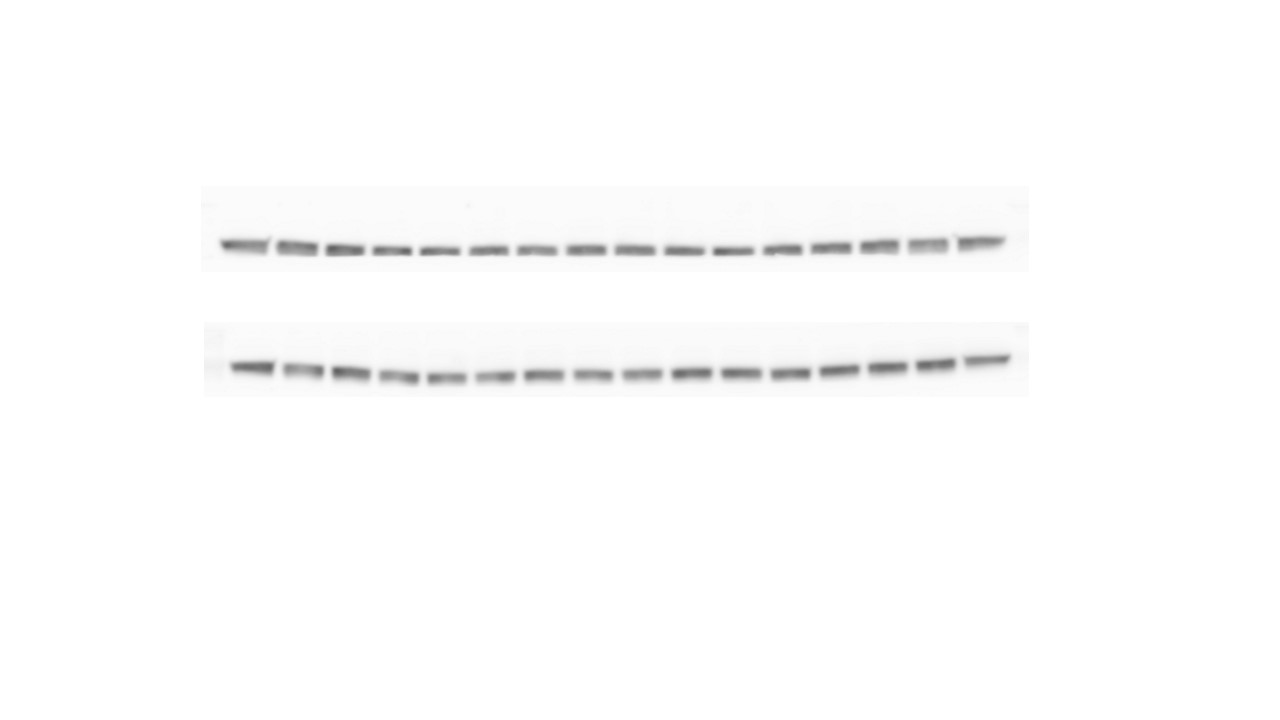

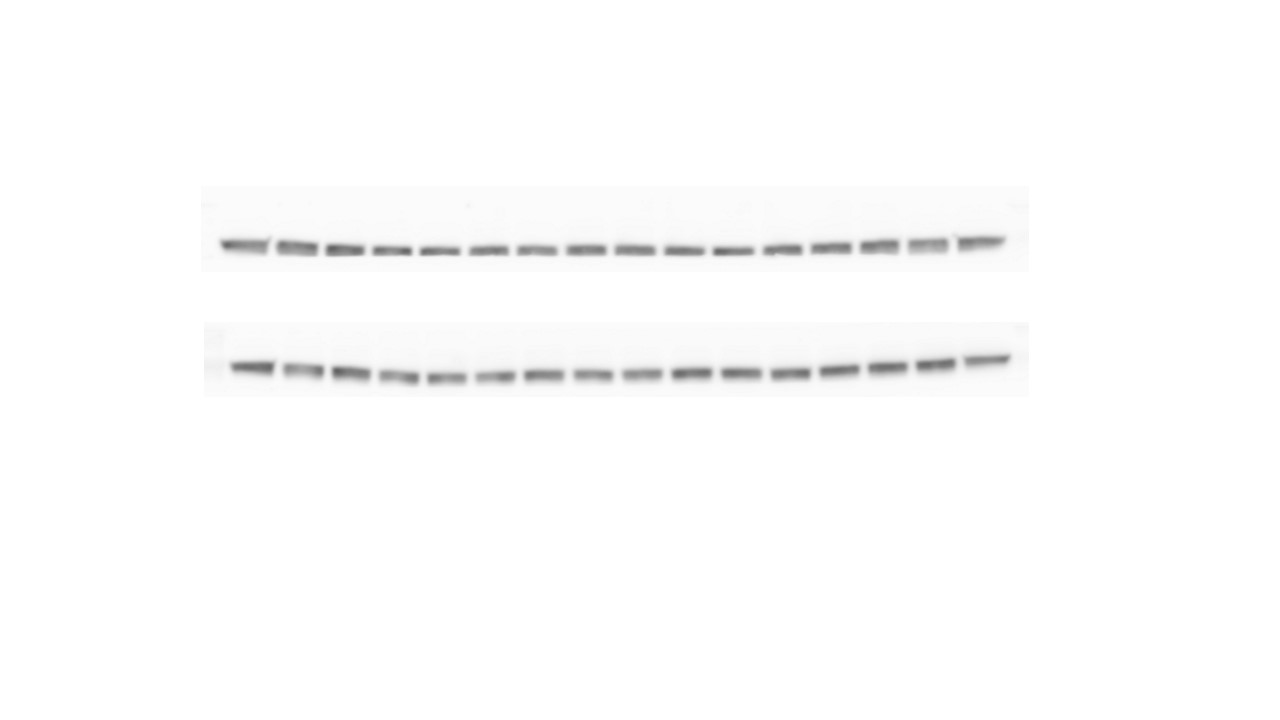

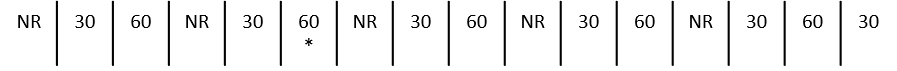

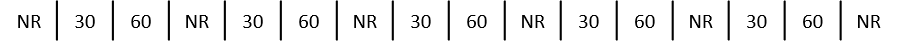

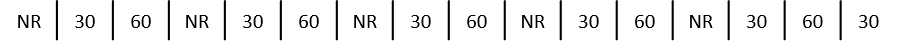

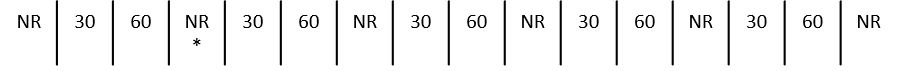

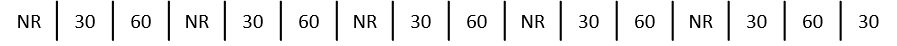

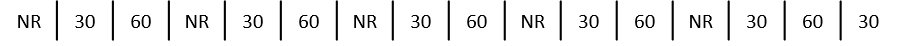

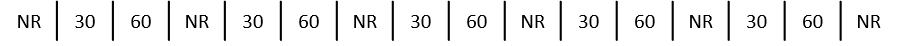

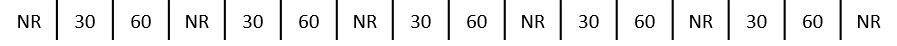

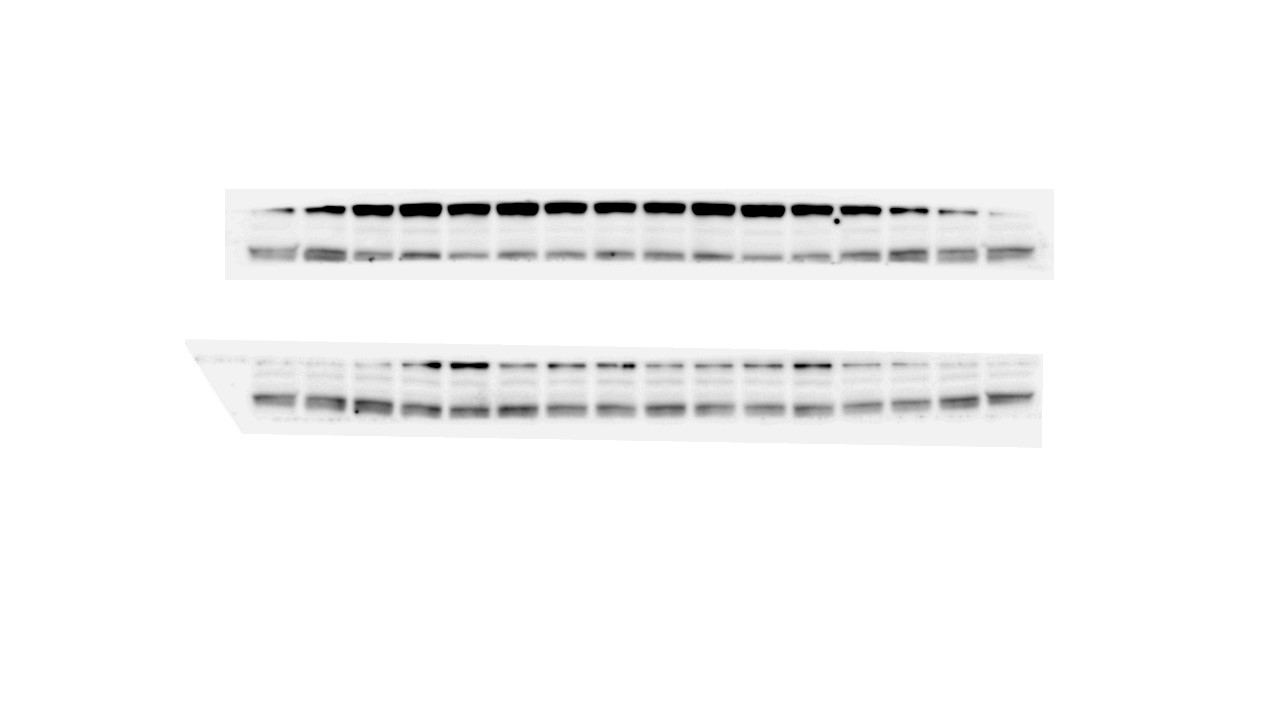

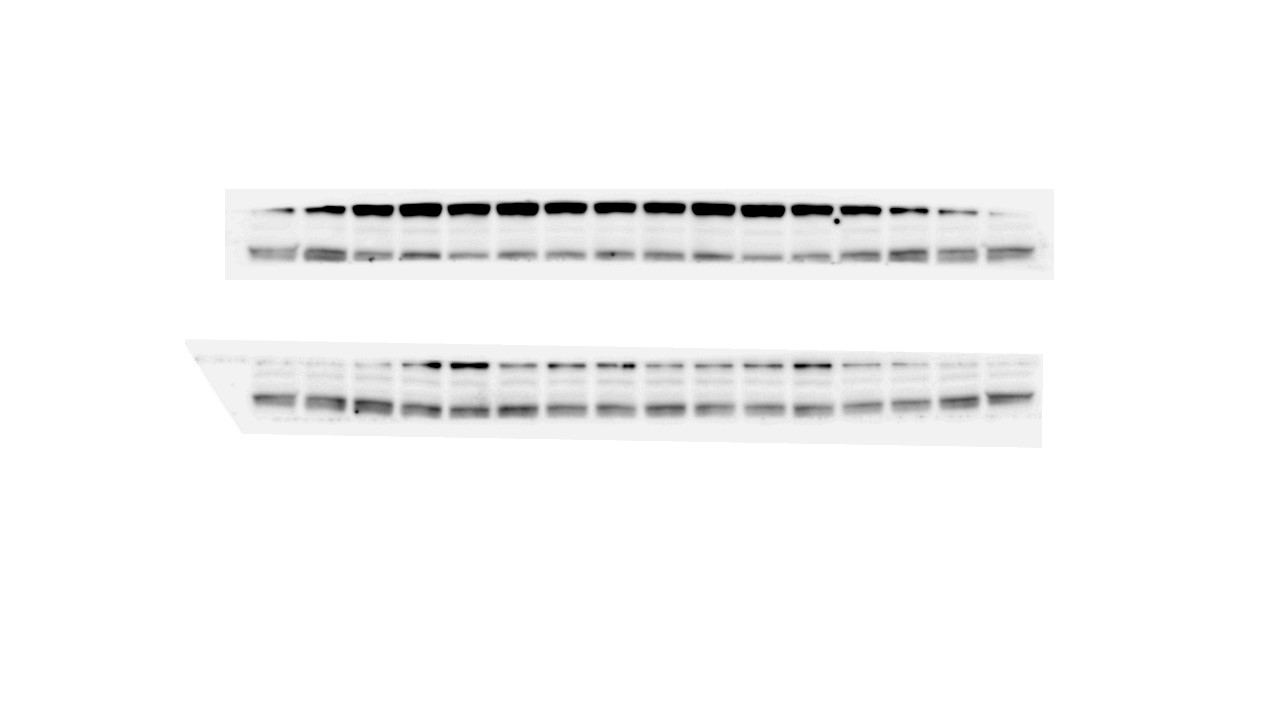

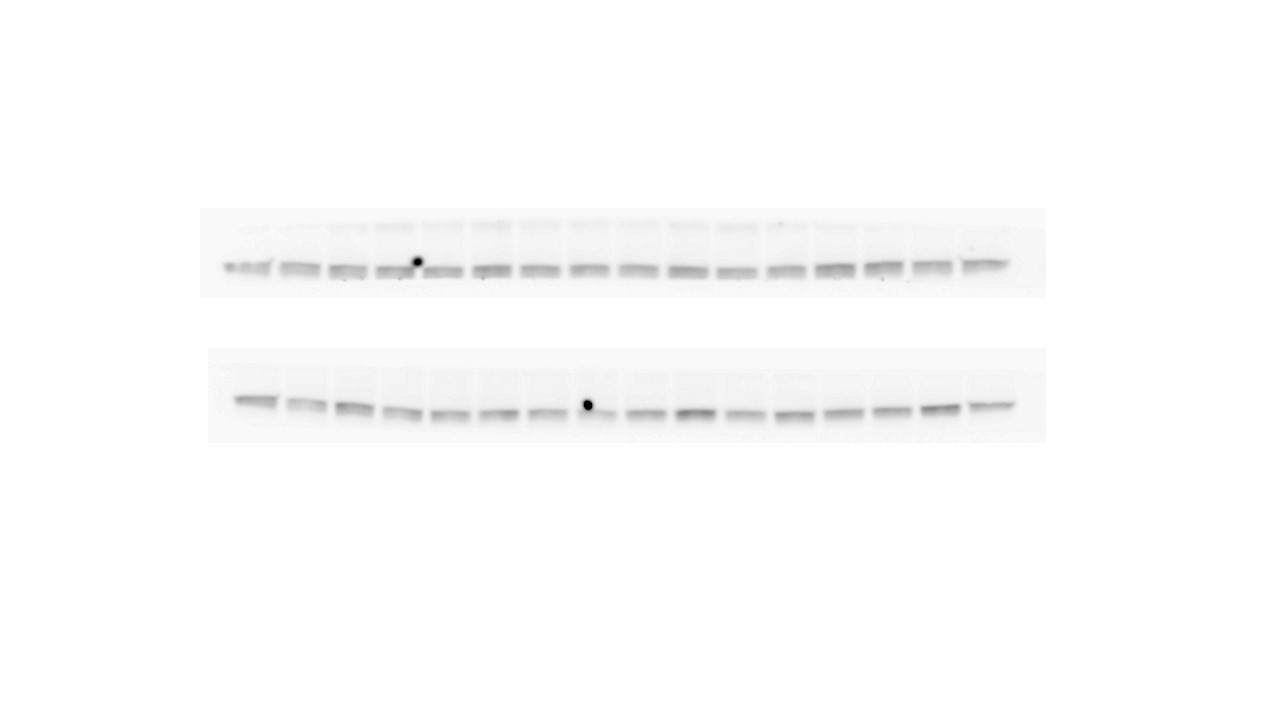

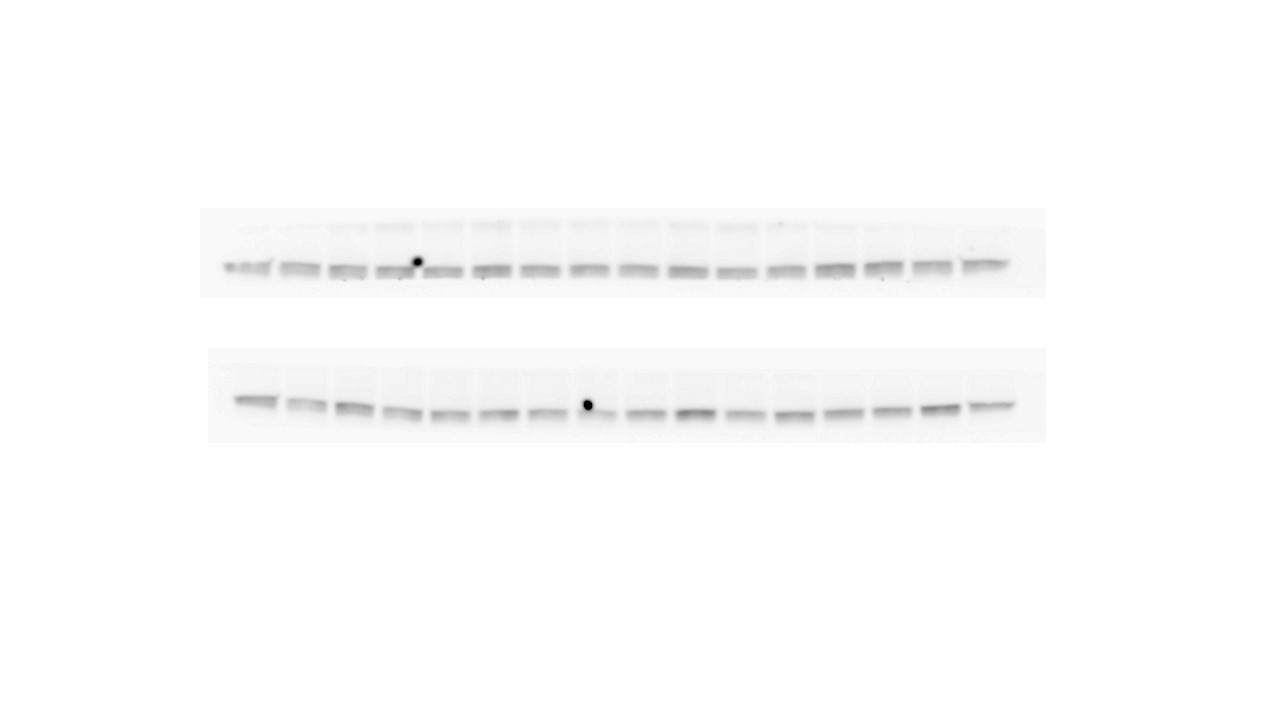

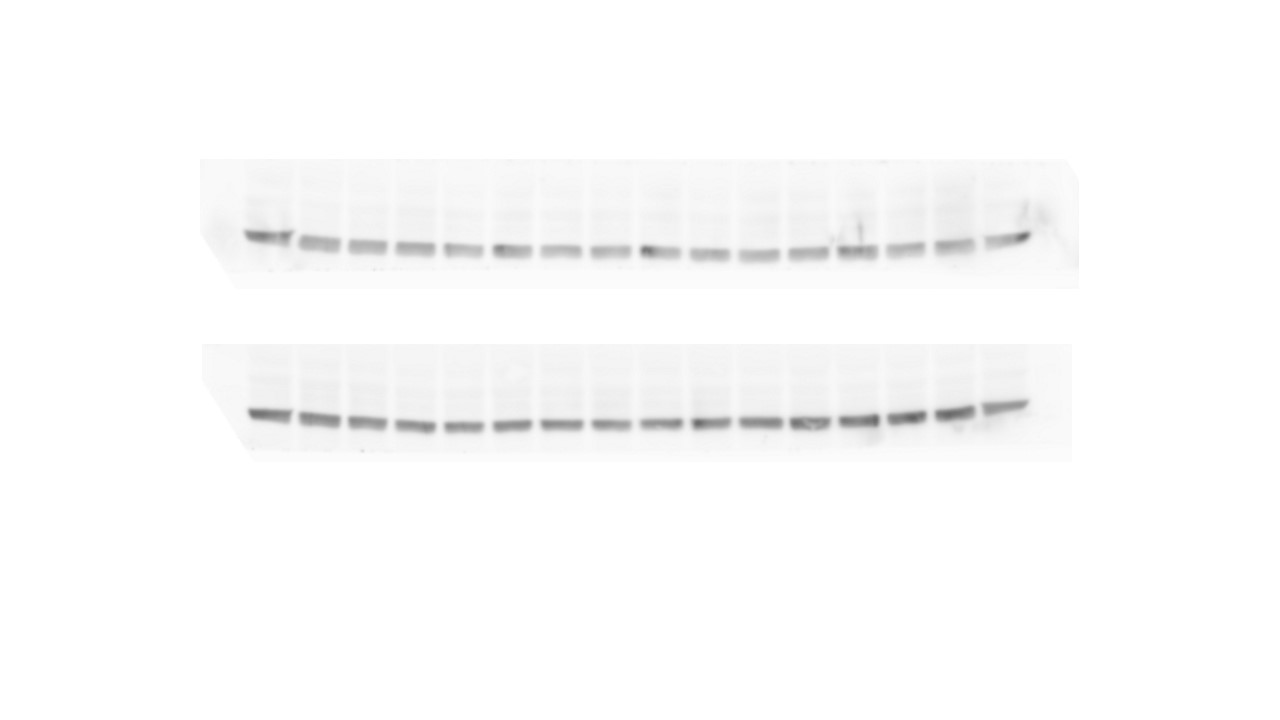

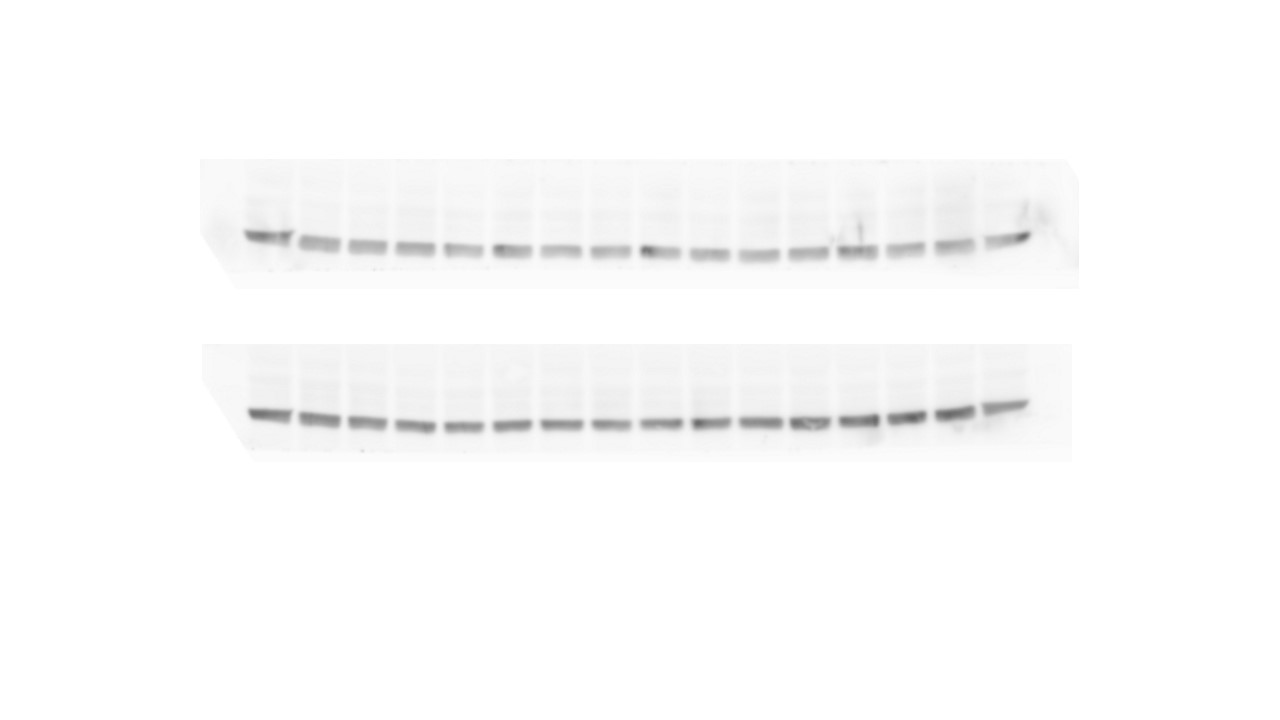

**NAc**

pAKT-Thr308 – 60 kDa

Total AKT - 60 kDa

GAPDH – 37 kDa

**B**

pAKT-Ser473 – 60 kDa

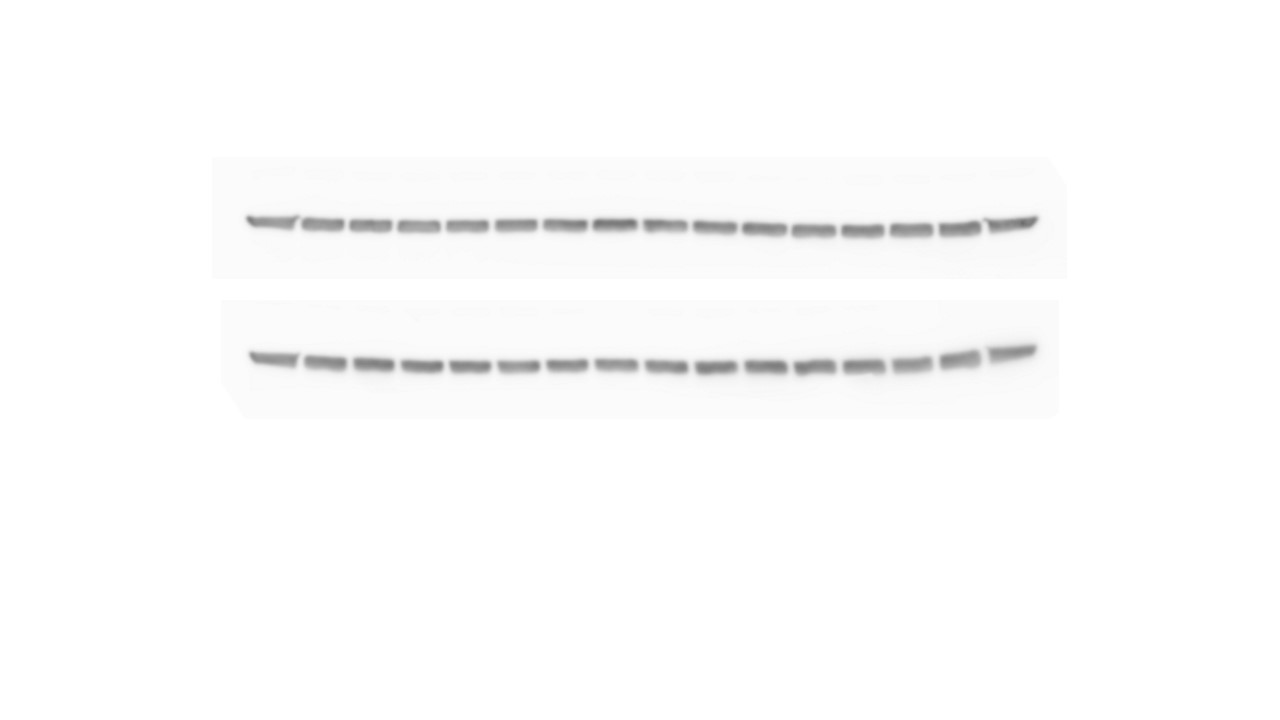

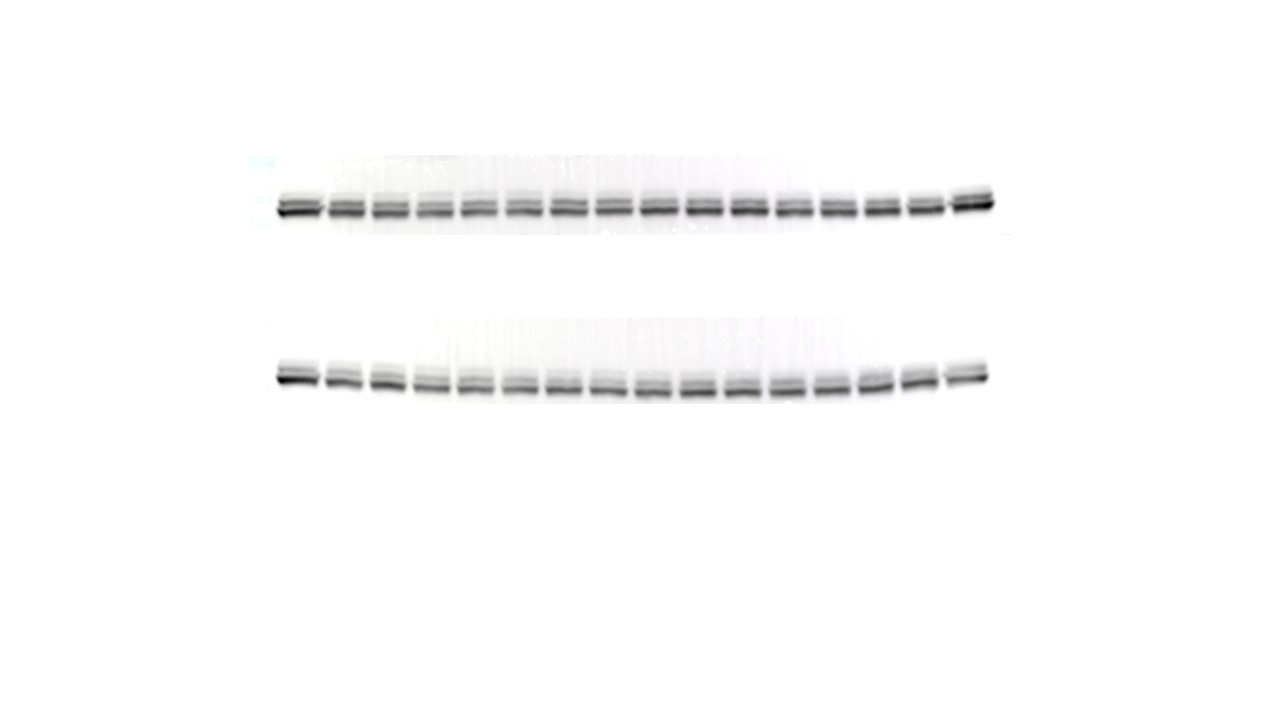

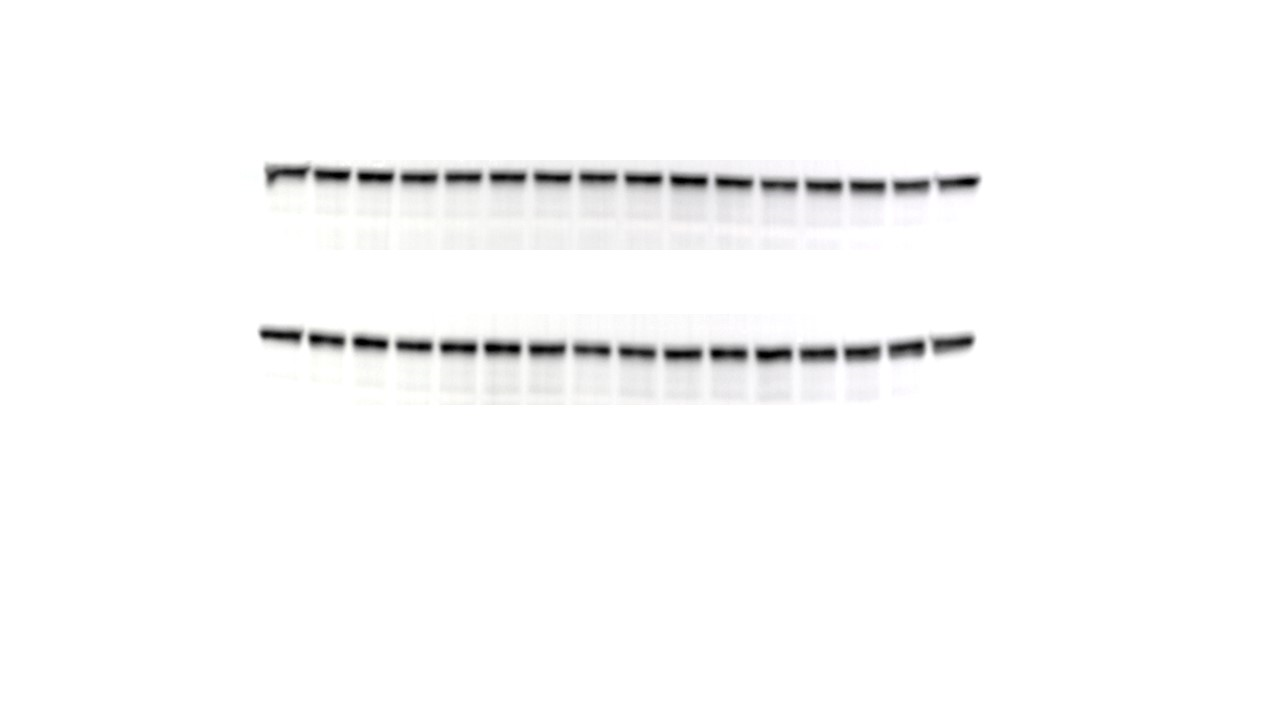

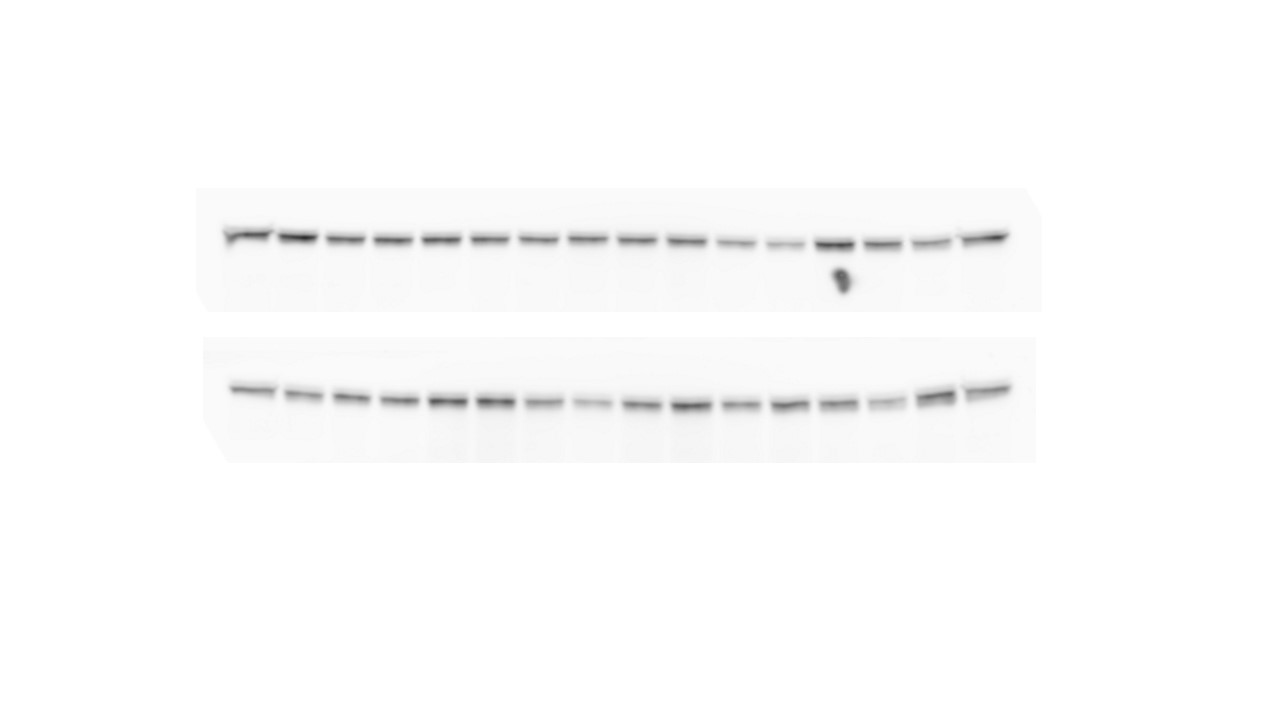

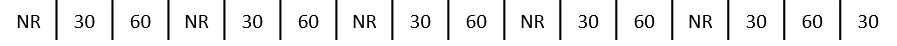

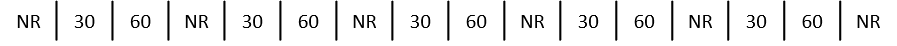

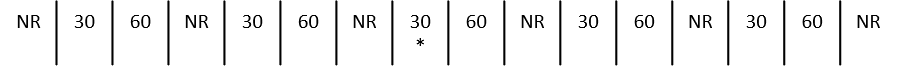

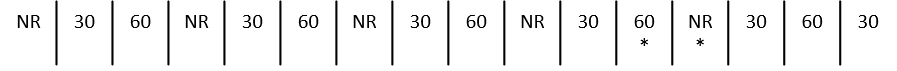

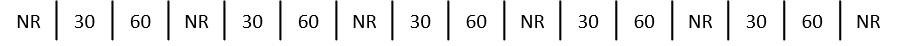

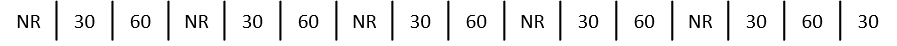

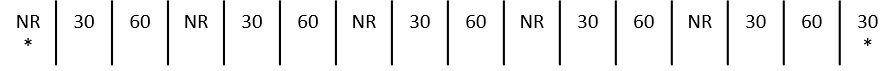

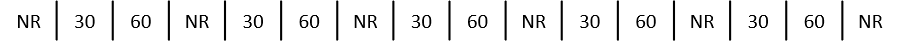

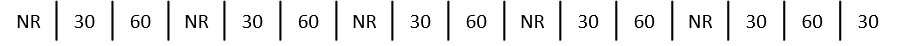

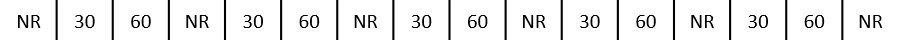

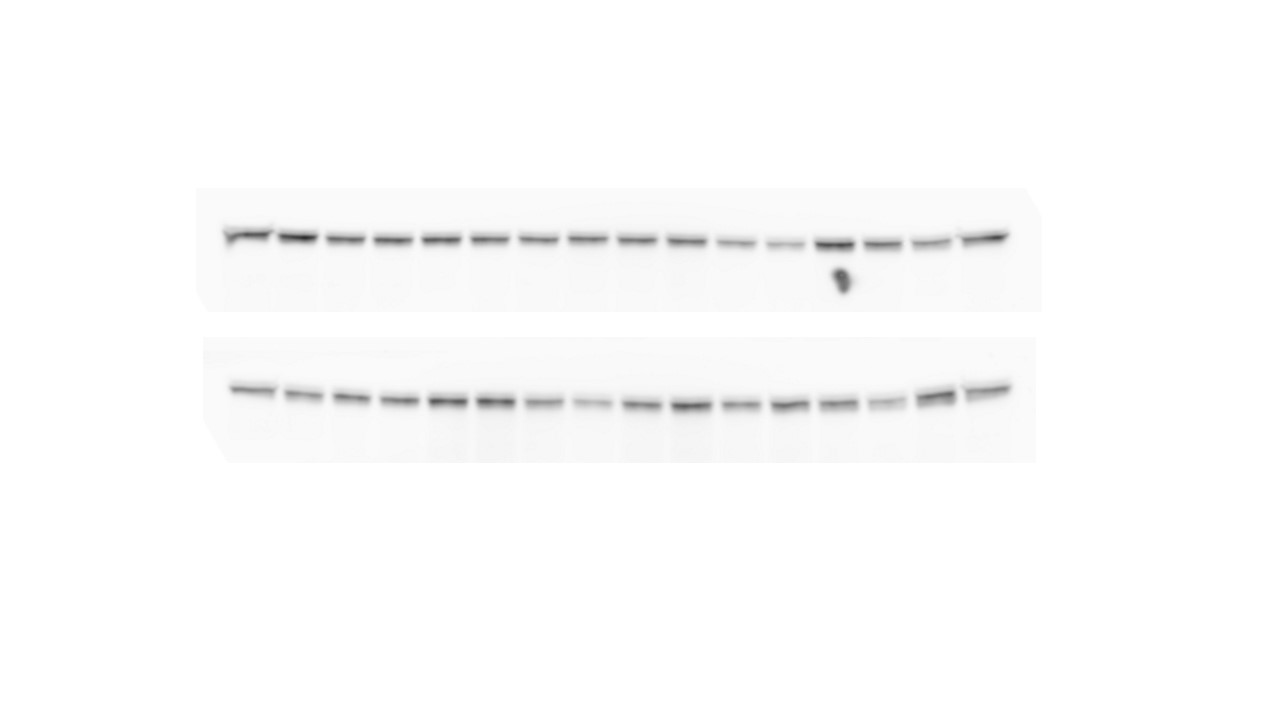

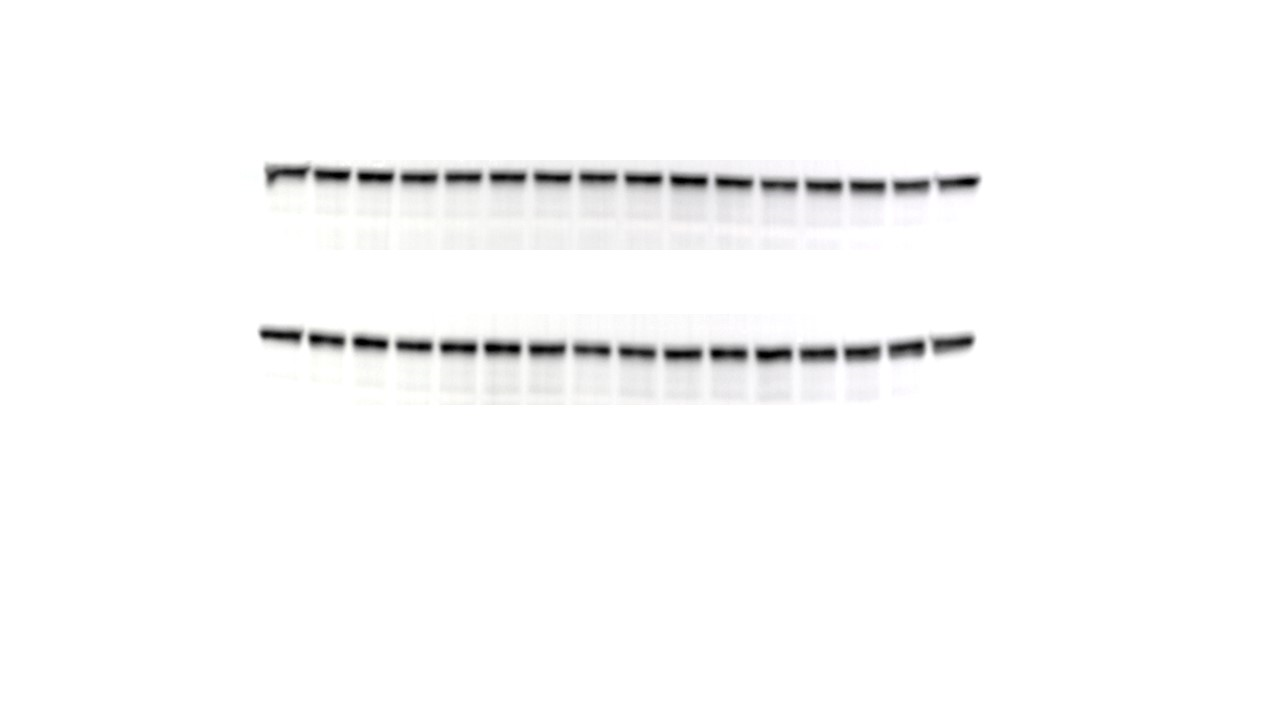

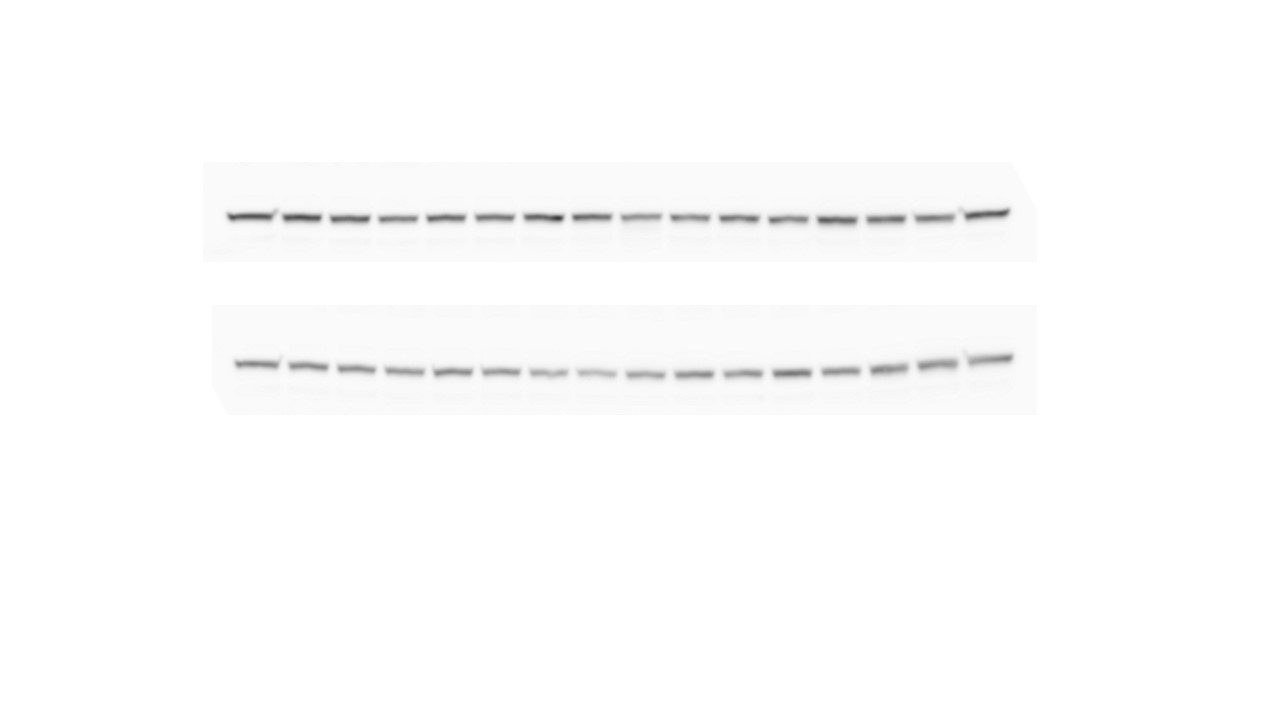

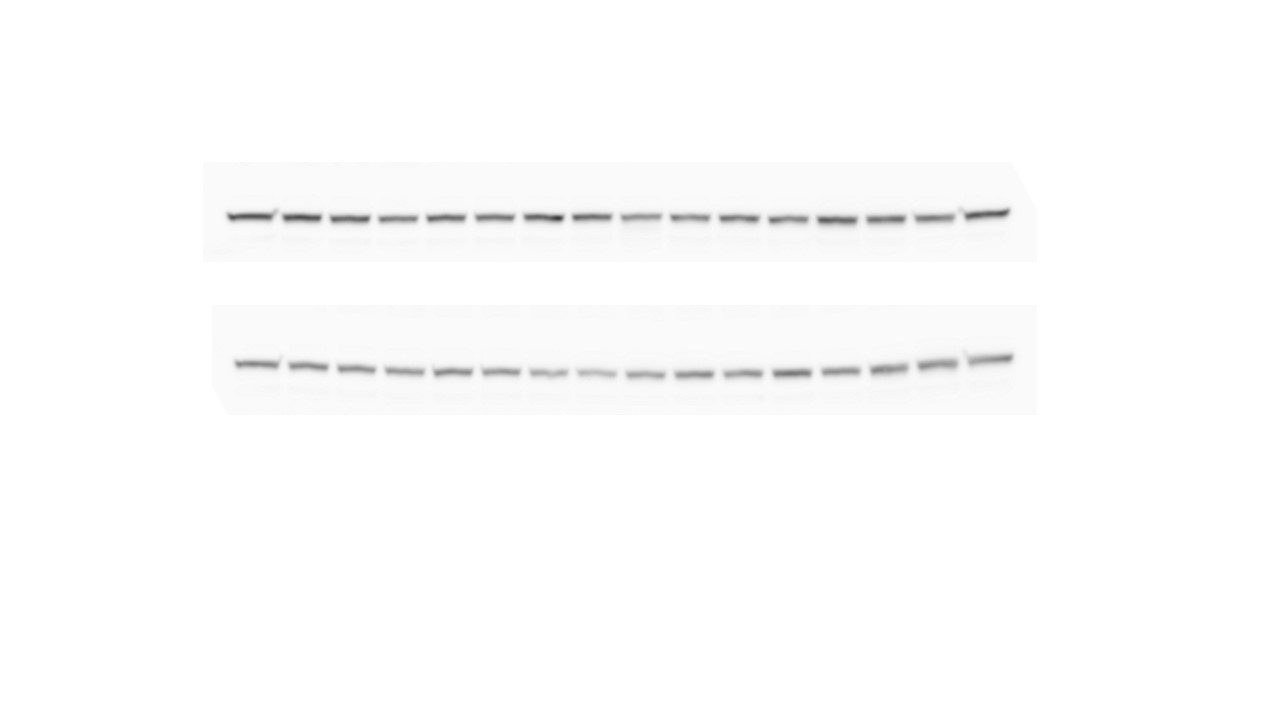

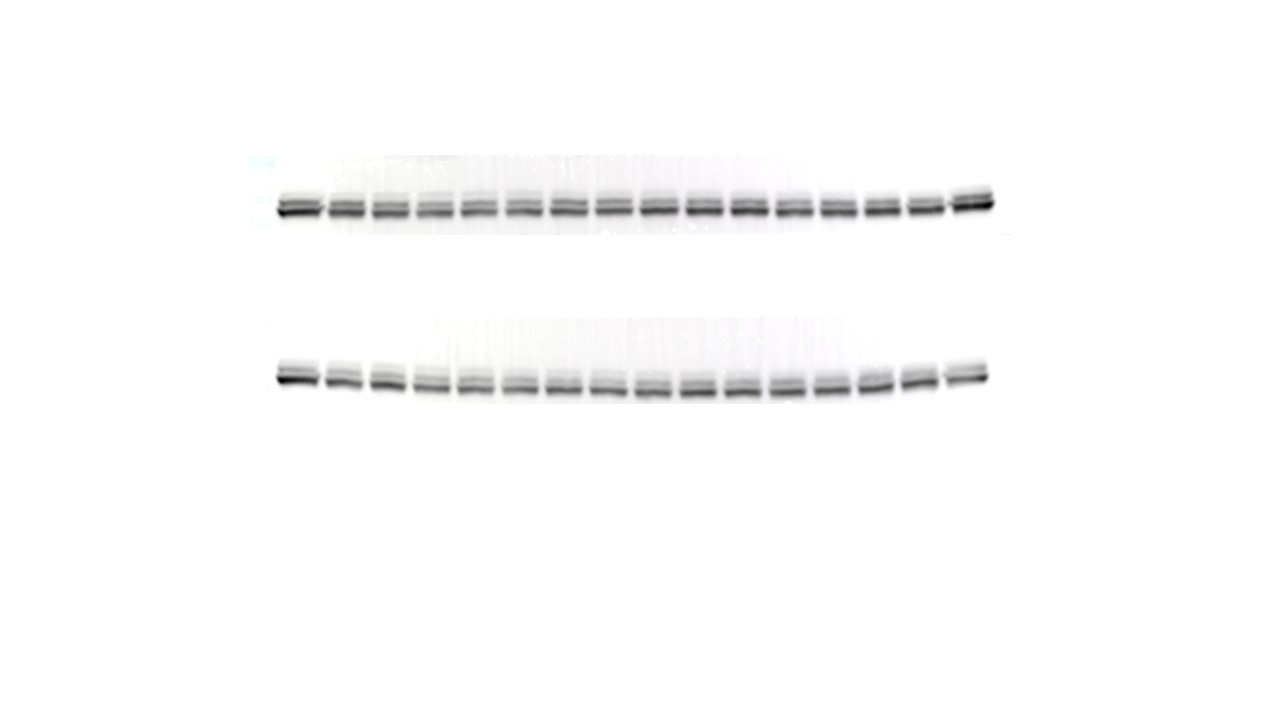

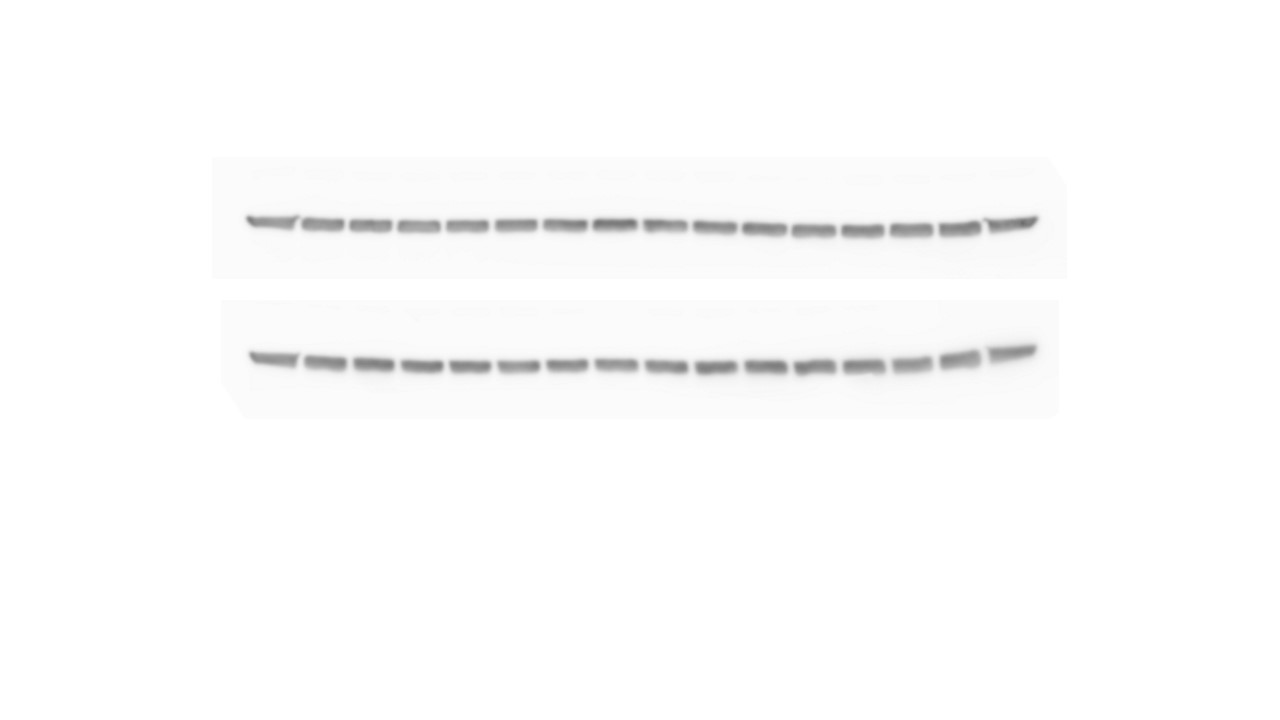

**NAc**

pGSK3β-Ser9 – 46 kDa

Total GSK3β – 46 kDa

pCRMP2-Thr514 – 60-80 kDa

Total CRMP2 – 60-80 kDa

GAPDH – 37 kDa

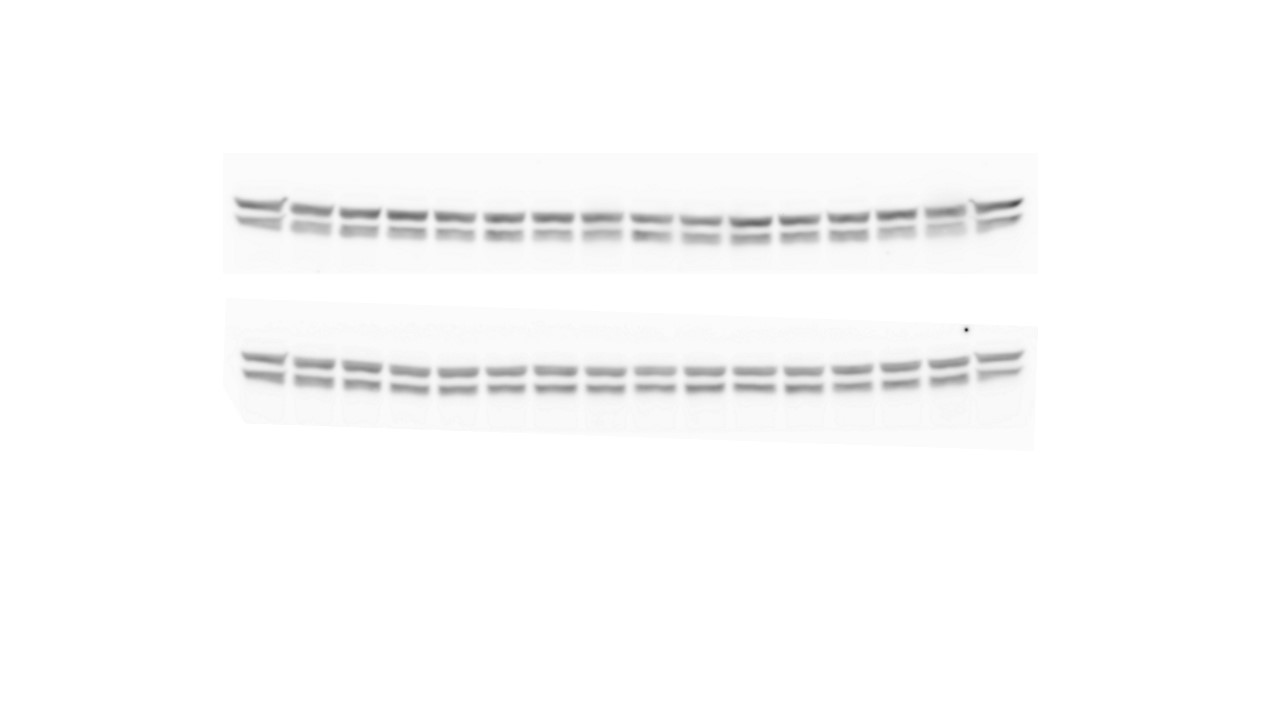

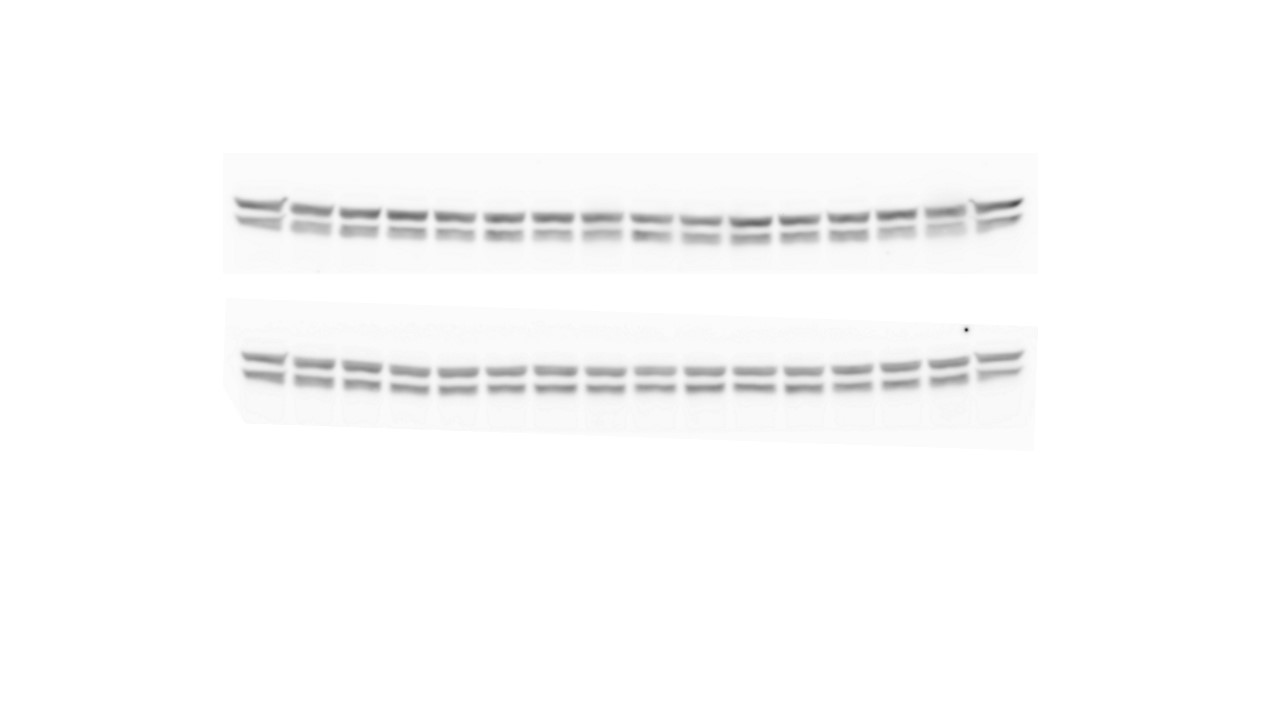

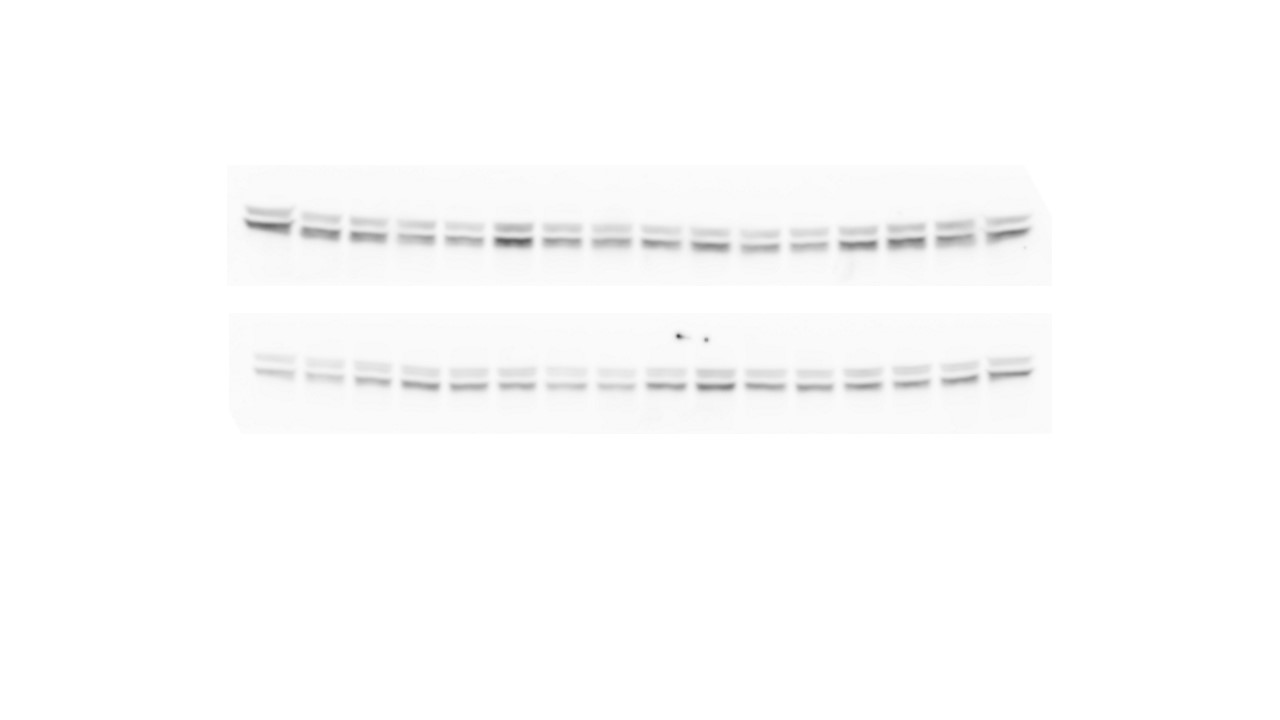

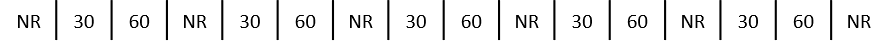

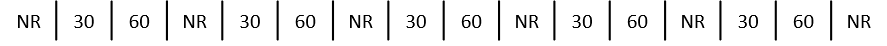

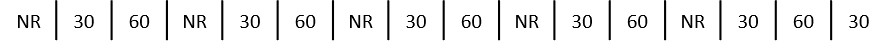

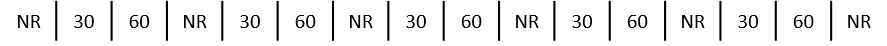

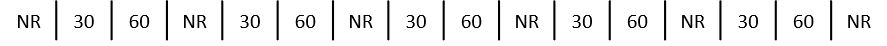

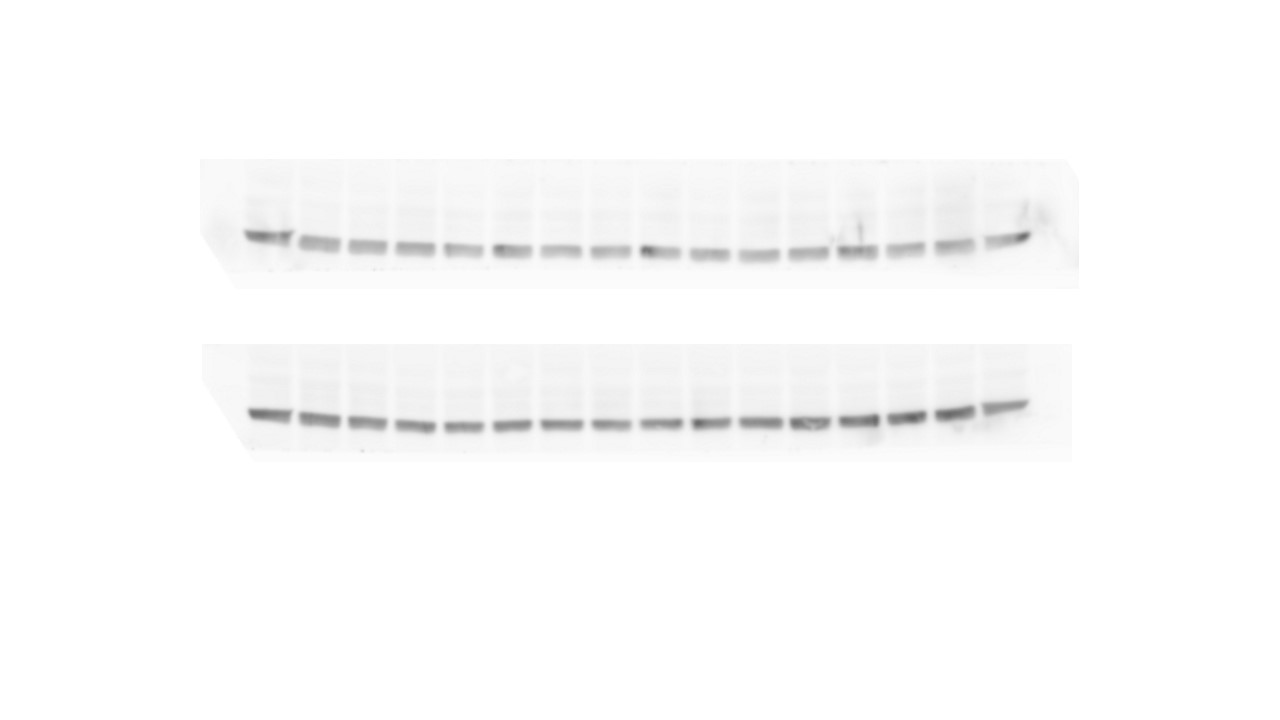

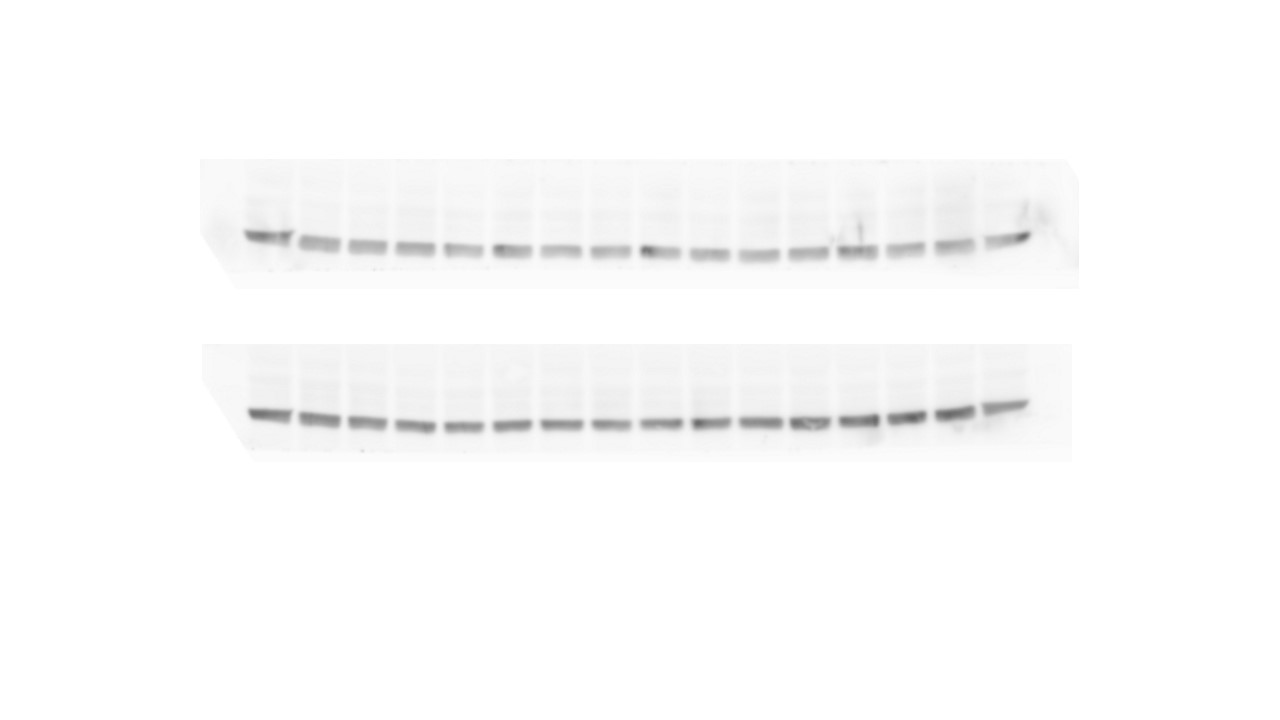

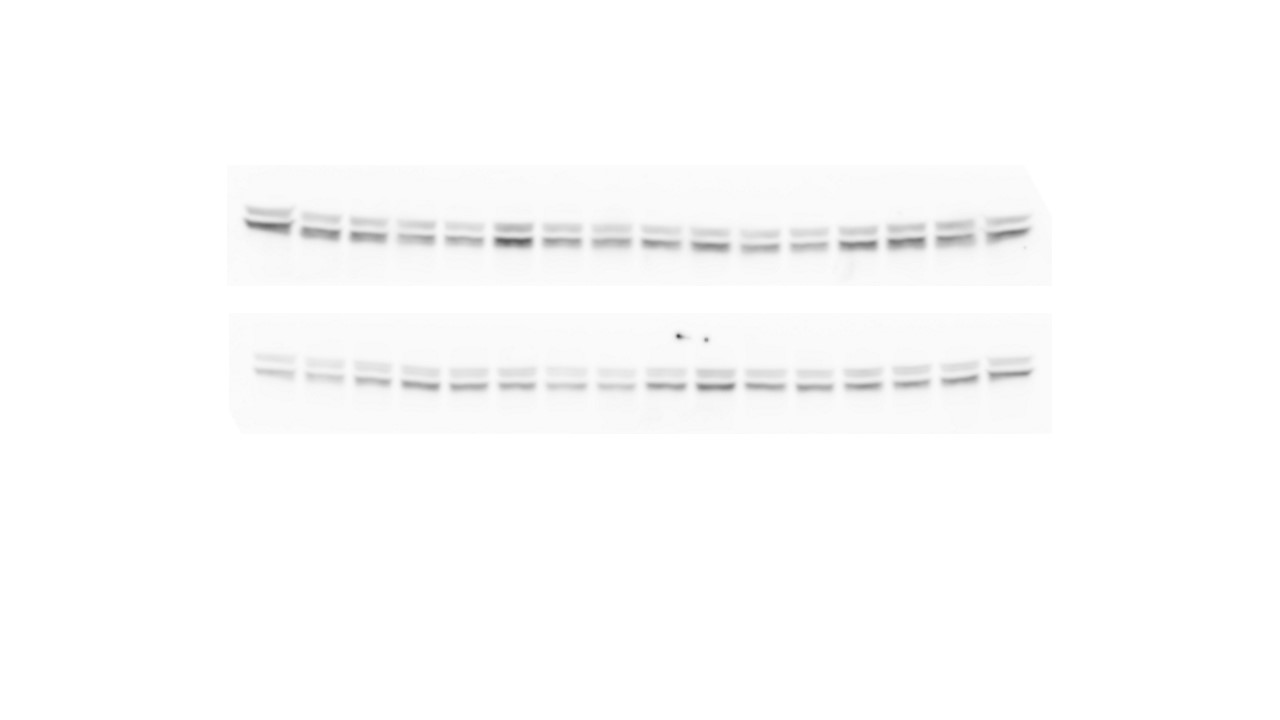

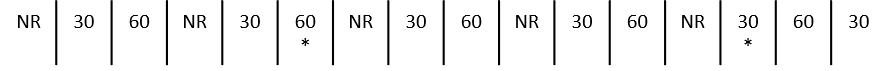

pERK1/2-Thr202/Tyr204 – 44/42 kDa

Total ERK1/2 – 44/21 kDa

**NAc**

GAPDH – 37 kDa

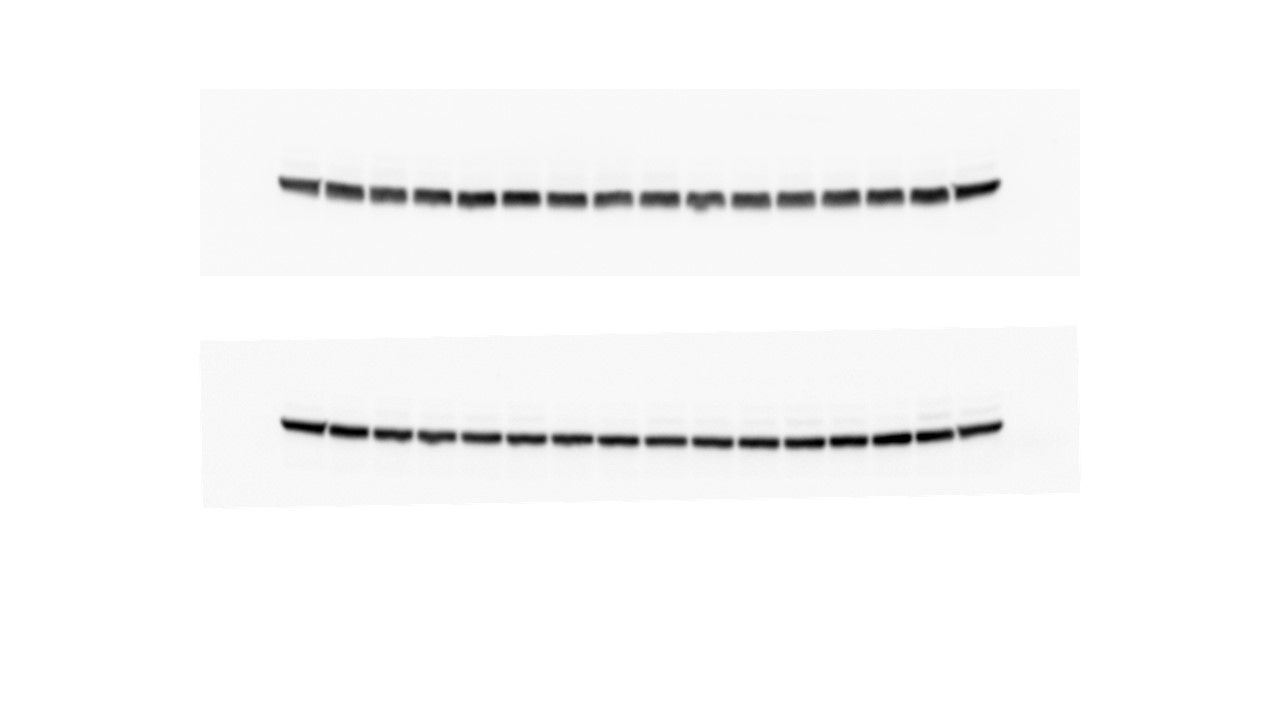

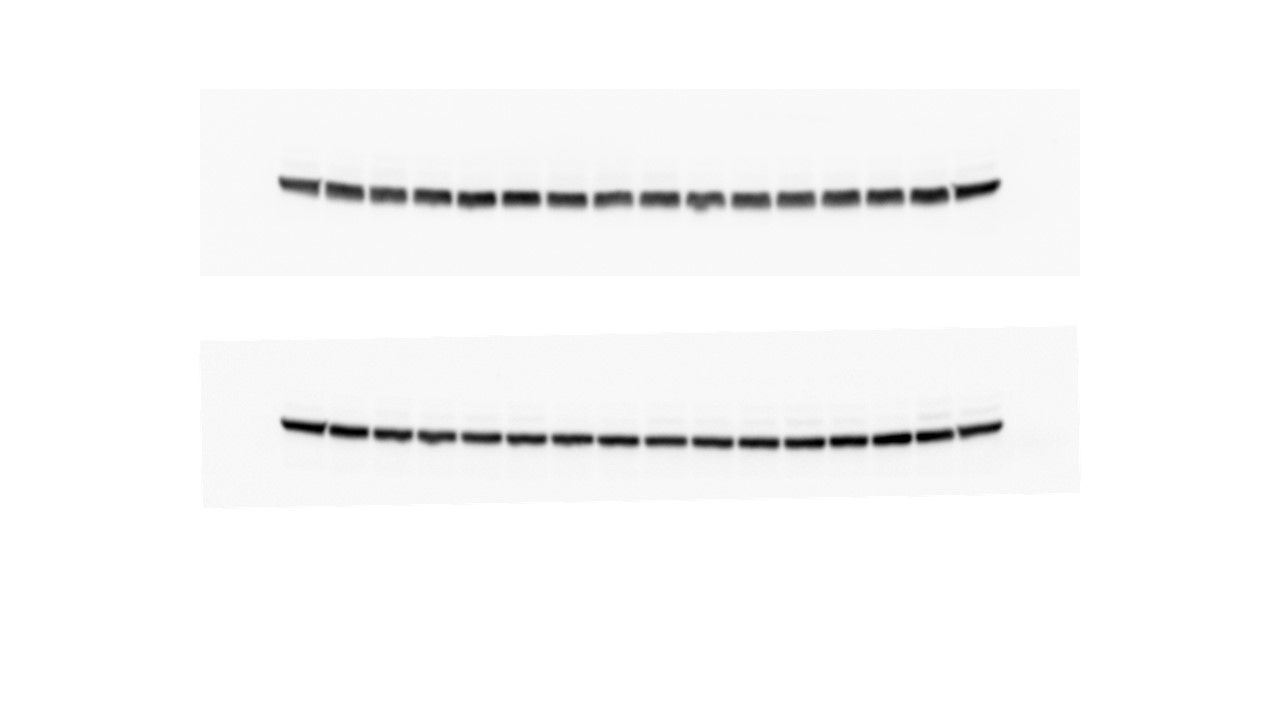

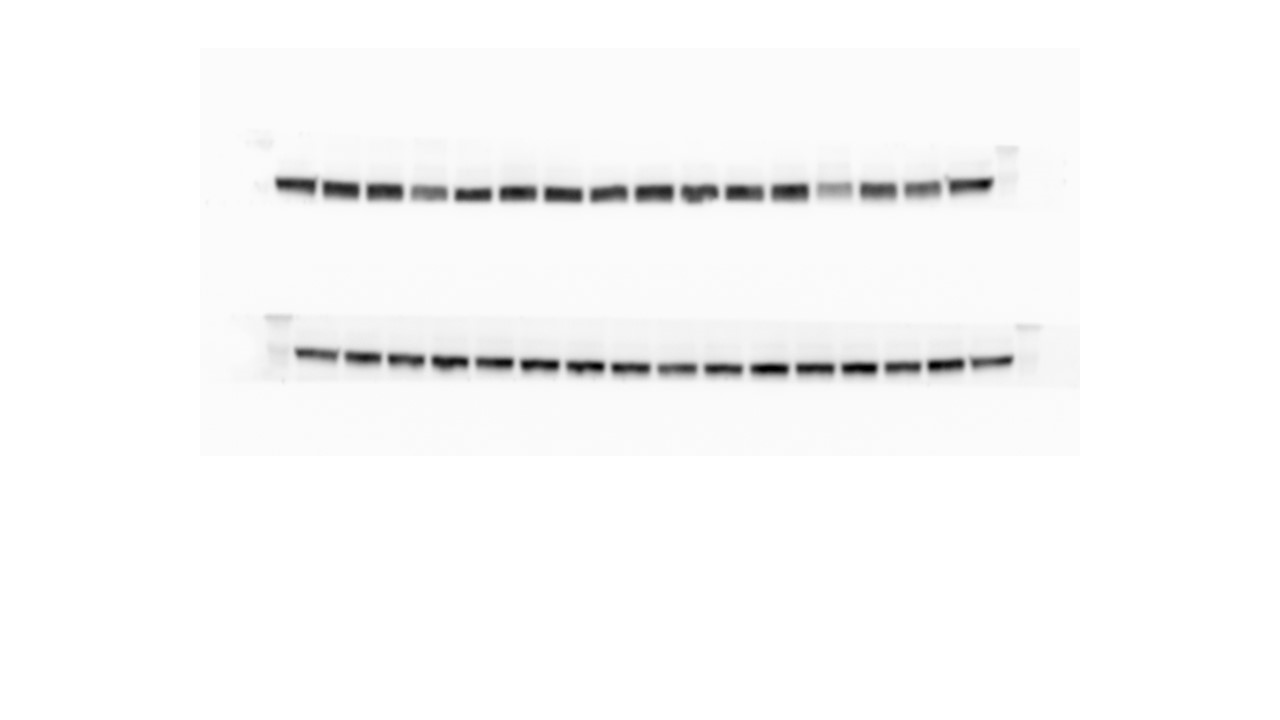

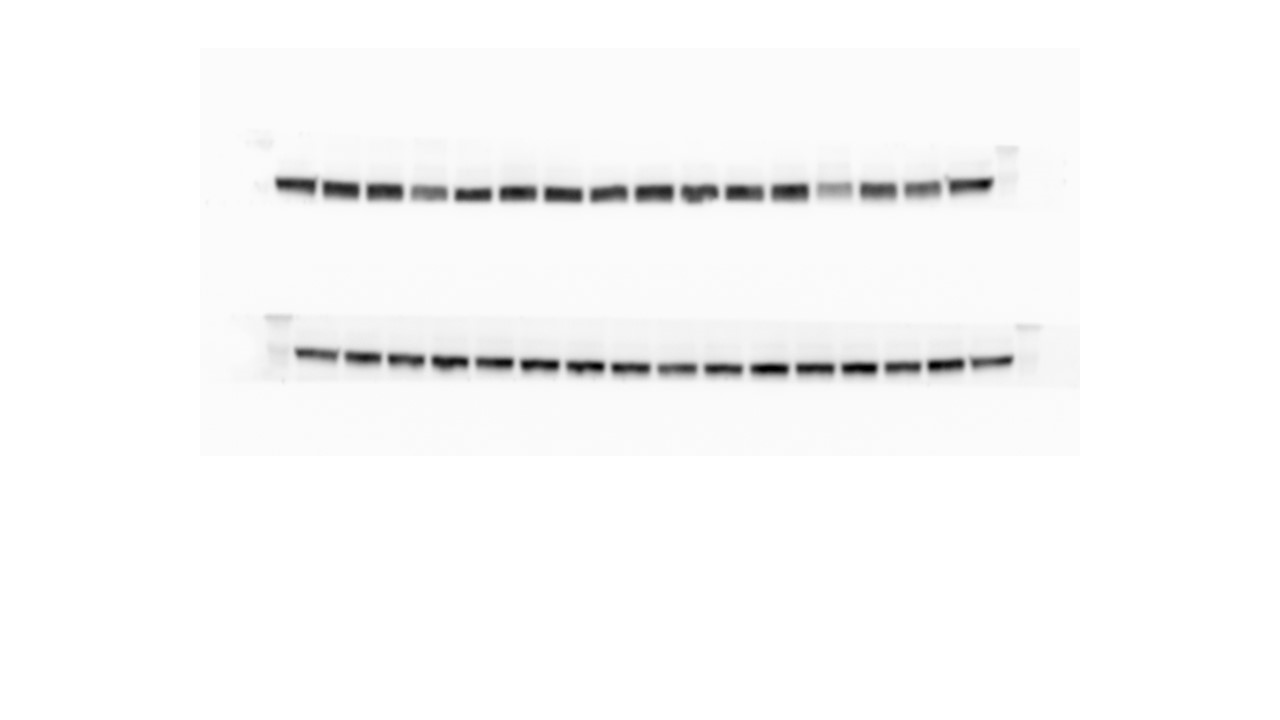

**Amygdala**

pAKT-Thr308 – 60 kDa

pAKT-Ser473 – 60 kDa

Total AKT - 60 kDa

GAPDH – 37 kDa

**C**

**Amygdala**

pGSK3β-Ser9 – 46 kDa

Total GSK3β – 46 kDa

pCRMP2-Thr514 – 60-80 kDa

Total CRMP2 – 60-80 kDa

GAPDH – 37 kDa

Total ERK1/2 – 44/21 kDa

**Amygdala**

GAPDH – 37 kDa

pERK1/2-Thr202/Tyr204 – 44/42 kDa

**Figure S1. Western-blot visualization of phospho-protein and total-protein levels in the mPFC (A), NAc (B), and amygdala (C) (Experiment 1).** The tissues were collected 30 min (30) or 60 min (60) after alcohol memory retrieval, or without a prior retrieval procedure (NR). Phospho-protein levels of AKT (Thr308, Ser473), GSK3β (Ser9), CRMP2 (Thr514) and ERK1/2 (Thr202/Tyr204) were normalized to the total protein immunoreactivity. Total protein levels of CRMP2 were normalized to GAPDH. AKT and ERK antibodies were blotted on the same set of membranes, and therefore share the GAPDH levels. GSK3β and CRMP2 were blotted on a different set of membranes. * Samples that were identified as outliers to their experimental group (z>2) and were therefore removed from the analysis. # Distorted bands that were removed from the analysis.

pERK1/2-Thr202/Tyr204 – 44/42 kDa

Total ERK1/2 – 44/21 kDa

GAPDH – 37 kDa

***Supplementary Figure 2***

**Figure S2. Western-blot visualization of phospho-protein and total-protein levels in the amygdala (Experiment 4).** The tissues were collected two hours after alcohol memory retrieval (R), or without a prior retrieval procedure (NR). Phospho-protein levels of ERK1/2 (Thr202/Tyr204) were normalized to the total protein immunoreactivity.

pERK1/2-Thr202/Tyr204 – 44/42 kDa

Total ERK1/2 – 44/21 kDa

GAPDH – 37 kDa

pS6 – 32 kDa

Total S6 – 32 kDa

***Supplementary Figure 3***

**Figure S3. Western blot visualization of phospho-protein and total-protein levels in the amygdala (Experiment 5).** The tissues were collected 90 min after injection of vehicle (V) or the MERK1/2 inhibitor SL-327 (50 mg/kg. i.p.) (SL). Phospho-protein levels of ERK1/2 (Thr202/Tyr204) and S6 (Ser235/236) were normalized to the total protein immunoreactivity. * Sample that was identified as an outlier to its experimental group (z>2) and was therefore removed from the analysis.
